# Noninvasive optical detection of Granzyme B from natural killer cells using enzyme-activated fluorogenic probes

**DOI:** 10.1101/2019.12.16.875070

**Authors:** Tomasz Janiszewski, Sonia Kołt, Dion Kaiserman, Scott Snipas, Shuang Li, Julita Kulbacka, Jolanta Saczko, Niels Bovenschen, Guy Salvesen, Marcin Drąg, Phillip I. Bird, Paulina Kasperkiewicz

## Abstract

Despite many studies on the cytotoxic protease granzyme B, key aspects of its function remain unexplored due to the lack of selective probes for its activity. In this study, we fully mapped the substrate preferences of GrB using a set of unnatural amino acids, demonstrating previously unknown GrB substrate preferences that we then used to design novel substrate-based inhibitors and a GrB-activatable activity-based probe. We showed that our GrB probes react poorly with caspases, making them ideal for the in-depth analysis of GrB localization and function in cells. With our quenched fluorescence substrate, we determined GrB within the cytotoxic granules of human YT cells. When used as cytotoxic effectors, YT cells loaded with the GrB attack MDA-MB-231 target cells, and active GrB influences its target cell killing efficiency.

## Introduction

In the last two decades, significant advances in the understanding of natural killer (NK) cells have been made^1^. These cytotoxic lymphocytes are key effectors of innate immunity and are involved in viral infection responses as well as controlling several types of tumors. The activation of these cells is initiated by major compatibility complex (MHC) class I protein loss in compromised cells^1^. NK cell activation changes the balance between the activating and inhibiting receptors on cell surfaces. Activated cells rapidly and quickly secrete cytokines: tumor necrosis factor α (TNFα) and interferon γ (IFNγ), leading to subsequent stimulation of the immune system. Reciprocal interactions with dendritic cells, macrophage T cells, and endothelial cells also enhance the immune system response. To prevent autoimmune damage, the NK cell inhibitory receptors recognize MHC class I proteins and protect healthy host cells^2^.

Among the various weapons of NK cells and cytotoxic T lymphocytes (CTLs), the most important, located in the cytotoxic granules, are perforin and granzymes (**Gran**ule associated en**zymes, Grs**). Grs are the family of homologous serine proteases and the five different human granzymes (A/B/H/K/M); the most studied and abundant are granzymes A and B^3, 4^. These enzymes are not only stored within the cytotoxic granules of immune killer cells but are also detected in primary breast carcinoma and in chondrocytes of articular cartilage^5^. Upon the activation of NK cells and binding to the target cell, the granule membrane fuses with the plasma membrane of the NK cell, and perforins (proteins that form pores in cell membranes) and granzymes are released from the cytotoxic effector cell into the intermembrane space^6, 7^. The mechanism of GrB entering the target cell through perforin-formed pores in the plasma membrane is still unclear^8^. GrB is transported into the target cell to carry out its effect. Within the cell, GrB initiates at least three distinct pathways of programmed cell death, namely, (1) the activation of caspase 3, which triggers apoptosis, (2) GrB caspase-3 substrate hydrolysis: the inhibitor of Caspase Activated DNAse (ICAD), Bid, (3) or the direct hydrolysis of lamin B.

The detection of individual granzymes with antibody-related techniques is challenging due to their structural similarities; for example, GrB antibody cross-reacts with GrA and GrH^3^. For this reason, the detailed analysis of individual Grs is challenging. To overcome this obstacle, chemical approaches for developing enzymatic inhibitors and activity-based probes (ABP) as efficient tools for granzyme exploration have been attempted. However, one problem is that the substrate specificity of GrB is similar to that of some caspases (casp-8)^9^, cleaving the same substrates at the same cleavage site (synthetic: IEPD tetrapeptide^10^ or natural ones: caspase-3 or BID^11^); therefore, the reliable detection of individual Grs with either chemical or antibody-related techniques remains challenging^10, 12^.

To date, only a few groups have succeeded in making functional activity-based probes for granzymes, but the major issues in these studies are the lack of selectivity of these probes toward individual enzymes and the low kinetic parameters^10, 13^. For example, an activity-based probe for granzyme B developed using combinatorial chemistry resulted in an inactivation rate constant for the target ABPs (GrB) equal to k_obs_/I = 460 M^−1^s^−1 13^.

Herein we design and characterize a set of selective chemical probes, including a quenched fluorescence substrate for GrB imaging in live cells. The development of new chemical tools for investigating Grs allows the examination of the unexplored functions of these proteases and their localization.

## Results

### Search for the optimal chain length for the substrate peptide

Our objective was to develop selective and potent substrates and probes for human granzyme B (GrB). Although the available GrB substrates are tetrapeptides, it was previously reported that lengthening the peptide substrate improves the substrate hydrolysis rate for some proteases^14^. To define the optimal peptide length for which GrB is most active, six fluorescent substrates were designed based on literature data. The S1 binding pocket of GrB has an unusual preference for aspartic acid^10, 15^; therefore, we incorporated Asp at P1, and we elongated the peptide to up to six amino acids. The investigated peptides were as follows: Ac-AAIEPD-ACC, Ac-AIEPD-ACC, Ac-IEPD-ACC, Ac-EPD-ACC, Ac-PD-ACC and Ac-D-ACC. We measured the activity of GrB against each of the new substrates (at equal concentrations) and observed that neither the tripeptide, the dipeptide or the single amino acid-based substrates were hydrolyzed. The longer tetra-, penta- and hexapeptides were cleaved by GrB. The pentapeptide and hexapeptides were more efficiently cleaved by GrB than was the classic tetrapeptide substrate (**Fig. 1A**). Because there was no significant difference in the efficiency of the cleavage of the penta- and hexapeptides, we selected the pentapeptides as the most appropriate chain length for activity-based probes and substrates.

**Figure 1.**
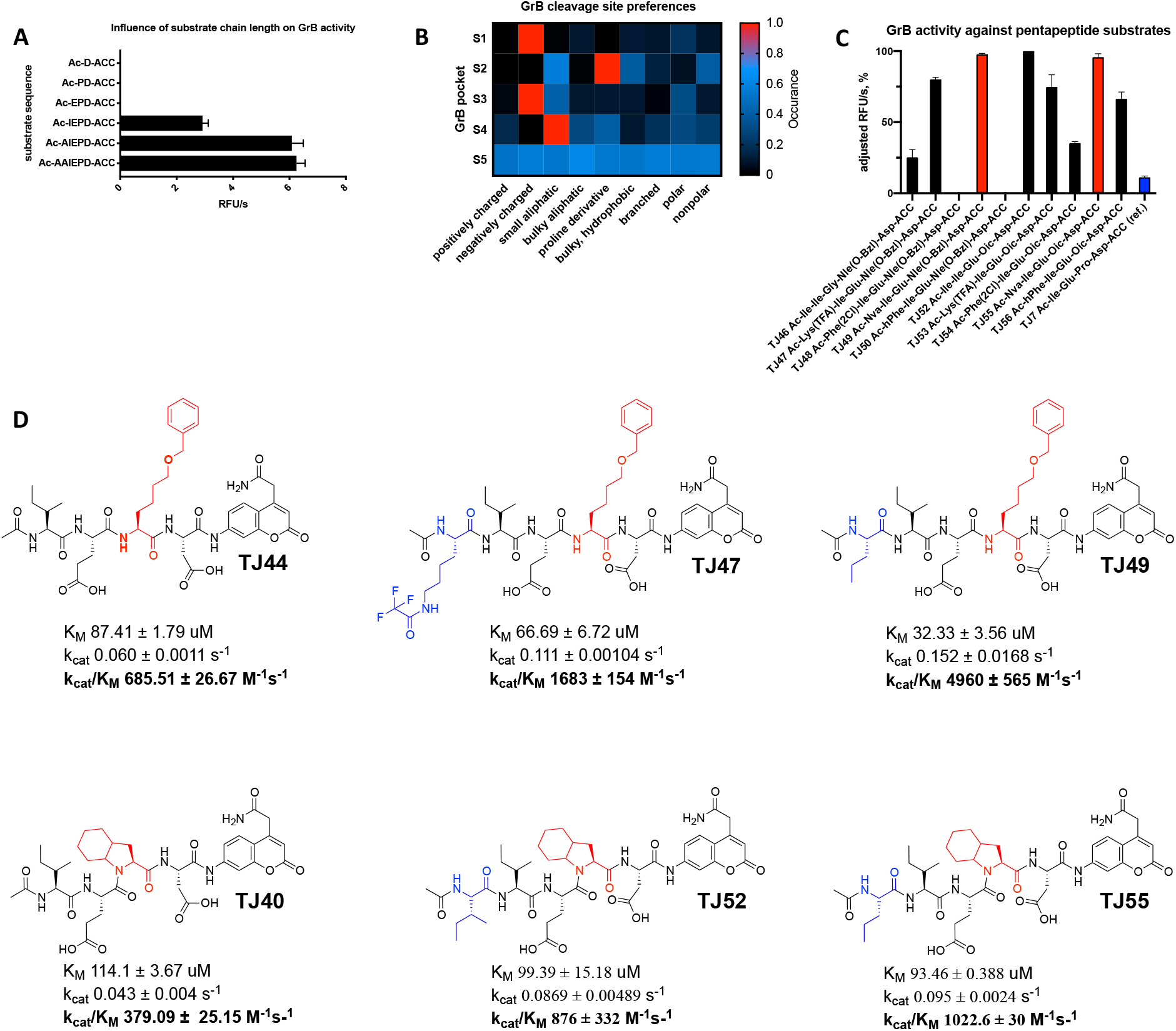
**A) Influence of substrate chain length on GrB activity.** The substrates (at the same concentrations) were treated with GrB. The data in the column diagram are presented as RFU/s values (relative fluorescence units/second). **B) GrB substrate cleavage site preferences.** The GrB specificity for the S4-S2 pockets was tested against Asp-HyCoSuL (Hybrid Combinatorial Substrate Library), while the GrB S5 pocket catalytic preferences were tested using the pentapeptides in the P5 library, which had the structure Ac-P5-X-Glu-X-Asp-ACC, and the P1 library, which had the structure Ac-Ile-Ser-Pro-P1-ACC. The results, shown as the relative fluorescent units per second (RFU/s), were divided into 9 groups according to the amino acid character and are presented as a heat map. The average value of RFU/s for each group was normalized with the highest value from all groups being equal to **C,D)** Screening, **structures of selected pentapeptide substrates and kinetic parameters.** Data are presented as the mean ± standard deviation and represent at least 2 independent experiments. All diagrams and k_cat_/K_M_ values were prepared using GraphPad Prism software.

### Catalytic preferences in nonprime S1-S5 enzyme pockets of GrB

To further explore the substrate specificity of GrB and allow better optimization of GrB substrates, we analyzed the S1-S5 pocket preferences using both combinatorial chemistry methods and the screening of defined peptides possessing natural and diverse unnatural amino acids^16, 17^.

The GrB S1 pocket almost exclusively recognizes aspartic acid, and this feature is also seen in cysteine protease caspases^9, 10, 17^. To reduce substrate cross-reactivity, especially with caspases that exhibit the most similar substrate preferences to GrB, we tested whether any acidic and nonacidic modifications of natural amino acid residues are accommodated by the GrB S1 pocket. For this purpose, we designed and synthesized (using a solid-phase peptide synthesis method, see Supplementary Data **Fig. S1**) a library of 95 defined peptides that share the same leading sequence and differ only by one amino acid residue at P1 (for the structures see Supplementary Data **Fig. S2**). ACC (7-aminocoumarin-4-acetic acid) was applied as a fluorescent leaving group, allowing the substrate hydrolysis rate to be determined based on the increase in fluorescence. The N-termini of the peptide substrates were acetylated to reduce the potential for hydrolysis by aminopeptidases commonly found in cells. In the remaining positions (P4-P2), defined natural amino acid residues were incorporated based on literature data (Ac-Ile-Ser-Pro-P1-ACC).

We tested the activity of GrB against the new P1 library and determined that GrB almost exclusively requires aspartic acid in the S1 pocket ((L-Asp, 100%); it can accommodate the methylated derivative (L-Asp(O-Me), 30%), and less potently, tyrosine (L-Tyr) and its derivatives (<10%) can be tolerated **(Fig. 1B and Fig. S3A)**. This confirms the literature data where, similar to caspases^10, 17^, the GrB S1 pocket is essentially restricted to aspartic acid due to its interaction with the positively charged guanidine group of Arg 226. Additionally, the carboxyl group of Asp forms hydrogen bonds with three water molecules within the S1 subsite of GrB^18^. Not only the charge but also the shape of this pocket dictates amino acid binding since only Asp or uncharged Asp derivatives with minimal modifications (Asp(O-Me)) can occupy this pocket, despite the lack of a negative charge, while more sizable Asp derivatives (Asp(O-Chx) and Asp(O-Bzl)) are not recognized by GrB, revealing the limited capacity of the S1 subsite and the involvement of additional interactions in peptide binding. The shape of Asp allows it to fit perfectly in the S1 pocket, and this is supported by the interaction between the guanidine group and the COOH group (Fig. S3).

Several approaches, including proteomics and PS-SCL (positional scanning substrate combinatorial libraries)^10, 19^, have demonstrated that GrB hydrolyzes after P1 Asp. Although this feature distinguishes GrB from other granzymes, many GrB substrates are also recognized by caspases^10^, so conventional strategies for enzyme activity analysis cannot be used to selectively follow GrB activity in cells. We therefore sought a substrate sequence that distinguishes GrB activity from caspases. To this end, we next determined the extended catalytic preferences of GrB based on S4-S2 using a well-established HyCoSuL strategy (Hybrid Combinatorial Substrate Library) incorporating a wide range of different nonproteinogenic amino acids^16, 17, 20^ **(Fig. S3)**. We observed that bulky hydrophobic proline derivatives such as octahydroindolecarboxylic acid (L-Oic), 1,2,3,4-tetrahydroisoquinoline-3-carboxylic acid (Tic) and O-benzyl-L-hydroxyproline (L-Hyp(Bzl)) are strongly preferred by GrB in P2, while substrates with hydroxyproline (L-Hyp) bearing unprotected hydroxyl groups are not hydrolyzed. Additionally, bulky hydrophobic 6-benzyloxy-L-norleucine (L-Nle(O-Bzl)), benzyl-L-histidine (L-His(Bzl)) and benzyl-L-serine (L-Ser(Bzl)) were tolerated in the S2 pocket, indicating that amino acids with benzyl groups can be accommodated within this pocket **(Fig. 1B, Fig. S3)** and revealing that the S2 pocket is very capacious. The crystal structure of GrB complexed with an inhibitor (Ac-IEPD-CO) with proline at P2 shows a cavity formed by the side chains of Phe 99, Tyr 94, Asp 102, His 57, and the main chain of Pro 96 is located beyond the pentameric pyrrolidine ring of the proline^18^. We speculate that this pocket is filled by the flexible cyclohexane group of proline derivative L-Oic or the benzyl groups of L-Nle(O-Bzl) or L-Ser(Bzl), enabling the amino acid residues to perfectly fill the S2 pocket.

The HyCoSuL strategy also revealed that the S3 pocket of GrB has a broad substrate scope; however, it has strong preferences for glutamic acid (L-Glu, 100%), hydrophobic tyrosine bearing a benzyl group (L-Tyr(Bzl)), 72%) and mono-oxidized methionine L-Met(O) **(Fig. 1B and Fig. S3)**. The strong preference for an acidic amino acid is due Asn 218 and Lys 192 in the S3 pocket of GrB, which interact with the side chain of L-Glu and stabilize the positively charged side chain of lysine^18^. Unlike the S2 pocket, the S3 pockets of all caspases^10^ share GrB’s preferences for glutamic acid, confirming the structural similarity of these enzymes^17^.

Our results demonstrated that the S4 pocket also possesses a broad substrate scope. It can accommodate branched amino acids such as isoleucine (L-Ile, 72%) or valine (L-Val, 36%) or linear hydroxyl-L-norvaline (L-Hnv, 82%) but also a proline derivative (1,2,3,4-tetrahydroisoquinoline-3-carboxylic acid, L-Tic, 100%), benzyloxymethyl-L-histidine (L-His(3-Bom), 82%) and some basic amino acids such as 2,4-diaminobutyrylic acid (L-Dab, 39%), citrulline (L-Cit, 10%) and 1,3-diaminopropionic acid (L-Dap, 10%) **(Fig. 1B and Fig. S3)**. According to Rotonda et al., this GrB pocket is a “shallow hydrophobic depression formed by aromatic rings” (Tyr174 and Tyr215) and the side chain of Leu172^18^; therefore, there is not enough space for phenylalanine within this pocket. Interestingly, our data from the HyCoSuL screening revealed that a bulkier amino acid (L-His(3-Bom)) was hydrolyzed by GrB (please see Supplementary Data **Fig. S3**).

To find the optimal amino acid for P5, we synthesized a combinatorial library of pentapeptides. For this purpose, based on the literature data and our results related to GrB specificity, we designed a library consisting of (1) defined amino acids at the P1 (Asp) and P3 (Glu), (2) equimolar mixtures (X) at P2 and P4 (a mixture of natural amino acids with L-Nle replacing L-Met and L-Cys), (3) and one of the 174 defined amino acids at P5 (Ac-P5-X-Glu-X-Asp-ACC) (for the structures, please see Supplementary Data Fig. S2). GrB was tested against the library, and we observed that it displays no S5 substrate specificity and is capable of hydrolyzing most substrates regardless of the residue at this position; however, the addition of an extra amino acid to the substrate (P5) clearly leads to a dramatic increase in substrate hydrolysis.

### Design, synthesis and kinetic analysis of pentapeptide substrates for GrB

To validate the results of the HyCoSuL screening, we selected the most promising amino acid residues for the S4-S2 positions (P4: L-Tic, L-His(3-Bom), and L-Ile; P3: L-Glu, and L-Tyr(Bzl); and P2: L-His(Bzl), L-Oic, L-Tic, and L-Hyp(Bzl)) and synthesized eighteen different fluorogenic tetrapeptides using a previously described method^21^. Afterwards, we tested the activity of GrB on the new substrates and observed that peptides with Ile at P4 were exclusively hydrolyzed by GrB (TJ40-44), while substrates with other selected amino acids, such as Tic (TJ2, TJ4, TJ6, and TJ35-39) and L-His(3-Bom) (TJ3 and TJ30-34), were not recognized by GrB under these assay conditions. Additionally, we confirmed that sequences containing L-Nle(O-Bzl) or L-Oic at S2 were preferred by GrB, confirming the large size of S2 **(Fig. S4A)**.

With this in mind, we performed a detailed kinetic analysis of the most hydrolyzed and promising substrates, namely, TJ40, TJ41, TJ42, TJ43, and TJ44 (Ac-Ile-Glu-Oic-Asp-ACC, Ac-Ile-Glu-Hyp(Bzl)-Asp-ACC, Ac-Ile-Glu-Tic-Asp-ACC, Ac-Ile-Glu-His(Bzl)-Asp-ACC, and Ac-Ile-Glu-Nle(O-Bzl)-Asp-ACC, respectively) (**Fig. S4**), and we compared those results with those of reference substrate TJ7 (Ac-Ile-Glu-Pro-Asp-ACC). Substrates TJ40, TJ43 and TJ44 were cleaved more rapidly by GrB than was TJ7, and their kinetic constants were k_cat_/K_M_ = 379.09 ± 25.15 M^−1^s^−1^, 267.89 ± 20.75 M^−1^s^−1^, and 685.51 ± 26.67 M^−1^s^−1^, respectively, and that of the reference is k_cat_/K_M_ = 76.95 ± 7.72 M^−1^s^−1^. This was in agreement with our initial substrate screening **(Fig. S4A)**. Since L-Oic and other proline derivatives are poorly or not recognized by caspases^10, 17, 22^, we decided to use the sequence of substrate TJ40 as the core for our future substrates and probes.

The optimization of the P4-P1 GrB peptide sequence resulted in a champion substrate that is well recognized by GrB and is less likely to show cross-reactivity with other granzymes and caspases. From the broad range of investigated amino acids, we selected the most promising structures (L-Lys(TFA), L-Nva, L-Ile, L-Phe(2-Cl) and L-hPhe), and we synthesized ten defined pentapeptide substrates, incorporating in the sequences previously selected for P4-P1 (L-Ile-L-Glu-L-Oic/L-Nle(O-Bzl)-L-Asp). After an initial screening of GrB activities against the new substrates (TJ46-TJ56) **(Fig. 1C)**, we selected substrates TJ47, TJ49, TJ52, and TJ55 as being hydrolyzed with the highest efficiency, and we studied their kinetics in detail **(Fig. 1D)**. Substrate TJ49 possesses the lowest K_M_ value (32.33 ± 10.81 μM) and, at the same time, the highest k_cat_ parameter (0.152 ± 0.017 s^−1^), with 40% higher activity (k_cat_/K_M_ of 4960 ± 565 M^−1^s^−1^) than that of the optimal tetrapeptide. Exchanging L-Nle(O-Bzl) at P2 (in TJ49) for L-Oic (affording TJ55) caused the enzyme activity to decrease by approximately a factor of four; however, we used this sequence in further analyses to minimize the cross-reactivity with majority of caspases^17^.

### Granzyme B activity-based probe design, synthesis and evaluation

The most valuable tools for GrB investigations are activity-based probes, both inhibitor- and substrate-based. However, because GrB and caspases share similar substrate specificity^10^, the development of potent and selective GrB probes is challenging. To address this limitation, we first designed an inhibitor-like molecule (activity-based probes) that is selective for GrB with low cross-reactivity with caspases.

We designed our probe to be built from three main functional segments. First, a reactive functional group that covalently binds to the active site of the enzyme. We selected diphenyl phosphonate, which mimics natural amino acids and potently reacts with serine proteases^23, 24^, thereby reducing the possibility of the probe binding with caspases. Second, we attached a peptide sequence to the warhead based on the optimal GrB catalytic preferences to ensure the best fitting of the probe within the active site of GrB. For this purpose, we selected our champion substrate (TJ55) (**Fig. 1D**) since GrB showed its highest activity with this substrate, and Oic at P2 disfavors binding with caspases. Third, we attached an affinity (biotin) (TJ55.Bt) or a fluorescent tag (Cy5) (TJ55.5) at the N-terminus for a pull-down assay or in-gel/membrane detection, respectively **(Fig. 2A)**. The probes were synthesized using a mix of classic solid-phase and solution-phase peptide synthesis methods. Briefly, the peptide sequence with Cy5 or biotin was attached to the warhead in solution following a procedure described elsewhere (see Supplementary Method **Fig. S5**). Using the same method, we obtained a covalent inhibitor of GrB with the general structure of Ac-Nva-Ile-Glu-Oic-Asp^P^(OPh)_2_. As we noticed from the calculation of the inhibition kinetic constants, GrB strongly prefers biotin as a tag relative to the Cy5 derivative, and it binds with TJ55.Bt almost 70 times more rapidly than it does to TJ55.5 **(Fig. 2A)**. We speculate that this substantial difference in enzyme binding to biotinylated versus fluorescent probes is due to the presence of a biotin binding exosite in the GrB structure, and biotin binding is preferred due to the three-dimensional enzyme structure and probe interactions.

**Figure 2.**
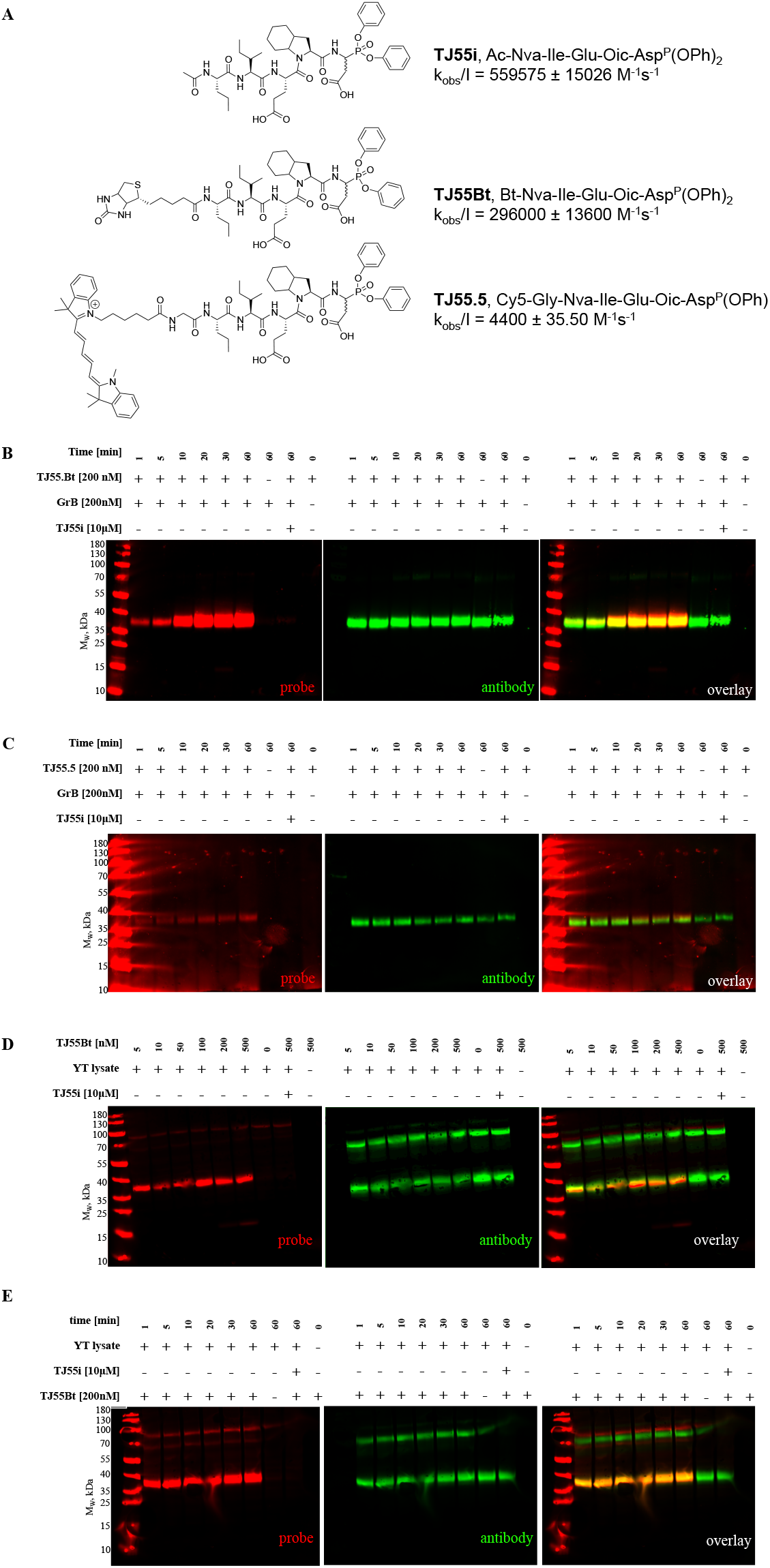
Specific granzyme B detection in cell lysates. A) Covalent GrB inhibitor and biotinylated and fluorescent activity-based probe structures and k_obs_/I values for GrB calculated using GraphPad Prism software. Data are reported as the mean ± standard deviation and represent at least 2 independent experiments. **B and C)** Optimization of the recombinant enzyme incubation time with TJ55.Bt (A) or TJ55.5 (B). GrB was incubated with TJ55.Bt or TJ55.5 for the indicated times (only the probe or only the enzyme was used in the controls). Afterwards, samples were analyzed by SDS-PAGE, followed by transfer to the membrane and streptavidin conjugate and antibody labeling. **D and E)** YT cell lysates corresponding to 1 × 10^7^ cells/mL were treated with TJ55.Bt for the indicated time (C) or samples were incubated with varying concentrations of GrB probe (from 5-500 nM) for 60 minutes (D). Afterwards, samples were analyzed by SDS-PAGE followed by transfer to the nitrocellulose membrane and immunoblotting using anti-GrB. As a control, lysates were pretreated with a competitive inhibitor prior to probe addition, or the probe and the lysates were run separately. The data reflect at least three separate biological replicates.

We tested the binding of our new probes to purified GrB and showed that both probes were capable of GrB labeling **(TJ55.Bt Fig. 2B and TJ55.5 Fig. 2C)**. First, we optimized the enzyme-probe incubation time for detecting GrB. After 5 minutes of incubation, we noticed a strongly labeled species between 35 and 40 kDa, corresponding to the size of purified GrB **(Fig. 2B, lane 2 and Fig. 2C, lane 2)**. As a control, to test whether the probe covalently binds in the GrB active site, we preinhibited GrB with a covalent inhibitor (Ac-Nva-Ile-Glu-Oic-<u>Asp^P^(OPh)_2_</u>) prior to probe addition **(Fig. 2B, lane 8 and Fig. 2C. lane 8)**. The inhibitor prevented probe binding; therefore, we concluded that TJ55.Bt and TJ55.5 bind to the active site of GrB. Additionally, TJ55.5 binds less potently than TJ55.Bt, which is consistent with their inhibition kinetic constants **(Fig. 1A)**. In addition, to confirm that the 35 and 40 kDa species represent GrB, we applied an anti-GrB monoclonal antibody, and the signals from the probes and antibodies overlapped exactly **(Fig. 2B and 2C, yellow band)**, confirming that our probe labels GrB. Furthermore, to test whether the sample autofluoresces in our assay conditions, we ran one sample without an activity-based probe and only detected a signal from the antibody **(Fig. 2B, lane 7 and Fig. 2C, lane 7)**. Thus, TJ55.Bt and TJ55.5 (1) bind GrB; (2) bind to the GrB active site; and (3) can be utilized in future active GrB detection assays.

### Active GrB detection in cell lysates

Since our activity-based probes allow efficient GrB labeling, in the next step, we verified their specificity in a complex system of cell lysates. We selected the human NK cell line YT since it constitutively expresses and releases GrB^25^ to observe if the probe binds with enzymes or cellular components other than the target enzyme. First, we optimized the concentration of the probe for efficient GrB labeling **(Fig. 2C)** and found that 5 nM is sufficient for GrB detection in complex systems. As indicated, even at a very high probe concentration (500 nM) and a long incubation time (60 minutes, **Fig. 2C, lane 7**), the probe binds almost exclusively with GrB; a signal from an additional band between 70-100 kDa was observed, but it does not have proteolytic activity^26^. Additionally, we observed that TJ55.Bt probe binds with GrB from YT cell lysates rapidly, and the complex was detectable after one minute of incubation (**Fig. 2D, lane 2)**. Pretreatment of the cell lysates with a competitive inhibitor of GrB activity prevented probe from binding (**Fig. 2D, lane 9**). (The ~100 kDa species seen in Fig. 2D and E is a nonspecific product of the streptavidin reagent, as it appears in samples not containing TJ55.Bt (lane 7).)

To verify whether the observed species corresponds to the target enzyme, we performed additional staining using an anti-GrB monoclonal antibody, and we noted that the signals from the probe and antibody overlapped, confirming probe binding with GrB. We also observed an additional species at 70 kDa when using the antibody. We speculate that this represents a complex between GrB and the cytosolic serpin PI-9 (Serpinb9), which forms postlysis and is detected by the monoclonal antibody of GrB.

Since our probe is specific to GrB even in a complex system of YT cell lysates, we speculated that it can be applied for GrB detection in other cells, especially cancer cell lines. To test that hypothesis, we screened GrB activity in different cell lines, such as YT (as a control cell line), MDA-MB-231, Su-DHL-4, Jurkat-T, NK92, MG63, SEMK2, REM, and NALM-6, using classic SDS-PAGE, and we observed that GrB is present in its active form mainly in the NK-like cells YT and NK92. Interestingly, there was far less GrB detected in NK92 cells (below the detection level of the antibody), emphasizing the sensitivity of this active site probe.

GrB was also detected in leukemia cell lines SEMK2, REM, and NALM but only in an inactive form, as it was strongly labeled with the antibody and not labeled with the activity-based probe (**Fig. 3**). We think that this may be due to the presence of inactive GrB within these cells or nonspecific antibody binding.

**Figure 3.**
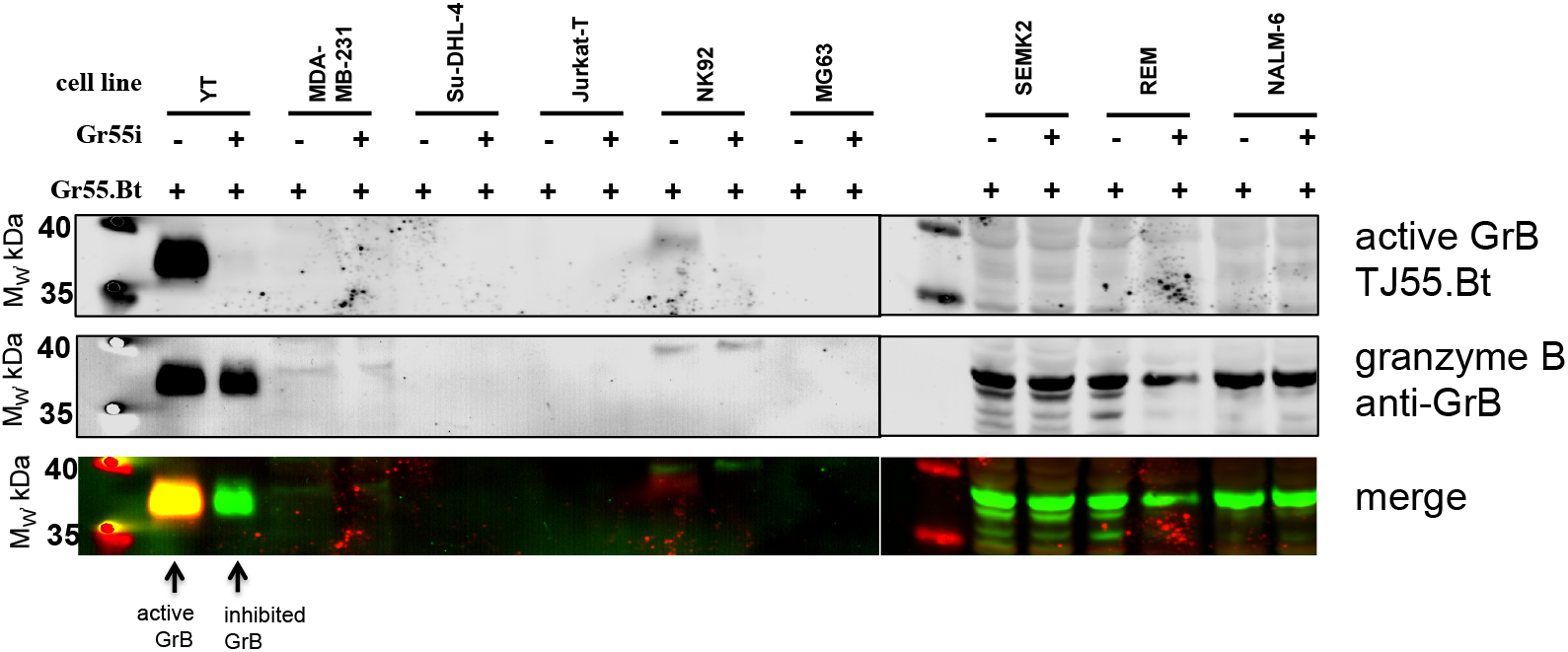
Detection of GrB in cell lines. **A)** Cells were harvested from liquid cultures and suspended to the same initial cell concentration (1 × 10^7^ cells/mL) in lysis buffer and then subjected to freeze-thaw cycles and sonication. Then, the same volume of each sample was treated with the same concentration of TJ55.Bt (250 nM) for 30 minutes. The samples were assessed by SDS-PAGE and transfer to a nitrocellulose membrane. The presented data are representative of at least two biological replicates.

### The design of a quenched fluorescence substrate probe for GrB

Biotinylated and fluorescent probes are valuable tools, but this type of inhibitor-like active site probe presents problems for enzyme monitoring in real time due to inhibition of the enzyme activity potentially leading to changes in its function. In addition, classic inhibitor-like probes are frequently equipped with unquenched fluorescence tags, which are always “on” despite the enzyme being active, and they emit a signal regardless of binding with the enzyme. This may cause false positive results since the cells, by pinocytosis or another mechanism, may take up the probe, and the signal will be detected regardless of binding with the target enzyme. Therefore, to avoid these limitations, we designed another type of molecule, a quenched fluorescent substrate probe, that emits fluorescence only after hydrolysis by the targeted enzyme. In addition, the probe will be substrate-based and therefore will not modify the biological function of the enzyme upon covalent binding.

To generate this probe, we first optimized its leading sequence. For this purpose, we used two previously optimized amino acid sequences for the P5-P1 positions based on TJ49 (Nva-Ile-Glu-Nle(O-Bzl)-Asp) and TJ55 (Nva-Ile-Glu-Oic-Asp), and we utilized variations of the P1’-P3’ sequence (Phe-Gly-Arg or Gly-Gly-Gly). Additionally, at the P6 position, we attached a PEG(4) linker (or not) to increase the distance between a recognition sequence and the fluorophore. To identify the most active sequence for GrB, we utilized ACC and Lys(Dnp) as the fluorescence donor-acceptor pair. After synthesizing the twelve substrates **(Table 1)**, we tested their hydrolysis rates by GrB, and we observed that in this type of GrB substrate, the predicted Oic (instead of the Nle(O-Bzl) group that was indicated in the HyCoSuL screening), is crucial at P2 for GrB substrate detection, and moreover, the S1’-S3’ pockets are significant for GrB activity. We selected two of the most promising sequences, TJ65: H_2_N-ACC-Peg(4)-Nva-Ile-Glu-Oic-Asp-Phe-Gly-Arg-Lys(Dnp)-CO(NH_2_) and TJ71: H_2_N-ACC-Nva-Ile-Glu-Oic-Asp-Phe-Gly-Arg-Lys(Dnp)-CO(NH_2_), and we calculated their exact kinetic parameters. Table 1 shows the structures and kinetic constants for all substrates. Regardless of the presence of a linker at P6, the substrates possess high k_cat_/K_M_ kinetic rates (TJ65 and TJ71). We also noticed a 33% reduction in K_M_ if PEG(4) was incorporated at P6. Surprisingly, the k_cat_/K_M_ for GrB on IQF substrates was dramatically higher than what was observed with shorter, classic substrates (TJ49 and TJ55). To test the selectivity of the most promising sequence (TJ71), we tested its activity with caspases since these enzymes share similar substrate specificity^10^, and we noticed robust activity with GrB but only minimal hydrolysis of TJ71 by caspase-6 and caspase-8 (**Fig. 4B and Table 2**); other caspases did not lead to fluorescence increases. Thus, the peptide sequence TJ71 was selected as a scaffold for future synthesis.

**Table 1.**
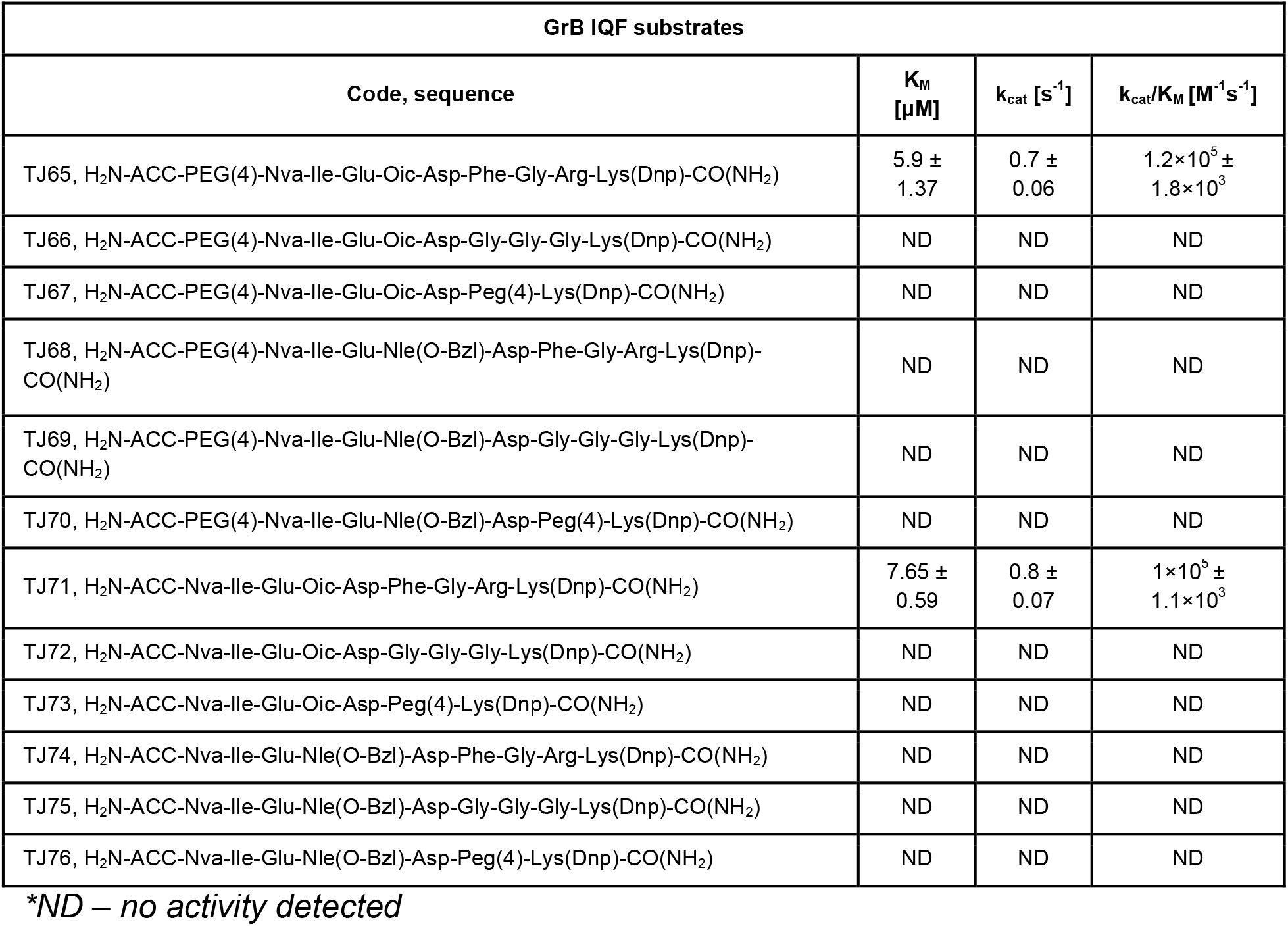
The structures of the GrB Internally quenched fluorescent (IQF) substrates and calculated kinetic constants for the hydrolyzed substrates. The k_cat_/K_M_ values were calculated using GraphPad Prism software. Data are the mean ± standard deviation and represent at least 2 independent experiments.

| GrB IQF substrates |  |  |  |
| --- | --- | --- | --- |
| Code, sequence | $K_M$<br>[$\mu M$] | $k_{cat}$ [$s^{-1}$] | $k_{cat}/K_M$ [$M^{-1}s^{-1}$] |
| TJ65, H<sub>2</sub>N-ACC-PEG(4)-Nva-Ile-Glu-Oic-Asp-Phe-Gly-Arg-Lys(Dnp)-CO(NH<sub>2</sub>) | 5.9 ± 1.37 | 0.7 ± 0.06 | 1.2×10<sup>5</sup> ± 1.8×10<sup>3</sup> |
| TJ66, H<sub>2</sub>N-ACC-PEG(4)-Nva-Ile-Glu-Oic-Asp-Gly-Gly-Gly-Lys(Dnp)-CO(NH<sub>2</sub>) | ND | ND | ND |
| TJ67, H<sub>2</sub>N-ACC-PEG(4)-Nva-Ile-Glu-Oic-Asp-Peg(4)-Lys(Dnp)-CO(NH<sub>2</sub>) | ND | ND | ND |
| TJ68, H<sub>2</sub>N-ACC-PEG(4)-Nva-Ile-Glu-Nle(O-Bzl)-Asp-Phe-Gly-Arg-Lys(Dnp)-CO(NH<sub>2</sub>) | ND | ND | ND |
| TJ69, H<sub>2</sub>N-ACC-PEG(4)-Nva-Ile-Glu-Nle(O-Bzl)-Asp-Gly-Gly-Gly-Lys(Dnp)-CO(NH<sub>2</sub>) | ND | ND | ND |
| TJ70, H<sub>2</sub>N-ACC-PEG(4)-Nva-Ile-Glu-Nle(O-Bzl)-Asp-Peg(4)-Lys(Dnp)-CO(NH<sub>2</sub>) | ND | ND | ND |
| TJ71, H<sub>2</sub>N-ACC-Nva-Ile-Glu-Oic-Asp-Phe-Gly-Arg-Lys(Dnp)-CO(NH<sub>2</sub>) | 7.65 ± 0.59 | 0.8 ± 0.07 | 1×10<sup>5</sup> ± 1.1×10<sup>3</sup> |
| TJ72, H<sub>2</sub>N-ACC-Nva-Ile-Glu-Oic-Asp-Gly-Gly-Gly-Lys(Dnp)-CO(NH<sub>2</sub>) | ND | ND | ND |
| TJ73, H<sub>2</sub>N-ACC-Nva-Ile-Glu-Oic-Asp-Peg(4)-Lys(Dnp)-CO(NH<sub>2</sub>) | ND | ND | ND |
| TJ74, H<sub>2</sub>N-ACC-Nva-Ile-Glu-Nle(O-Bzl)-Asp-Phe-Gly-Arg-Lys(Dnp)-CO(NH<sub>2</sub>) | ND | ND | ND |
| TJ75, H<sub>2</sub>N-ACC-Nva-Ile-Glu-Nle(O-Bzl)-Asp-Gly-Gly-Gly-Lys(Dnp)-CO(NH<sub>2</sub>) | ND | ND | ND |
| TJ76, H<sub>2</sub>N-ACC-Nva-Ile-Glu-Nle(O-Bzl)-Asp-Peg(4)-Lys(Dnp)-CO(NH<sub>2</sub>) | ND | ND | ND |

\*ND – no activity detected

**Table 2.**
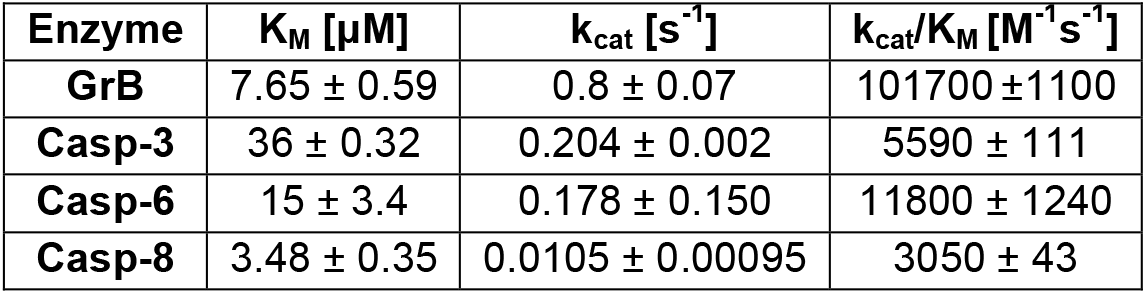
TJ71 specificity. k_cat_/K_M_ values were calculated using GraphPad Prism software. Data are the mean ± standard deviation from at least 2 independent experiments.

| Enzyme | $K_M$ [μM] | $k_{cat}$ [$s^{-1}$] | $k_{cat}/K_M$ [$M^{-1}s^{-1}$] |
| --- | --- | --- | --- |
| **GrB** | 7.65 ± 0.59 | 0.8 ± 0.07 | 101700 ± 1100 |
| **Casp-3** | 36 ± 0.32 | 0.204 ± 0.002 | 5590 ± 111 |
| **Casp-6** | 15 ± 3.4 | 0.178 ± 0.150 | 11800 ± 1240 |
| **Casp-8** | 3.48 ± 0.35 | 0.0105 ± 0.00095 | 3050 ± 43 |

**Figure 4.**
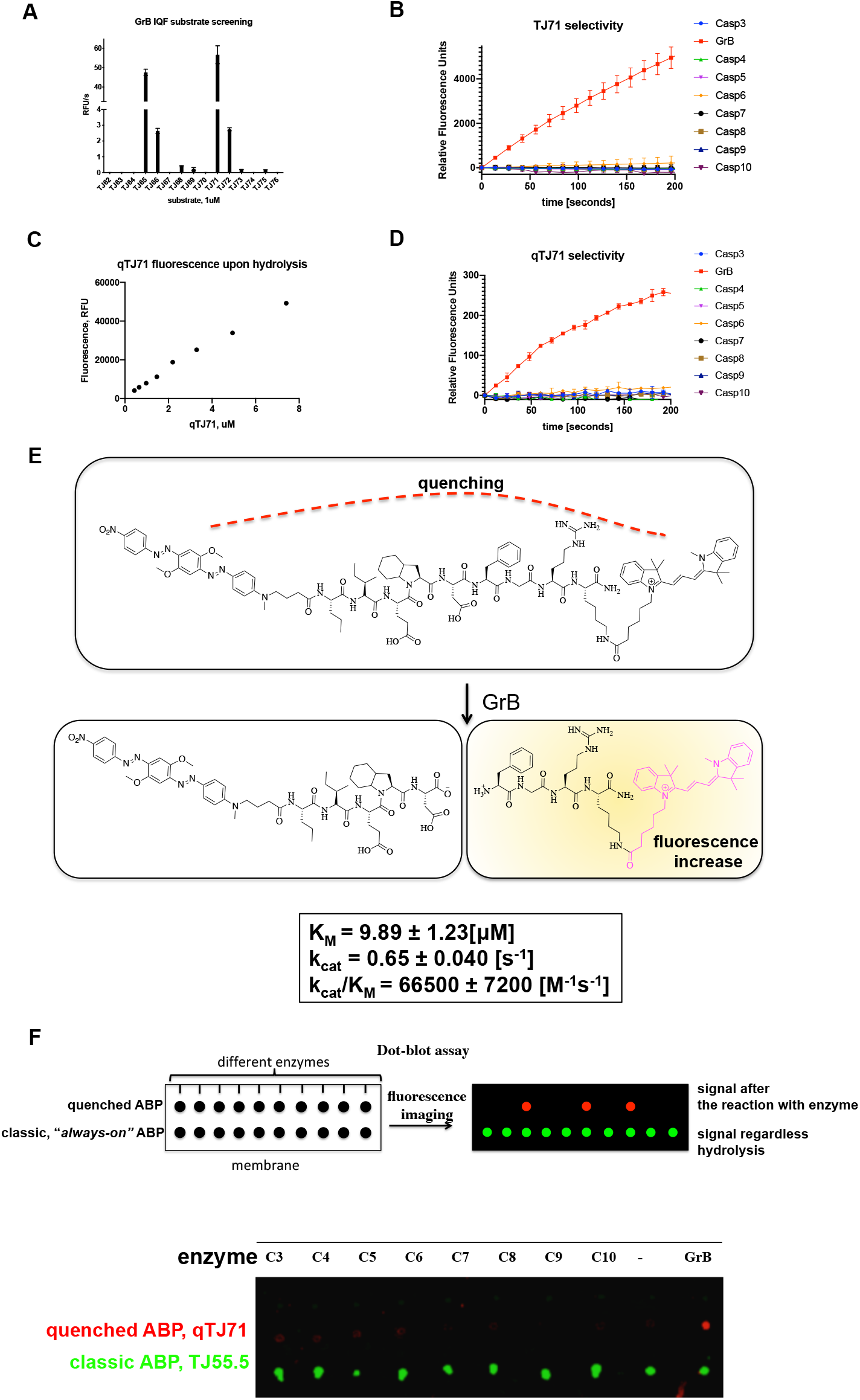
**A) Hydrolysis rate of GrB IQF substrates**. The increase in fluorescence over time was measured using a spectrofluorometer and analyzed in GraphPad Prism**. B) TJ71 selectivity.** Experiments were performed using GrB and caspases at equal concentrations. The increase in fluorescence over time was measured and analyzed in GraphPad Prism. **C) The fluorescence increase upon hydrolysis of qTJ71 is proportional to the substrate concentration. D) qTJ71 selectivity.** The experiments were performed with GrB and caspases at equal concentrations. The increase in fluorescence over time was measured and analyzed in GraphPad Prism. **E) A scheme of the qTJ71 cleavage site and kinetic parameters of the GrB hydrolysis of qTJ71** qTJ71 exhibits no fluorescent signal until it interacts with GrB. **F) Dot-blot analysis of quenched qTJ71 compared to unquenched TJ55.5.** The upper panel shows a scheme of the experiment demonstrating that the two types of chemical markers exhibit significantly different fluorescence properties upon interaction with the investigated enzymes. The bottom panel shows the test of qTJ71 utility in dot-blot analysis and its specificity determination. The classic inhibitor like activity-based probe (TJ55.5) with the unquenched fluorescent moiety (“always on”), regardless binding with the enzymes. All data (A-F) are the mean of at least two independent experiments performed in duplicate, and the standard error of the mean is provided.

### The GrB quenched fluorescent substrate is activated by recombinant human GrB

The ACC moiety is not a suitable fluorophore for the live imaging of enzymes within cells since its fluorescence emission is close to the natural autofluorescence of cells, which may cause a false positive result. Additionally, ACC possesses a poor quantum yield compared to other fluorophores; therefore, to use our **quenched fluorescent substrate** for in-cell GrB investigation, we exchanged ACC with cyanine derivative Cy3, which is more stable and convenient for cell-based analysis. Classic, “always-on” probes contain a fluorophore that exhibits a signal regardless of binding with the targeted enzyme. To prevent false positive signals and minimize the background fluorescence from the fluorophore of classic “always-on” probes, we applied a nonfluorescent quencher in the fluorescent substrate sequence that efficiently silences the fluorescence signal of the unhydrolyzed substrate. After hydrolysis, due to the separation of the donor-acceptor pair, Cy3 fluorescence is activated by the enzyme, and the substrate is released. We applied Black Hole Quencher^®^ 2 (BHQ2) as the quencher since it is characterized by a high quenching yield of Cy3, and we speculated that it can be utilized in cell-based analyses. The lack of measurable activity toward caspases made this sequence (TJ71) an ideal candidate for the recognition sequence of the quenched fluorescent substrate. Importantly, the fluorophore was attached to the C-terminus of the substrate and the quencher was attached to the N-terminus since upon substrate hydrolysis by an enzyme, the product containing Cy3 will be amplified in place of hydrolysis (**Fig. 4E**). The qTJ71 quenched fluorescent substrate was synthesized using a mixture of solid-phase and solution-phase synthesis techniques (**Supplementary data Fig. S7**). First, the peptide sequence was synthesized using classic SPPS on Rink amide followed by BHQ2 coupling at the N-terminus. Afterwards, the peptidic sequence with a quencher was cleaved from the resin, and Cy3-NHS ester was attached to the amine group of lysine with DIPEA/DMSO.

We tested the activation of qTJ71 by GrB utilizing kinetic analysis and as indicated in **Fig. 4C**. The total RFU from the entire peptide hydrolysis increased proportionally to the substrate concentration, confirming our hypothesis that qTJ71 is activated by GrB and that the fluorescence quenched by BHQ2 is released upon hydrolysis. Since caspases have also been reported to recognize Asp at the P1 position, we evaluated the hydrolysis of our substrate with these enzymes. For this purpose, we monitored the fluorescence increases as a function of time upon qTJ71 hydrolysis with GrB and caspases (at equal concentrations). As indicated in **Fig. 4D**, qTJ71 is hydrolyzed exclusively by GrB. Therefore, in the next step, we calculated k_cat_/K_M_ parameters for the hydrolysis of qTJ71 by GrB to be as high as 66500 M^−1^s^−1^. Next, we tested the cleavage site of qTJ71 by GrB, and observed that, as expected, the cleavage site was after the Asp residue (**Fig. 4E and Supplementary Data Fig. S8**).

qTJ71 was designed to be utilized in cell culture-based assays; however, first, we tested the utility of both quenched fluorescent substrate qTJ71 and unquenched TJ55.5 probe in a simple dot-blot assay with purified enzymes, and we confirmed if a signal specific to GrB could be detected and if any cross-reactivity of qTJ71 with caspases would be detected. Therefore, we added GrB or caspases-3, −4, −5, −6, −7, −8, −9, and −10 (separately) to the qTJ71 or TJ55.5, and afterwards, we performed dot-blot analysis. We observed increases in the signal only in the samples containing GrB and qTJ71, and as a control, we utilized an untreated quenched fluorescent substrate in an assay buffer and did not observe fluorescence. In the same test using unquenched TJ55.5, we observed a strong false positive signal in every sample from the so-called “always-on” activity-based probe (TJ55.5), in which fluorescence is emitted regardless of binding with the enzyme. Therefore, we concluded that our qTJ71 is selective for GrB and can be utilized in cell culture assays for real-time imaging of GrB within cells **(Fig. 4F)**.

### qTJ71 quenched fluorescent substrate cellular uptake

qTJ71 possesses selectivity toward GrB and is not recognized by active caspases; therefore, in the next step, we used it to follow GrB activity in YT cells. First, as a pilot experiment, we tested qTJ71 cellular uptake over time (**Supplementary data Fig. S9**). We performed live cell imaging of GrB in YT cells (in growth media) treated with 500 nM of qTJ71 and noticed that the quenched fluorescent substrate allows detection of GrB immediately after addition; however, the optimal labeling time for living cells was 15 minutes. Next, we addressed whether the fluorescence from qTJ71 cleavage is GrB-specific based on if it can be prevented by covalent inhibition of GrB. YT cells were pretreated with TJ55i and subsequently labeled with qTJ71. We observed a strong fluorescence increase in YT cells labeled with qTJ71 but no fluorescence in labeled cells pretreated with TJ55i, demonstrating that qTJ71 hydrolysis is prevented by the GrB covalent inhibitor (**Fig. 5A**). We concluded that this compound is selective to GrB and can be utilized for future GrB monitoring in living cells.

**Figure 5.**
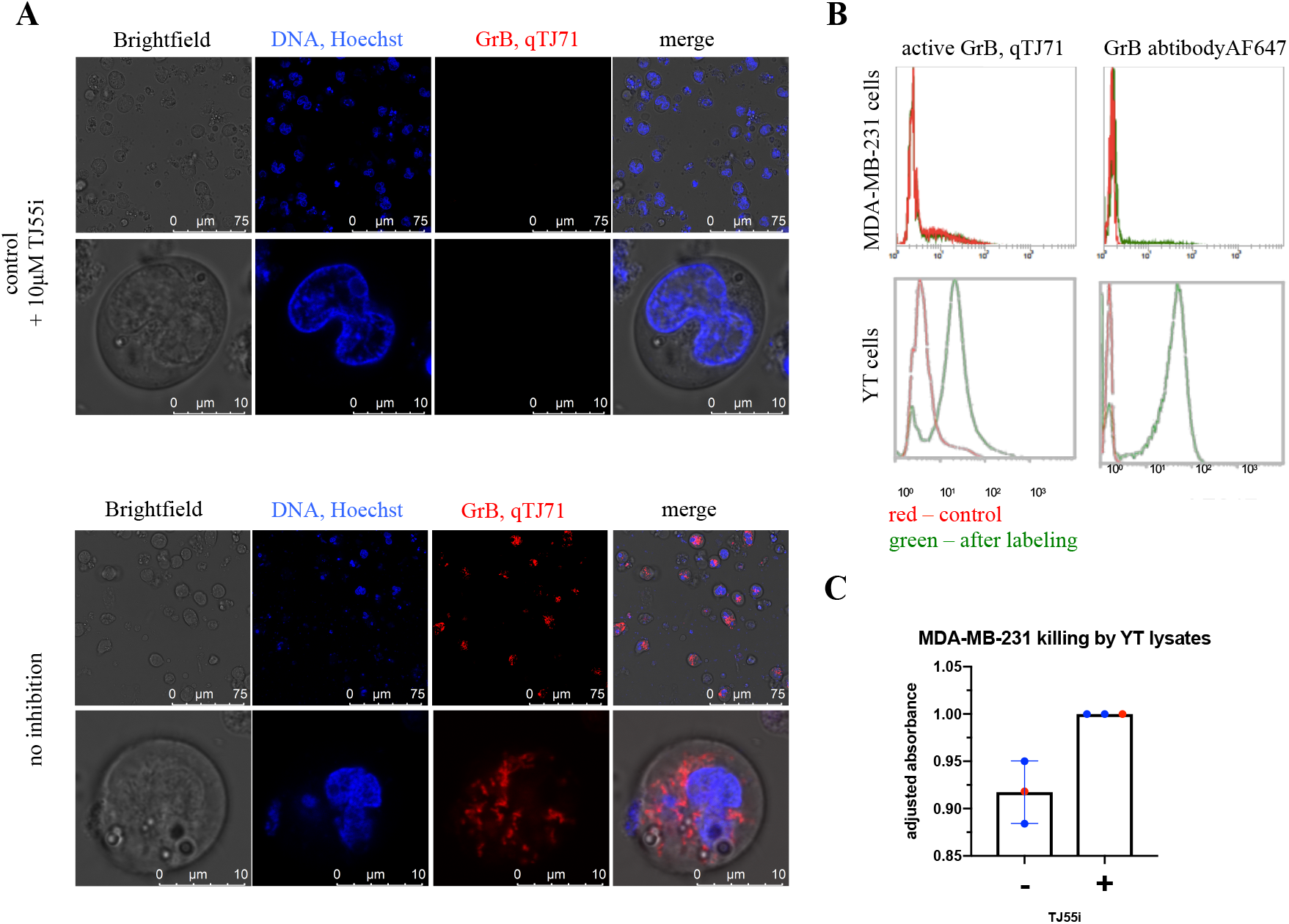
**A) Active GrB detection in YT cells with qTJ71 by confocal microscopy.** YT cells were treated with qTJ71 for 5 minutes and with Hoechst 33342 for 5 minutes followed by live imaging using a Leica TCS SP8 confocal microscope. As a control, cells were pretreated with GrB inhibitor prior to quenched fluorescent substrate addition. **B) GrB detection in YT cells and the absence of active GrB in MDA-MB-231 cells labeled with the qTJ71.** YT and MDA-MB-231 cells were stained with the qTJ71, fixed with PFA, and then analyzed at Sysmex CyFlow Cube. As a control, unlabeled cells and anti-GrB-labeled cells were analyzed. **C)** quenched fluorescent substrate. MDA-MB-231 cells were treated with YT cell lysates for 4 hours, and an MTS assay was performed. As a control, lysates were pretreated with TJ55i. Data are from 3 independent biological replicates.

### GrB activity monitoring in direct NK cell recognition of tumor targeted cells

GrB is considered to be involved in target cell killing, acting in concert with perforin, and by entering the target cell and, among other things, hydrolyzing caspase substrates, it can cause target cell death. Therefore, our next goal was to observe the accumulation of GrB activity in the breast cancer cell line MDA-MB-231 (target cells, T) after the addition of YT cells (effector cells, E). As indicated previously using the unquenched TJ55.5 probe (**Fig. 3**), active GrB is not usually present in MDA-MB-231 cells, while YT cells produce a substantial amount of this enzyme.

However, to test the utility of qTJ71 in GrB imaging in living cells, we performed two controls. First, GrB was inactivated using a covalent GrB inhibitor prior to the addition of the qTJ71 quenched fluorescent substrate. We observed no fluorescence, but when we used the qTJ71 alone, fluorescence was detected. This demonstrates that other components in the cell do not hydrolyze our quenched fluorescent substrate and it therefore can be used in cell-based assays. Second, to test the selectivity of the qTJ71, we treated both MDA-MB-231 and YT cell lines separately with the quenched fluorescent substrate and analyzed the samples by flow cytometry. We also analyzed naive cells and cells stained with anti-GrB followed by a fluorescently labeled secondary antibody. We observed robust fluorescence in YT cells treated with qTJ71, and in contrast, MDA-MB-231 cells showed no fluorescence. This was confirmed with antibody staining, which indicated the absence of GrB in MDA-MB-231 and its presence in YT (**Fig. 5B**). These experiments demonstrated that our quenched fluorescent substrate can detect active GrB within cells using simple and straightforward assays and classic methods such as FACS and microscopy (**Fig. 5A and B**).

GrB is considered one of the most important enzymes involved in target cell killing. Therefore, with a specific GrB inhibitor, we tested whether the inhibition of GrB in YT cell lysates influences its target cell killing efficiency. To do so, we measured MDA-MB-231 cell viability in the presence of YT cell lysates or YT cell lysates pretreated with GrB inhibitor (TJ55i). As indicated in **Fig. 5C**, the inhibition of GrB led to a 10% increase in cell viability but did not prevent target cell killing. With this in mind, we speculate that GrB is important but is not the only YT cell component that can cause target cell death.

## Materials and methods

### Chemicals

All chemicals were purchased from commercial suppliers and were used without further purification. All Fmoc-amino acids (purity > 99%) used for libraries and individual substrate synthesis were purchased from Iris Biotech GmbH, Combi-blocks, QMBIO, CreoSalus, Bachem, and APExBIO. Fmoc-Rink amide AM polystyrene resin (loading 0.74 mmol/g), DICI (diisopropylcarbodiimide, peptide grade), HBTU (*O*-benzotriazole-*N*,*N*,*N*’,*N*’-tetramethyluronium hexafluorophosphate, peptide grade), HATU (2-(1-H-7-azabenzotriazol-1-yl)-1,1,3,3-tetramethyl uronium hexafluorophosphate methanaminium, peptide grade), and TFA (trifluoroacetic acid, purity 99%) were purchased from Iris Biotech GmbH, Germany. HOBt (*N*-hydroxybenzotriazole, purity > 98%) was purchased from CreoSalus, USA. DCM (dichloromethane, analytically pure), MeOH (methanol, analytically pure), Et_2_O (diethyl ether, analytically pure), and AcOH (acetic acid, purity 99%) were purchased from POCh. DIPEA (*N*,*N*-diisopropylethylamine, peptide grade) was purchased from VWR International, Poland. Pip (piperidine, purity 99%), collidine (2,4,6-trimethylpyridine, peptide grade), and TIPS (triisopropylsilane, purity 99%) were purchased from Sigma Aldrich sp. z o.o., Poland. DMF (*N*,*N*’-dimethylformamide, peptide grade) and ACN (acetonitrile, HPLC grade) were purchased from Avantor, Poland. Triphenyl phosphite was purchased from Sigma Aldrich sp. z.o.o., Poland. BHQ-2 and BHQ-3 succinimidyl esters were purchased from Future Synthesis, Poland, Cyanine5 NHS ester was purchased from Lumiprobe, Germany.

All individual substrates were purified by HPLC (Waters M600 solvent delivery module, Waters M2489 detector system) using a C8 Supelco Discovery Bio Wide column (Sigma Aldrich). The solvents were as follows: phase A (95% water/0.1% TFA), phase B (5% acetonitrile/0.1% TFA). The purity of the substrates was determined by analytical HPLC (C8 Supelco Discovery Bio Wide analytical column, Sigma Aldrich). Finally, molecular weights were confirmed with a high-resolution mass spectrometer (Waters LCT Premier XE high-resolution mass spectrometer, electrospray ionization (ESI), and time-of-flight (TOF) detector) and LC-MS QDa (Waters e2695,2489 UV/Vis Detector, Acquity QDa detector).

### Kinetic assays

All kinetic experiments were performed using a spectrofluorometer (SpectraMax Gemini XPS, plate reader, Molecular Devices) and analyzed using SoftMax^®^ software (Molecular Devices), Microsoft Excel^®^ and GraphPad Prism^®^.

The granzyme B assay buffer contained 50 mM Tris-base, 100 mM NaCl, 25 mM CaCl_2_, 0.1% Tween, pH 7.4. The buffer was prepared at room temperature, and the enzyme kinetic studies were performed at 37°C.

a. **A HyCoSuL screening was utilized to define the substrate specificity of granzyme B**. P4, P3, and P2 fluorogenic substrate sublibraries (33 μM) with P1-Asp, with the general structure Ac-P4-X-X-Asp-ACC, Ac-X-P3-X-Asp-ACC, and Ac-X-X-P2-Asp-ACC (where P4-2 is a defined amino acid, “X” is an equimolar mixture of natural amino acids with Nle replacing Met and Cys) were scanned with GrB (48 nM). The final volume of the reaction mixture was 30 μL. The substrate hydrolysis (fluorescence release in the presence of enzyme) was monitored in real time at λ_ex_= 355 nm and λ_em_= 460 nm for a minimum of 30 minutes, but for each substrate, only the linear part of the curve was used to calculate the reaction rate (RFU/s, relative fluorescent units per second). Each sublibrary was screened twice, and the average value was calculated. For each sublibrary, the amino acid with the highest RFU/s was set to 100%, and the other amino acids were adjusted accordingly. The results are presented as a bar diagram (P4-P2) **(Supplementary Data Fig. S3).** Additionally, the adjusted values for amino acids sharing similar chemical properties were added and then divided by the number of events, showing the GrB preferences for a particular molecular feature. Data are presented as a heatmap (**Fig. 1A**). In the same manner, the P5 library (Ac-P5-X-E-X-D-ACC, 166 μM final concentration) was screened using 12 nM of GrB and analyzed as above. Data are presented as a violin diagram of GrB amino acid preferences.
b. **GrB activity against classic substrates (mono-, di-, tri-, tetra-, penta-, and hexapeptides).** The substrates TJ2-7 and TJ30-44 (166 μM) substrates were placed in a well of a 384-well plate (Corning^®^, opaque) and treated with 29 μL of 12 nM GrB. A total of 100 μM of Ac-D-ACC, Ac-PD-ACC, Ac-EPD-ACC, Ac-IEPD-ACC, Ac-AIEPD-ACC and Ac-AAIEPD-ACC were treated with 28 nM of GrB, and 166 μM of TJ46-56 substrate was placed in a well of a 384-well plate (Corning, opaque) and treated with 29 μL of 12 nM GrB. Substrate hydrolysis (fluorescence release in the presence of enzyme) was monitored in real time at λ_ex_= 355 nm and λ_em_= 460 nm for a minimum of 30 minutes. The linear region of the curve was used to calculate the reaction rate. RFU/s values were then adjusted to 100%, and the data are presented as bar graph with each value being the average of 3 replicates. For selected substrates, k_cat_/K_M_ values were calculated using substrates in a range of 666-5 μM for TJ7, TJ40, TJ43, and TJ44 or 333-2.6 μM for TJ47, TJ49, TJ52 and TJ55 with the Michaelis-Menten equation in GraphPad Prism. Data are presented as the mean with a standard deviation and represent at least 2 independent experiments.
c. **Quenched fluorescent substrate sequence optimization and k_cat_/K_M_ value determination**. IQF substrates TJ65-76 (6.6 μM) were screened with GrB (2.1 nM final) in 96-well Corning^®^ opaque plates with λ_ex_= 355 nm and λ_em_= 460 nm for a minimum of 30 minutes. Data were analyzed as above (“**GrB activity against classic substrates**”) and are presented as a bar chart. To calculate the k_cat_/K_M_ value, serial dilutions of TJ65 or TJ71 in a range from 14.81 μM to 0.86 μM were placed into a 96-well Corning^®^ black plate, and the enzyme at 12 nM in assay buffer was added. The fluorescence increase was monitored as a function of time for at least 30 minutes. The linear region of the curve was used to calculate the reaction rate using the Michaelis-Menten equation in GraphPad Prism. Data are presented as the mean with a standard deviation and represent at least 2 independent experiments.
d. **GrB activity against qTJ71.** The optimal excitation/emission wavelengths were determined to be λ_ex_= 540 nm and λ_em_= 580 nm. Serial dilutions of qTJ71 from 33.3 μM to 1.95 μM were placed in a 96-well Corning^®^ black plate, and 12 nM enzyme was added. The fluorescence increase over time was monitored for 30 minutes. The linear region of the curve was analyzed to calculate the reaction rate and analyzed using the Michaelis-Menten equation in GraphPad Prism. Data are presented as the mean with a standard deviation and represent at least 2 independent experiments.
e. **qTJ71 specificity.** Kinetic measurements were performed using black Corning^®^ plates with λ_ex_= 540 nm and λ_em_= 580 nm in assay buffers optimized for particular enzymes: (1) 50 mM Tris-base, 100 mM NaCl, 25 mM CaCl_2_, 0.1% Tween, pH 7.4 for granzyme B; (2) 10 mM Pipes, 100 mM NaCl, 1 mM EDTA, 10% sucrose, pH 7.4 for caspases 3-7; and (3) 10 mM sodium citrate, 10 mM Pipes, 100 mM NaCl, 1 mM EDTA, 10% sucrose, pH 7.4 for caspases 8-10. qTJ71 (1 μM) was placed into 96-well Corning^®^ black plates, and after the addition of the enzyme (120 nM), the fluorescence increase over time was monitored for 30 minutes. Only the linear regions of the curves were used for calculations. The k_cat_ and K_m_ values were calculated using GraphPad Prism and Microsoft Excel software. All measurements were repeated three times, and the data presented are averages of these replicates.

### Synthesis of P1 fluorogenic substrate library

To synthesize peptides with the sequence Ac-amino acids-ACC, we used the same method described previously^16, 21^ with small modifications. A total of 100 mg of Rink AM resin (0.74 mmol/g) was added to a glass reaction vessel containing DCM, and the mixture was stirred gently once every 10 minutes for 30 minutes. Then the mixture was filtered and washed 3 times with DMF. The Fmoc protecting group was then removed by treatment with 20% piperidine (v/v) in DMF for 5 minutes, 5 minutes, and then 25 minutes. After each cycle, the resin was washed with DMF and then filtered, and after the final deprotection, it was washed carefully six times with DMF. Afterwards, 2.5 eq of Fmoc-ACC-OH was preactivated with 2.5 eq of HOBt and 2.5 eq of DICI in DMF for 5 minutes, and that mixture was poured into the resin. The reaction was gently stirred for 24 hours at room temperature. The resin was then washed three times with DMF, and the Fmoc-ACC-OH coupling reaction was repeated using 1.5 eq of the above reagents to improve the yield of the coupling reaction. After this, the reaction mixture was removed, and the resin was washed five times with DMF, affording ACC-resin **(1)**. As previously described, to remove the Fmoc protecting group, 20% piperidine in DMF (v/v) was added to the reaction vessel, and three deprotection cycles were conducted (5 minutes, 5 minutes, and 25 minutes). The mixture was filtered and washed with DMF after each cycle to afford H_2_N-ACC-resin **(2)**. Next, 2.5 eq of Fmoc-P1-OH was preactivated with 2.5 eq of HATU and 2.5 eq of 2,4,6-trimethylpiridine in DMF, and this mixture was added to the reaction vessel with **2**. The reaction was carried out overnight at room temperature with gentle stirring. Then, the resin was washed with DMF three times, and the Fmoc-P1-OH coupling reaction was repeated using 1.5 eq of the reagents to increase the reaction yield and to obtain Fmoc-P1-ACC-resin **(3)**. The Fmoc protecting group was removed as described above, and the peptide chain elongation was continued using 2.5 eq of Fmoc-aa-OH (aa-amino acid), 2.5 eq of HOBt, and 2.5 eq of DICI until the desired peptide length was obtained. Each time, the coupling reaction or deprotection efficiency was tested using the ninhydrin test. At the end of this process, the N-terminus was protected with an acetyl group using 5 eq of AcOH, 5 eq of HBTU, and 5 eq of DIPEA in DMF with gentle stirring for 60 minutes at room temperature to afford Ac-peptide-ACC-resin **(4)**. Afterwards, the resin was washed five times with DMF, three times with DCM, and twice with MeOH and dried over P_2_O_5_ overnight. Next, peptide cleavage from the dry resin was performed with a mixture of cold TFA/TIPS/H_2_O (v/v/v; 95:2.5:2.5) for an hour at room temperature with gentle stirring once every 10 minutes. Then, the peptide was precipitated from cold Et_2_O for an hour and centrifuged. The supernatant was removed while the pellet was recrystallized from an additional portion of cold Et_2_O and centrifuged. The product, as a light-yellow pellet, was dried overnight at room temperature, purified by HPLC, and then lyophilized. Product purity was confirmed by analytical HPLC and HRMS analysis. All substrates were stored as 20 mM solutions at −80°C until use.

### Synthesis of internally quenched substrates

A total of 100 mg of Rink AM resin was added to a glass reaction vessel or 48-well reaction vessel containing DCM and stirred gently once every 10 minutes for 30 minutes, and then the mixture was filtered and washed 3 times with DMF. The Fmoc protecting group was then removed by treatment with 20% piperidine (v/v) in DMF for 5 minutes, 5 minutes, and 25 minutes. After each cycle, the piperidine was removed, and the resin was washed with DMF three times. Prior to transfer to the glass reaction vessel, 2.5 eq of Fmoc-Lys(Dnp)-OH was preactivated for 5 minutes by mixing with 2.5 eq of HOBt and 2.5 eq of DICI in DMF. Afterwards, the reaction was stirred for 12 hours at room temperature. The reaction mixture was filtered, and the resin was washed three times with DMF, and the Fmoc protecting group was removed from Fmoc-Lys(Dnp)-resin **(5)** as above using 20% piperidine in DMF (v/v) to afford H_2_N-Lys(Dnp)-resin **(6)**. Next, 2.5 eq of Fmoc-AA-OH (P3’ position) was preactivated with 2.5 eq HOBt and 2.5 eq DICI in DMF for 5 minutes and added to a reaction vessel with H_2_N-Lys(Dnp)-resin. The reaction was carried out for three hours with gentle stirring. Then, Fmoc-AA-Lys(Dnp)-resin **(7)** was filtered and washed with DMF three times. Afterwards, the Fmoc protecting group was removed as above, and the peptide chain elongation was continued until the peptide with the desired length was obtained (using 2.5 eq of HOBt and 2.5 eq of DICI as the coupling reagents). Next, 2.5 eq of Fmoc-ACC-OH was attached to H_2_N-peptide-Lys(Dnp)-resin **(8)** using 2.5 eq of HOBt and 2.5 eq of DICI, and the mixture was stirred for 24 hours at room temperature. Resin was then washed three times with DMF, and the reaction was repeated using 1.5 eq of the above reagents to improve the yield of the coupling of Fmoc-ACC-OH to **(8)**. The final product, H_2_N-ACC-peptide-Lys(Dnp)-resin **(9)**, was cleaved from Fmoc-ACC-peptide-Lys(Dnp)-resin protecting group and precipitated from Et_2_O as above. The product, a bright-yellow pellet, was dried overnight at room temperature, purified by HPLC, and lyophilized. Product purity was confirmed by analytical HPLC and HRMS analysis. All substrates were stored as 20 mM solutions at −80°C until use.

### Fluorogenic and biotinylated activity-based probe and inhibitor synthesis (TJ55.5: Cy5-Gly-Nva-Ile-Glu-Oic-Asp^P^(OPh)_2_, TJ55.Bt: Biot-Nva-Ile-Glu-Oic-Asp^P^(OPh)_2_ and TJ55i: Ac-Nva-Ile-Glu-Oic- Asp^P^(OPh)_2_)

In the first step, H_2_N-Nva-Ile-Glu-Oic-resin was synthesized using solid-phase synthesis on 2-chlorotityl chloride (CTC) resin. CTC resin (100 mg) was activated with 5 mL of anhydrous DCM and gently stirred once every 5 minutes for 30 minutes. Then, the mixture was filtered, and the filtrate was washed three times with DCM. Afterwards, 3 eq of Fmoc-Oic-OH (P2 position) was preactivated with 5 eq of DIPEA in anhydrous DCM and added to the glass reaction vessel with the CTC resin. The reaction was stirred gently in an argon atmosphere for 12 hours at room temperature. After that, the reaction mixture was filtered, and the resin was washed three times with DCM and two times with DMF to afford Fmoc-Oic-CTC **(10)**. The Fmoc protecting group was then removed from **10** using 20% piperidine (v/v) in DMF for 5, 5 and 25 minutes, as described above, to afford H_2_N-Oic-CTC **(11).** Peptide elongation with other amino acids (2.5 eq of P3: Fmoc-Glu(O-tBu)-OH, P4: Fmoc-Ile-OH, and P5: Fmoc-Nva-OH) was achieved with a series of coupling (with 2.5 eq of HOBt and 2.5 eq of DICI) and deprotection (20% PIP/DMF) reactions to obtain H_2_N-Nva-Ile-Glu(O-tBu)-Oic-CTC **(11).** Next, Boc-Gly-OH was introduced using 2.5 eq of HOBt and 2.5 eq of DICI (for TJ55.5) **(12)** or D-Biotin was introduced using 3 eq of DIPEA (for TJ55.Bt) **(13).** After the coupling of the last amino acid, the resin was washed five times with 2 mL portions of DMF, five times with 2 mL portions of DCM and three times with 2 mL portions of MeOH and dried over P_2_O_5_ overnight. The crude Boc-Gly-Nva-Ile-Glu(O-tBu)-Oic-COOH **(12)** and Biot-Nva-Ile-Glu(O-tBu)-Oic-COOH **(13)** peptides were cleaved from the resin using a mixture of TFE/AcOH/DCM (v/v/v, 1:1:8) for one hour. The supernatant was collected in a round-bottom flask, a portion of hexane was added, and the volatile products were evaporated under reduced pressure. As-obtained **13** was dissolved in a mixture of ACN/H_2_O (v/v, 3:1), frozen and lyophilized. The purities of **12 and 13** were confirmed with analytical HPLC, and the molecular weights were determined with HRMS.

The aspartic acid phosphonate warhead **(14)** was synthesized according to Mahrus et al. with slight modifictaions^13^. In a round-bottom flask, 8.64 g of Meldrum’s acid was mixed with 28 mL of triethyl orthoformate and stirred under reflux at 80°C. After three hours, the mixture was concentrated under reduced pressure to obtain a crude compound as a brown oil, which was stored at +4°C for 24 hours (it solidified at the temperature). The solid material (2.37 g) was vigorously stirred for one hour in 2 M HCl (34 mL). The suspension was then partitioned between diethyl ether and brine (3 × 100 mL). The organic phase was collected and dried over MgSO_4_, and the volatile components were evaporated under reduced pressure to give the product as a yellow solid. Next, to a 50 mL round-bottom flask were added formyl Meldrum’s acid (1.3 g, 7.5 mmol), p-nitrobenzyl alcohol (1.2 g, 7.5 mmol), and toluene (15 mL). The reaction mixture was stirred under reflux in an oil bath for 30 minutes, and afterwards, the solvent was removed under reduced pressure. The obtained crude oil (1.5 g) was used in the next step without purification. In the next step, in a 50 mL round-bottom flask, 1.5 g of *p*-nitrobenzyl formylacetate, 0.9 g of benzyl carbamate, 2.1 mL of triphenyl phosphite, and 2.1 mL of glacial acetic acid were stirred for 1 hour under reflux at 80°C, and then the solvent was removed under reduced pressure. The obtained product **(15)** was purified over SiO_2_ (2:3 ethyl acetate/hexane), and 1.2 g of pure compound was obtained. Afterwards, **15** was treated with 30% HBr in AcOH (10 mL) with stirring for 1 hour at room temperature. The solvent was then evaporated under reduced pressure, and the material was purified by reversed-phase HPLC. Compound **16** was analyzed by HRMS and stored at −80°C until use.

In the next step, 2.5 eq (25 mg) of the aspartic acid phosphonate warhead ( H_2_N-Asp^P^(OPh)_2,_ **16)** was coupled with **12 or 13** using 2.5 eq of HATU and 2.5 eq of collidine in DMF. Each reaction was carried out for 2 hours at room temperature. Afterwards, each mixture was diluted in ethyl acetate and washed 2 times with 5% NaHCO_3_ solution, 2 times with 5% citric acid solution, and 2 times with brine. The organic phase was collected and dried over MgSO_4_, and the volatile components were evaporated under reduced pressure. Next, deprotection of the aspartic acid carboxyl group from the warhead was performed by hydrogenolysis (H_2_, Pd/C, MeOH), and global deprotection was achieved in TFA/DCM (1:1, v/v; with 3% TIPS for 30 minutes) to afford H_2_N-Gly-Nva-Ile-Glu-Oic-Asp-Asp^P^(OPh)_2_ **(17) and** Biot-Nva-Ile-Glu-Oic-Asp-Asp^P^(OPh)_2_ **(18)**. Crude **17** was purified by HPLC, analyzed using HRMS and lyophilized. In the next step, 1 eq of Cy5-NHS was dissolved in DMF, 5 eq of DIPEA was added, and the mixture was stirred occasionally. This mixture was then added to pure, dry **17**. The reaction was carried out at room temperature for four hours with gentle stirring. The reaction progress was monitored by analytical HPLC. The final products, Cy5-Gly-Nva-Ile-Glu-Oic-Asp^P^(OPh)_2_ **(18)** and Biot-Nva-Ile-Glu-Oic-Asp^P^(OPh)_2_ **(19)**, were purified on HPLC, analyzed using HRMS, lyophilized, dissolved to a concentration of 20 mM in DMSO and stored at −80°C until use.

To obtain the inhibitor, the amine group of **11** was acetylated with 5 eq of AcOH, 5 eq of DIPEA, and 5 eq of HBTU in DMF. The reaction was carried out at room temperature for one hour. The remaining steps were analogous to the synthesis of the activity-based probe.

### qTJ71 synthesis (BHQ2-Nva-Ile-Glu-Oic-Asp-Phe-Gly-Arg-Lys-Cy3)

In the first step, a peptide derivative was obtained using solid-phase synthesis with Rink amide^®^ (AMR) resin in the same manner as above; however, a quencher was coupled to the *N*-terminus. For that coupling, 12.5 mg of Black Hole Quencher^®^ 2 (BHQ-2000S) was preactivated with 2.5 eq of DIPEA in DMF and added to a glass reaction vessel containing H_2_N-Nva-Ile-Glu(O-tBu)-Oic-Asp(O-tBu)-Phe-Gly-Arg(Pbf)-Lys(Boc)-AMR. The reaction was carried out at room temperature overnight, and the product was cleaved from the resin and purified as above. The purity of the obtained BHQ2-Nva-Ile-Glu(O-tBu)-Oic-Asp(O-tBu)-Phe-Gly-Arg(Pbf)-Lys(Boc)-NH_2_ **(19)** was confirmed by analytical HPLC, and the molecular weight of the compound was determined by HRMS. The protecting groups on the side chains of the amino acids in **(19)** were removed with TFA/DCM (1:1, v/v; supplemented with 3% of TIPS) in 30 minutes, and after deprotection, the volatile products were evaporated. Then, 1 eq of Cy3-NHS was attached to the side chain of Lys with 2.5 eq of DIPEA in DMF. The final product was purified with HPLC to afford BHQ2-Nva-Ile-Glu-Oic-Asp-Phe-Gly-Arg-Lys-Cy3 **(20)**. The compound was analyzed by HRMS, and stored as a 20 mM solution at −80°C until use.

### Inhibition kinetic assay

The inhibitory constants of TJ55i, TJ55.5 and TJ55.Bt were measured using Opaque Corning^®^ plates (Corning) with a spectrofluorometer (Spectramax Gemini XPS, Molecular Devices) and analyzed using SoftMax software (Molecular Devices) and Microsoft Excel^®^. The measurements were performed in an assay buffer containing 50 mM Tris-base, 100 mM NaCl, 25 mM CaCl_2_, 0.1% Tween, pH 7.4 at 37°C with excitation and emission wavelengths of 355 and 460, respectively, with a cutoff of 455 nm. To each reaction well was added 20 μL of an inhibitor at various concentrations (375 nM – 32 nM for TJ55i, 2 μM – 175 nM for TJ55.Bt and 166 μM −10 μM for TJ55.5) followed by 20 μL of substrate (TJ71, 24 μM) and then 60 μL of GrB (130 nM). The inhibitory efficiency and potency were calculated using k_obs(app)_/I (apparent second-order rate constant for inhibition) under pseudo-first-order conditions. k_obs_/I values were calculated taking into account the K_M_ value for the assay substrates (S) using the equation k_obs_/I = k_obs(app)_/I × [1+([S]/K_M_)].

### Dot-blot analysis of quenched fluorescent substrate and activity-based probe fluorescence and selectivity

Recombinant caspase-3, −4, −5, −6, −7, −8, −9, or −10, or granzyme B (120 nM) were each incubated with 1 μM of the quenched fluorescent substrate probe (qTJ71) or classic activity-based probe (TJ55.5) in an assay buffer (50 mM Tris-base, 100 mM NaCl, 25 mM CaCl_2_, 0.1% Tween, pH 7.4 for granzyme B; 10 mM Pipes, 100 mM NaCl, 1 mM EDTA, 10% sucrose, pH 7.4 for caspases 3-7; and 1 M sodium citrate, 10 mM Pipes, 100 mM NaCl, 1 mM EDTA, 10% sucrose, pH 7.4 for caspases 8-10) for 20 minutes at 37°C. Afterwards, 10 μL of each solution was dotted onto a dry nitrocellulose membrane (Bio-Rad, 0.2 μm) and allowed to dry for 5 minutes. Then, the membrane was scanned with a Sapphire− Biomolecular Imager with two lasers dedicated to Cy5 (658 nm) and Cy3 (520 nm), and the data were analyzed using Azure Biosystems software. The experiment was repeated 3 times.

### Cell culturing

SU-DHL1, YT, NALM-6 and Jurkat-T cells were cultured in 75 cm^3^ flasks in RPMI1640 medium supplemented with 10% fetal bovine serum and 1% penicillin/streptomycin; SEMK2 and MG63 were cultured on culture plates in Eagle’s minimum essential medium (EMEM) supplemented with 10% fetal bovine serum and 1% penicillin/streptomycin; and MDA-MB-231 cells were cultured on culture plates in Dulbecco’s modified Eagle’s medium (DMEM) supplemented with 10% fetal bovine serum and 1% penicillin/streptomycin under incubating conditions at 37°C, 90% relative humidity and 5% CO_2_. The primary concentration for optimal growth was 1 × 10^5^ cells/mL, and the media were changed every other day. For all further experiments, cells with a low passage number (up to 30) were taken approximately 24 hours after the last seeding.

### Western blot analysis of the cell lysates

To prepare the cell lysates, 1 × 10^7^ cells/mL were lysed with 1 mL of cold lysis buffer containing 15 mM KCl, 5 mM MgCl_2_, 10 mM Tris-HCl, 0.5% Triton-X 100 (v/v). Afterwards, the cells were sonicated (2.0 kJ for 10 seconds) and immediately treated with an activity-based probe (TJ55.5 or TJ55.Bt) for the indicated time (0 minutes to 1 hour) at 37°C. The reaction was stopped by the addition of 30 μL of 3 × SDS/DTT to 60 μL of sample (lysate + probe) followed by boiling at 95°C for 5 minutes. Afterwards, 30 μL of sample was loaded on 4-12% Bis-Tris Plus gel (Life Technologies); electrophoresis was performed at 200 V for 30 minutes, followed by transfer to a nitrocellulose membrane (0.2 μm, Bio-Rad, 1620112) for 60 minutes at 10 V. Then, the membrane was blocked with 2% BSA in TBS-T (Tris-buffered saline with 0.1% (v/v) Tween-20) for 60 minutes at room temperature, and when TJ55.Bt was used, the membranes were treated with fluorescent streptavidin conjugate (Steptavidin, Alexa Fluor− 647 conjugate, cat. no S21374, Invitrogen) (1:10 000) for 1 hour at room temperature, followed by rabbit recombinant monoclonal granzyme B antibody (Abcam, ab208586) and incubated overnight at 4°C. Then, the membrane was incubated with the secondary antibody (Alexa Fluor^®^532 goat anti-rabbit IgG [H+L], Invitrogen A11009) for 30 minutes at room temperature. The fluorescence was scanned at wavelengths of 649 nM for AF647 or Cy5 and 554 nM for AF532 using a Sapphire− Biomolecular Imager and Azure Biosystems software. The blots were then analyzed using Image Studio software.

### Flow cytometry

All experiments were performed using CyFlow Cube6 (Sysmex) and analyzed in Sysmex software. MDA-MB-231 cells (1 × 10^5^ cells/mL) in culturing media were incubated with (or without, as a control) 250 nM of qTJ71 for 1 hour. YT cells (1 × 10^5^ cells/mL) were incubated in cell culture media with (or without, as a control) 250 nM of qTJ71 for 1 hour. Afterwards, all samples were spun down, fixed with 4% PFA for 20 minutes, and washed twice with DPBS. Then, the cells were treated with 10% BSA in DPBS for 30 minutes, and the samples were spun down and treated with anti-GrB primary antibody (rabbit recombinant monoclonal granzyme B antibody, Abcam, ab208586) and incubated overnight at 4°C. After washing, DPBS secondary antibody (Alexa Fluor^®^532 goat anti-rabbit IgG [H+L], Invitrogen A11009) was added, and the mixture was incubated for 1 hour at 37°C. Then, samples were washed twice with DPBS, resuspended in 200 μL of DPBS, and analyzed with CyView^®^ software (Sysmex) using blue and red lasers and FL2 (580 nm) and FL4 (675 nm) filters. The experiment was repeated three times.

## Discussion

Granzyme B, a serine protease involved in programmed cell death through the cleavage of caspase substrates (BID and ICAM), is one of the key factors within NK cells and CTLs leading to cell death^10, 12 15^. GrB enters the target cell via perforin, and by hydrolyzing cellular proteins, it activates caspases, which are pivotal enzymes in cell death. The mechanism of GrB entrance into target cells using perforins remains unclear; however, a recent discovery led to the hypothesis that perforin may form a pore within the target cell that allows GrB to enter^27^. Within the cell, GrB is capable of cleaving different substrates, and since this enzyme can hydrolyze the same substrates as caspases, it can induce cell death through a caspase-independent cascade or the activation of caspases and is therefore involved in caspase-dependent cell death induction.

The classic artificial substrates of GrB described previously are tetrapeptide derivatives^13^. In our work, we analyzed the influence of peptide chain length with substrates containing from one to six amino acids. The incorporation of one or two additional amino acid residues increased GrB activity against the substrates, while shorter substrates (tri- or dipeptides) are not hydrolyzed by GrB. This analysis also indicated the importance of P4-Ile and P5-P6 amino acids in the interactions with the enzyme active site cavity and demonstrated that an additional amino acid reside at P5 of the substrate improves the interactions with the enzyme. We demonstrated that the P5 preference is broad but leans toward aliphatic residues such as Ile or Nva. Therefore, to construct our activity-based probe, we incorporated a pentapeptide as a leading sequence.

To precisely determine the GrB substrate specificity in the S1 pocket, we synthesized a set of substrates sharing the same P4-P2 sequence and differing at P1. We aimed to test whether Asp is the only amino acid that can be accommodated by the S1 pocket of GrB, and we observed that this enzyme almost exclusively hydrolyzes peptides after the Asp residue or its methylated derivative Asp(O-Me). This is due to the shape and chemical properties of the S1 pocket, which are controlled by the presence of a positively charged arginine 226 moiety oriented into the S1 cavity by the cis-proline conformation in the AA221-Pro224-Pro225-Arg226 motif, which is conserved between human and mouse GrB^18^. Therefore, GrB interacts with negatively charged Asp and its methylated derivative (Asp(O-Me)), but surprisingly, GrB does not cleave glutamic acid, which has one additional methylene group, and this is probably due to the shape and depth of the cavity, allowing for additional interactions with the Asp backbone.

On the other hand, the substrate specificity of the S2 pocket of GrB is similar to that of other serine proteases, and it recognizes proline and its derivatives. This specificity is characteristic of a majority of serine proteases, including the granzyme family^10^, neutrophil elastase^28^, cathepsin G^28^, proteinase 3^28^, chymotrypsin, and blood coagulation factor II. It is caused by Phe99 forming a wall in the active site of GrB, and as demonstrated previously, proline forms no specific interactions with the S2 pocket, but above the proline ring, an additional cavity was detected, and we speculate that this cavity can accommodate the proline extension because we observed that installing Oic at P2 dramatically improves substrate binding. Oic is a proline derivative modified with a cyclohexane ring that can adopt a three-dimensional structure known as chair conformation, giving it flexibility. Interestingly, bulky hydrophobic Nle(O-Bzl) at P2 was also recognized, confirming the presence of an additional cavity around the S2 subsite. In our opinion, the S2 pocket is crucial for distinguishing between granzyme B and caspases since it is the only position in which these enzymes possess distinct specificity. With our peptide library screening of GrB, we confirmed that it recognizes glutamic acid at P3 and shares this feature with caspases^10, 17, 22, 29^. The carboxyl group of glutamic acid interacts with Lys192 and Asn218 in the GrB S3 pocket, consistent with GrB specificity. Rotonda et al. demonstrated that the GrB S3 pocket is formed by Arg 41, Gly 43, Val 138, Gly 142, Gly 142, Gly 193, Pro 198, Tyr 32, Trp 141 and Met 30, and it is speculated that it possesses more than one subsite^18^. Likewise, S4 possesses a very narrow substrate specificity with a tendency to recognize small, aliphatic amino acids. This can be explained by the presence of the aromatic rings Tyr174, Tyr215 and Leu172 in the S4 pocket, making it a shallow and hydrophobic depression, as explained by Rotonda et al.^18^.

Our champion tetrapeptide sequence (TJ44) is hydrolyzed by GrB approximately 9-times more rapidly than are previously described GrB tetrapeptide substrates, while the pentapeptide substrate (TJ49) is approximately 65-times better, and the optimal sequence, containing Oic at P2, is 14-times better. Since longer peptides with an additional amino acid at P6 were not hydrolyzed more efficiently, we decided to utilize our optimal pentapeptide sequence for further probe construction.

Activity-based probes are the method of choice for the specific detection of GrB activity. Previously, Mahrus et al. synthesized a biotinylated activity-based probe with k_obs_/I 460 ± 35 that allowed GrB detection in NK cell lysates. In our work, we aimed to improve the kinetic parameters of activity-based probes for in-gel detection of GrB, and to do that, we synthesized two activity-based probes with the leading sequence based on the TJ55 substrate (1) with biotin (TJ55.Bt) or (2) with cyanine derivative Cy5 (TJ55.5), and we demonstrated the utility of these probes in GrB detection in both pure enzyme solutions and cell lysates. Since our new probes are very potent, with k_obs_/I = 296000 M^−1^s^−1^ for TJ55.Bt and k_obs_/I = 4400 M^−1^s^−1^ **(Fig. 2A)** for TJ55.5, we used them for GrB detection, and we observed that TJ55.Bt and TJ55.5 can immediately detect active enzyme (even after 1 minute) **(Fig. 2B and C)**. Because of caspase cross-reactivity with GrB substrates, we confirmed the specificity of our probes in a complex system, and because GrB is present in NK cells and CTLs, we selected the YT cell line as a model in further experiments. We demonstrated that both our probes (biotinylated and fluorogenic) possess high selectivity for GrB in a complex cellular environment since they label a protein with a size between 35-40 kDa (GrB migration that overlaps with a signal from GrB antibody). In addition, preincubation with a covalent GrB inhibitor prevented the binding of the activity-based probe, demonstrating that our probes attach to the GrB active site.

GrB is present within the NK and CTLs, and we aimed to test for its presence and activity in human cancer cell lines (YT, MDA-MB-231, Su-DHL-4, Jurkat-T, NK92, MG63, SEMK2, REM, and NALM-6). We demonstrated that, as indicated by TJ55.Bt, active GrB is almost exclusively present in the NK cell line YT, and a substantial amount of GrB can be detected in the NK92 cell line **(Fig. 3)**. This confirms the overall theory that NK cells are loaded with GrB and deliver it to the targeted cancer cells during invasion.

Activity-based probes are valuable tools for protease detection in cells and can be used to analyze enzyme functions in cells. However, because of their inhibitory activity, the utility of such probes is limited, and this type of molecule cannot be applied in live cell imaging because it may cause false positive readouts. To circumvent this, we designed a quenched fluorescent substrate that fluoresces only after hydrolysis by GrB. To do that, we started be designing design an optimal and selective peptide sequence by a design, synthesis and kinetic evaluation of internally quenched substrates. Afterwards, we selected the most promising structure, which was characterized by a dramatic increase in the k_cat_/K_M_ value, up to 100 000 M^−1^s^−1^ relative to classic substrates (approximately 5000 M^−1^s^−1^), and high specificity for GrB. We concluded that Oic at the P2 position is crucial for this specificity, while elongation with P1’-P3’ improves the enzyme activity. Consequently, we exchanged a fluorophore/quencher pair (Lys(DNP)-ACC) for Cy3/BHQ2 since it is more suitable for cell-based experiments. This new sequence was specific to GrB and was barely recognized by caspase-6 and caspase-8.

The activity of GrB depends on the pH, and GrB stored in low pH granules is not enzymatically active. However, after treatment of YT cells with our qTJ71, we detected a strong fluorescence, which was confirmed by FACS analysis **(Fig. 5A and B)**. Signal was dismissed by the presence of GrB inhibitor, confirming that hydrolysis occurs indeed by GrB. These data suggest that YT cells constitutively release active GrB via granule exocytosis, consistent with a previous study^25^. We speculate that the treatment of YT with granule destabilizer LLOMe should markedly increase GrB binding^30^.

GrB delivery to target cells was previously reported to follow a mechanism involving perforin, also known as a cytoplasmic granule toxin, which allows appropriate GrB entrance to the cells, and once inside the cell, GrB triggers apoptosis. It was demonstrated that both perforin (PFN-SG) and GrB (GrB-SG) form complexes with anionic serglycine (Ser-Gly), and although PFN-SG is less membranolytic than free PFN, the complex displays a similar or even greater ability to deliver GrB-SG and free GrB to the target cell. Additionally, membrane pore formation is not mandatory for GrB delivery and subsequent apoptosis^31^. The other possible mechanism involves GrB arginine and lysine residues on the surface structure that can bind to heparan sulfate^32^ or the negatively charged target cell membrane since the structure of GrB is highly basic (with a calculated pI of 10.4)^33^. We believe that our quenched fluorescent substrate will allow the monitoring of GrB delivery from NK cells to target cancer cell lines, which, as observed in MDA-MB-231 cells, do not have active GrB (**Fig. 5B**), and the observed fluorescence increase is associated with the substrate interaction with GrB.

In summary, in our work, we designed and obtained a set of specific tools for GrB investigation: (1) substrates, (2) an inhibitor, (3) inhibitor-based activity-based probes, and (4) a quenched fluorescent substrate. The designed compounds are characterized by high selectivity for GrB and potent activity. With this, we were able to detect active GrB in lysates from different cell lines, and we noted the presence of this enzyme in NK-like cells: YT and NK92.

## Acknowledgments

This work was supported by the HOMING Programme, a Grant Project of the Foundation for Polish Science, funded by the European Union under agreement No. 2016-3/24. PK is a beneficiary of L’Oreal Poland and Polish Ministry of Science and Higher Education scholarships.

