## Supplementary data for "Noninvasive optical detection of Granzyme B from natural killer cells using enzyme-activated fluorogenic probes"

**Non-invasive optical detection of killer cell Granzyme B using enzyme activated fluorogenic probes**

[1.] Wroclaw University of Science and Technology, Department of Chemistry, wyb. Wyspianskiego 29, 50-370 Wroclaw, Poland, [2.] Monash University, Monash Biomedicine Discovery Institute, Department of Biochemistry and Molecular Biology, 23 Innovation Walk, Clayton VIC 3800, Australia, [3.] NCI-designated Cancer Center, Sanford-Burnham Prebys Medical Discovery Institute, La Jolla, CA 92037, USA, [4.] Department of Pathology, University Medical Center Utrecht, Heidelberglaan 100, Utrecht 3584 CX, The Netherlands, [5.] Wroclaw Medical University, Department of Molecular and Cellular Biology, ul. Borowska 211A, 50-556, Wroclaw, Poland

Supplementary data

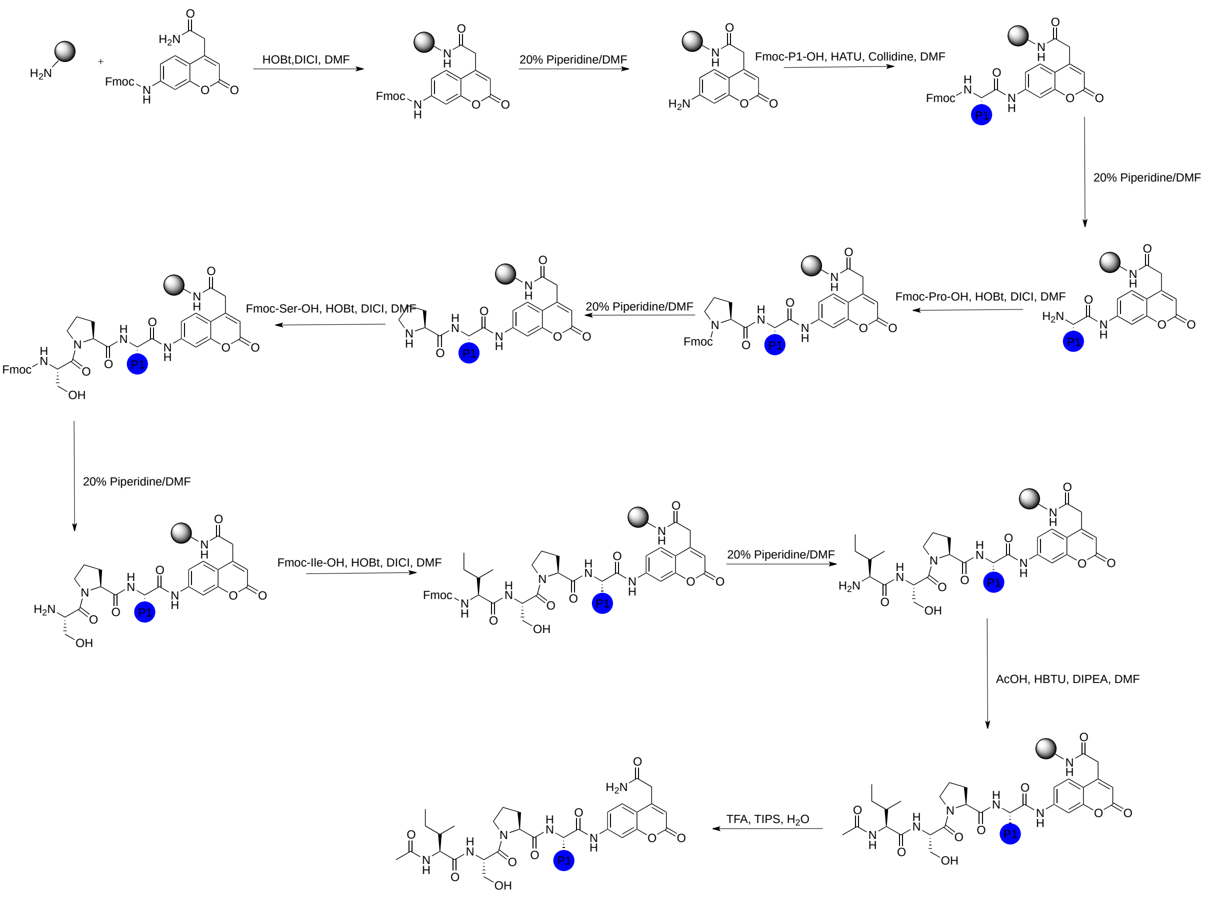


**Figure S1**. Solid phase peptide synthesis method. P1 library synthesis scheme.

### P1 and P5 LIBRARY STRUCTURES

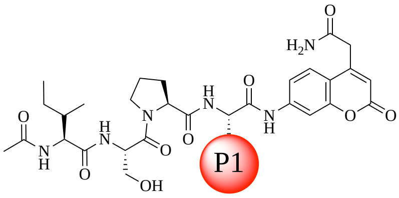


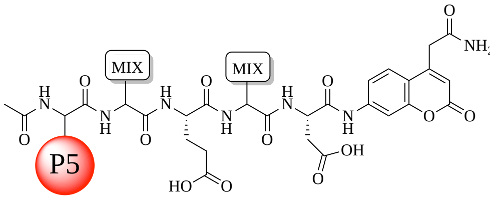


1 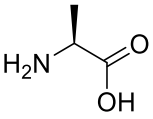 *L*-Ala

2 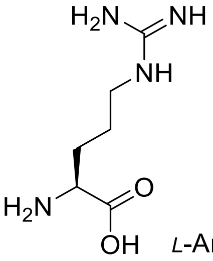 *L*-Arg

3 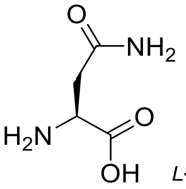 *L*-Asn

4 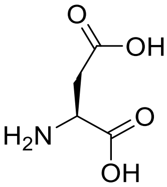 *L*-Asp

5 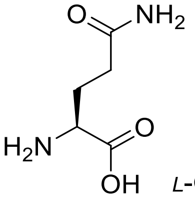 *L*-Gln

6 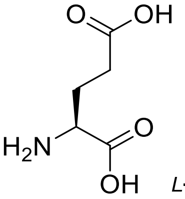 *L*-Glu

7 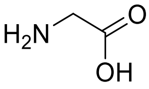 Gly

8 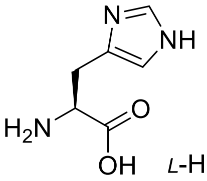 *L*-His

9 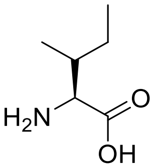 *L*-Ile

10 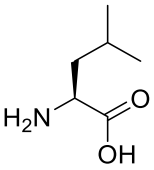 *L*-Leu

11 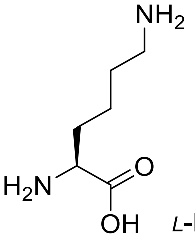 *L*-Lys

12 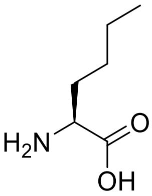 *L*-Nle

13 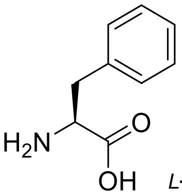 *L*-Phe

14 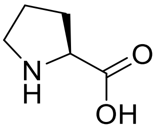 *L*-Pro

15 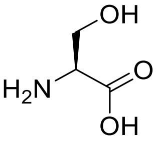 *L*-Ser

16 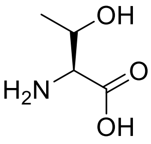 *L*-Thr

17 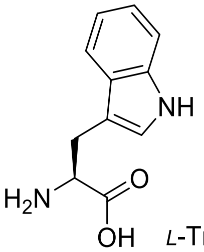 *L*-Trp

18 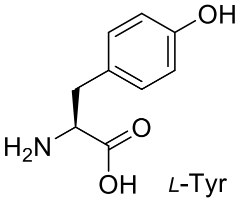 *L*-Tyr

19

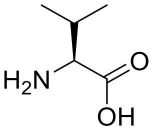


L-Val

20

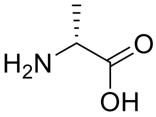


D-Ala

21

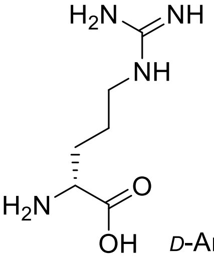


D-Arg

22

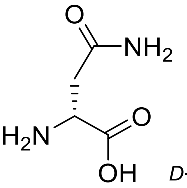


D-Asn

23

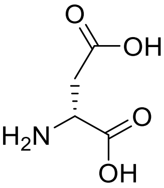


D-Asp

24

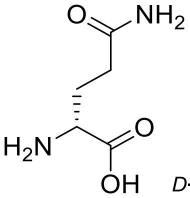


D-Gln

25

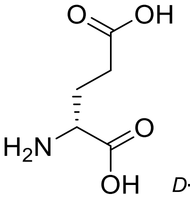


D-Glu

26

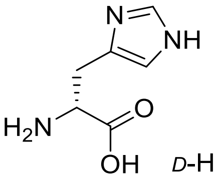


D-His

27

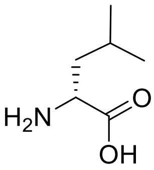


D-Leu

28

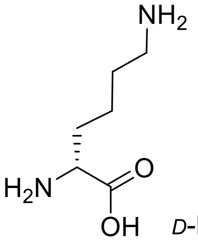


D-Lys

29

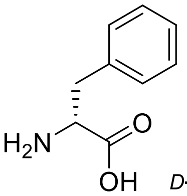


D-Phe

30

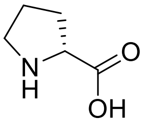


D-Pro

31

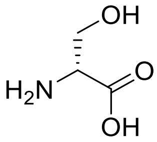


D-Ser

32

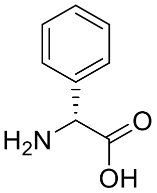


D-Phg

33

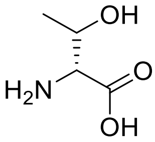


D-Thr

34

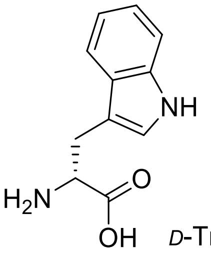


D-Trp

35

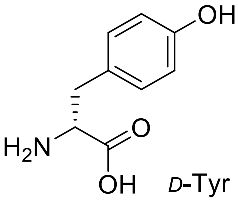


D-Tyr

36

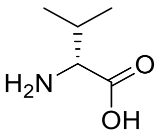


D-Val

37

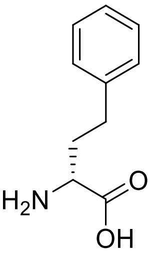


D-hPhe

38

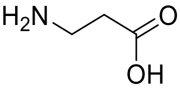


[illegible]-Ala

39

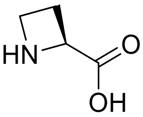


L-Aze

40 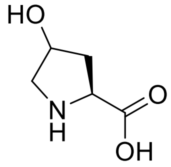 *L*-Hyp

41 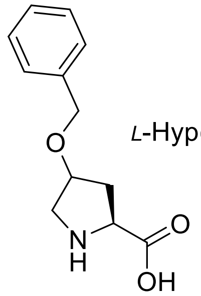 *L*-Hyp(Bzl)

42 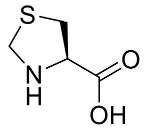 *L*-Thz

43 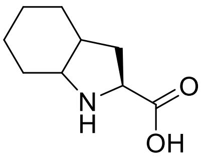 *L*-Oic

44 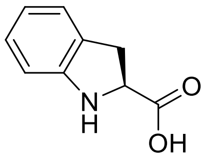 *L*-Idc

45 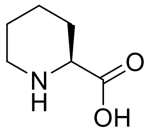 *L*-Pip

46 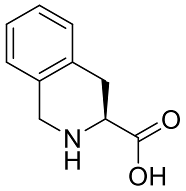 *L*-Tic

47 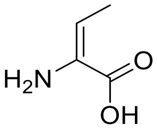 dhAbu

48 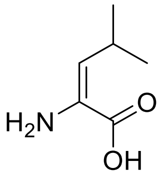 dhLeu

49 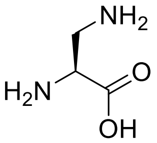 *L*-Dap

50 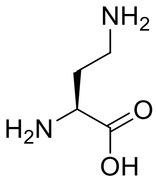 *L*-Dab

51 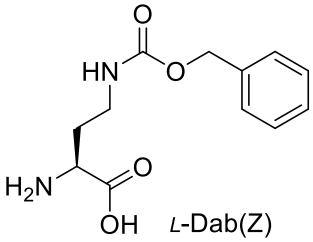 *L*-Dab(Z)

52 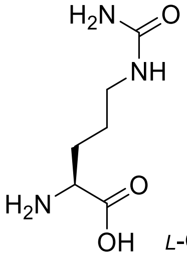 *L*-Cit

53 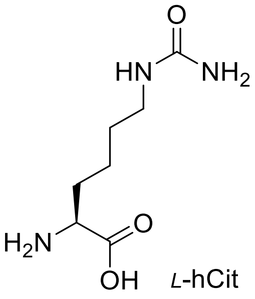 *L*-hCit

54 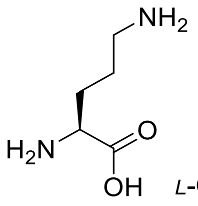 *L*-Orn

55 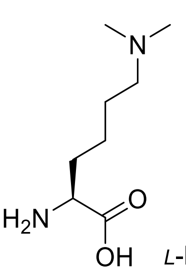 *L*-Lys(Me)$_2$

56 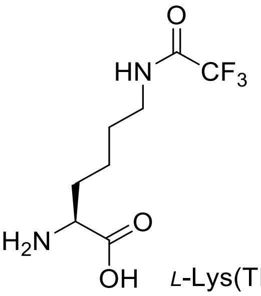 *L*-Lys(TFA)

57 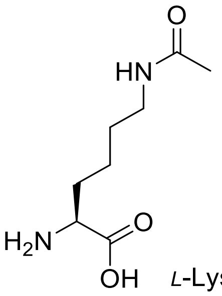 *L*-Lys(Ac)

58
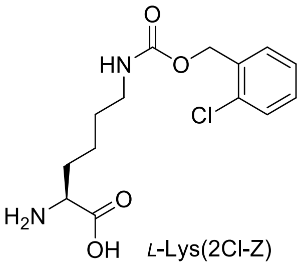
*L*-Lys(2Cl-Z)

59
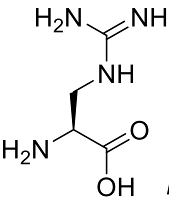
*L*-Agp

60
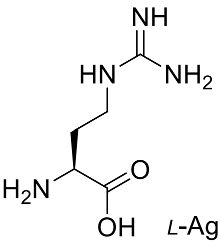
*L*-Agb

61
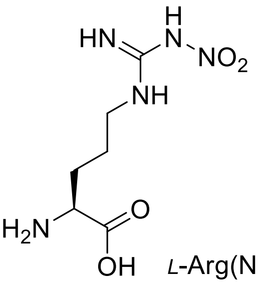
*L*-Arg($NO_2$)

62
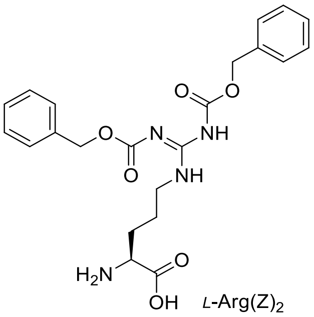
*L*-Arg(Z)$_2$

63
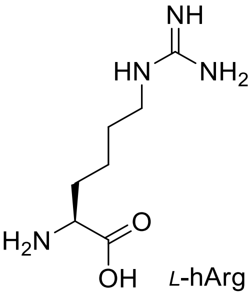
*L*-hArg

64
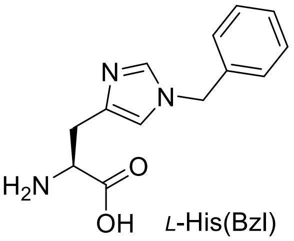
*L*-His(Bzl)

65
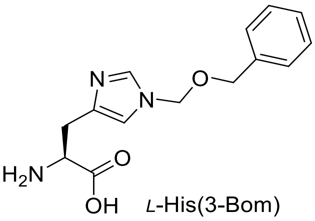
*L*-His(3-Bom)

66
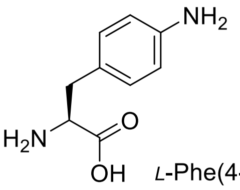
*L*-Phe(4-$NH_2$)

67
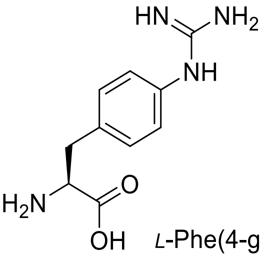
*L*-Phe(4-guan)

68
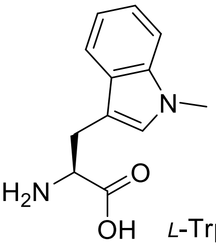
*L*-Trp(Me)

69
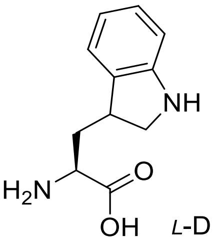
*L*-Dht

70
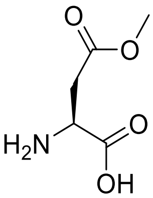
*L*-Asp(Me)

71
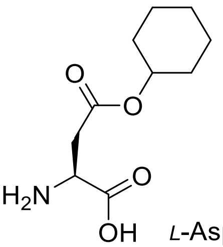
*L*-Asp(Chx)

72
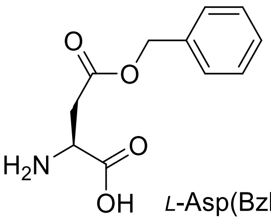
*L*-Asp(Bzl)

73
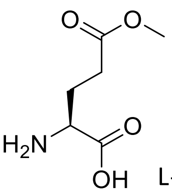
L-Glu(OMe)

74
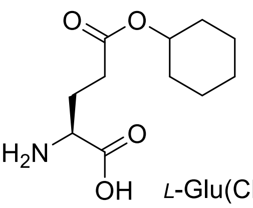
*L*-Glu(Chx)

75
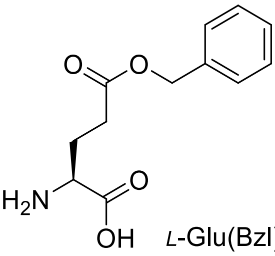
*L*-Glu(Bzl)

76 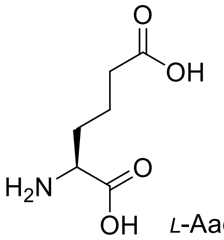 L-Aad

77 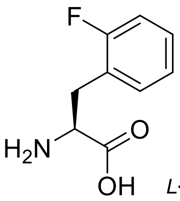 L-Phe(2-F)

78 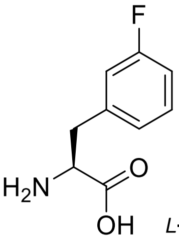 L-Phe(3-F)

79 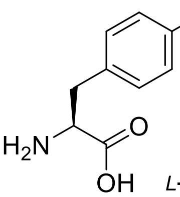 L-Phe(4-F)

80 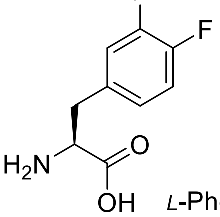 L-Phe(4,3-$F_2$)

81 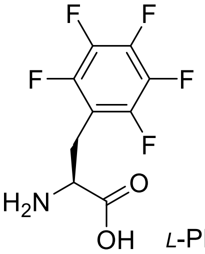 L-Phe($F_5$)

82 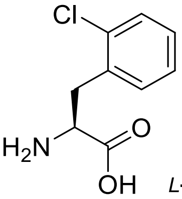 L-Phe(2-Cl)

83 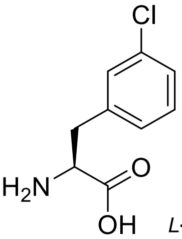 L-Phe(3-Cl)

84 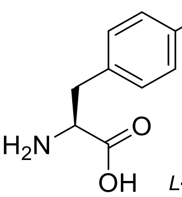 L-Phe(4-Cl)

85 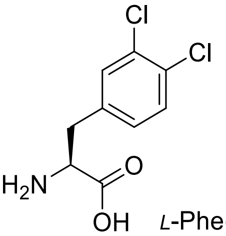 L-Phe(3,4-$Cl_2$)

86 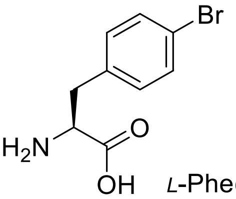 L-Phe(4-Br)

87 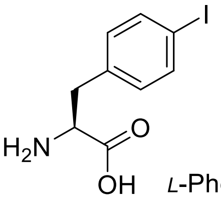 L-Phe(4-I)

88 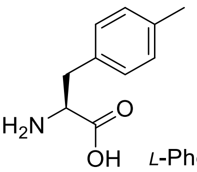 L-Phe(4-Me)

89 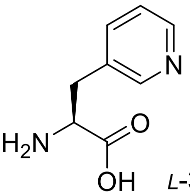 L-3-Pal

90 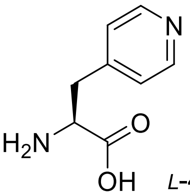 L-4-Pal

91 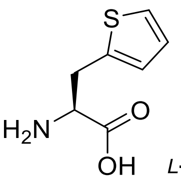 L-Ala(2-th)

92 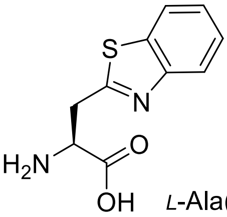 L-Ala(Bth)

93 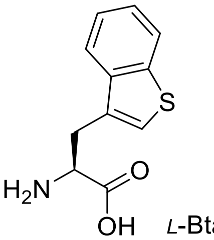 L-Bta

94 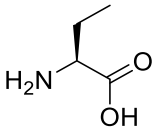 *L*-Abu

95 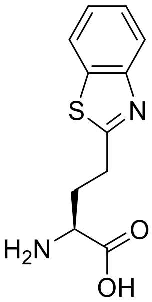 *L*-Abu(Bth)

96 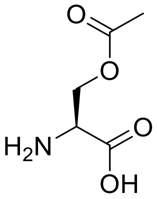 *L*-Ser(Ac)

97 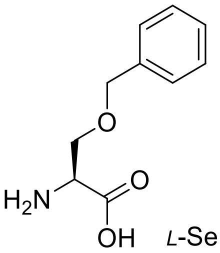 *L*-Ser(Bzl)

98 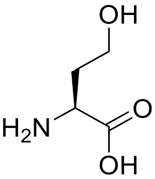 *L*-hSer

99 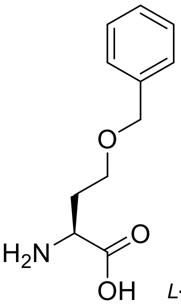 *L*-hSer(Bzl)

100 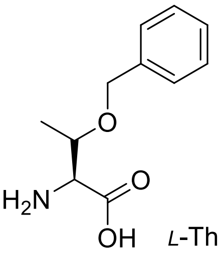 *L*-Thr(Bzl)

101 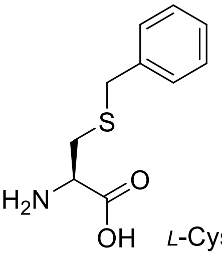 *L*-Cys(Bzl)

102 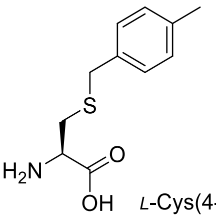 *L*-Cys(4-MeBzl)

103 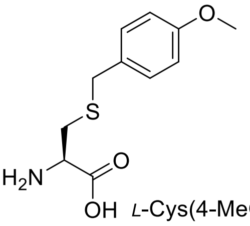 *L*-Cys(4-MeOBzl)

104 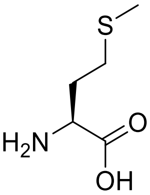 *L*-Met

105 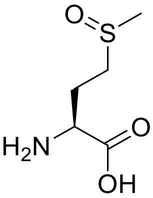 *L*-Met(O)

106 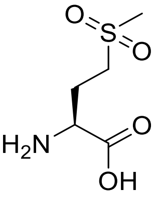 *L*-Met($O_2$)

107 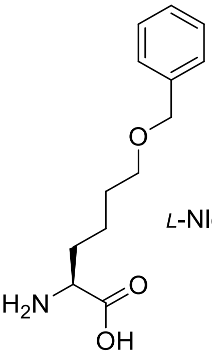 *L*-Nle(*O*-Bzl)

108 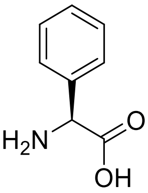 *L*-Phg

109

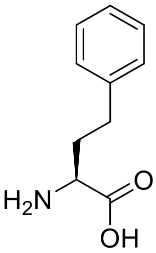

*L*-hPhe

110

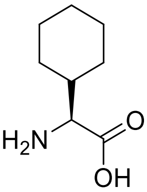

*L*-Chg

111

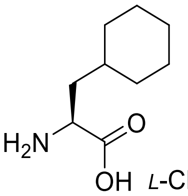

*L*-Cha

112

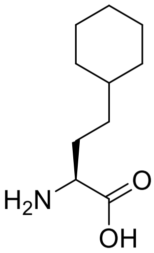

*L*-hCha

113

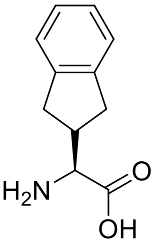

*L*-Igl

114

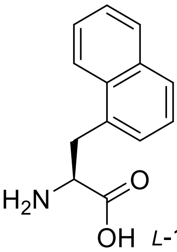

*L*-1-Nal

115

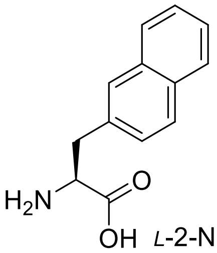

*L*-2-Nal

116

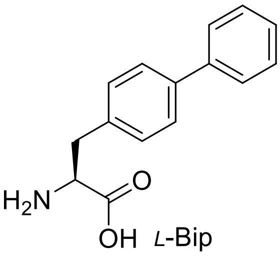

*L*-Bip

117

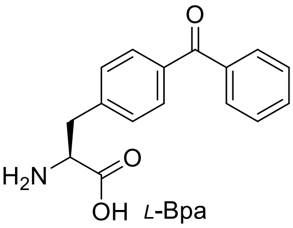

*L*-Bpa

118

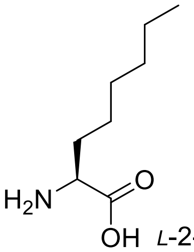

*L*-2-Aoc

119

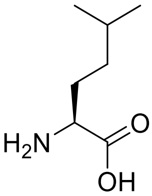

*L*-hLeu

120

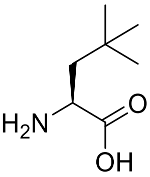

*L*-NptGly

121

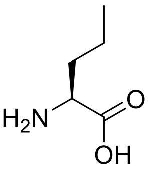

*L*-Nva

122

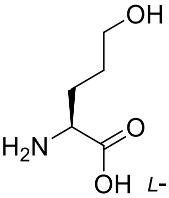

*L*-Hnv

123

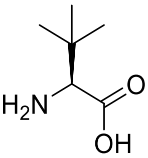

*L*-Tle

124

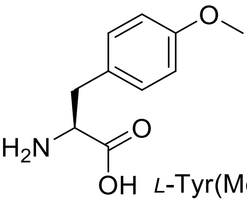

*L*-Tyr(Me)

125

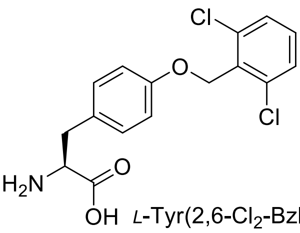

*L*-Tyr(2,6-Cl$_2$-Bzl)

126

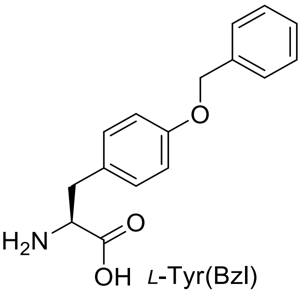

*L*-Tyr(Bzl)

127 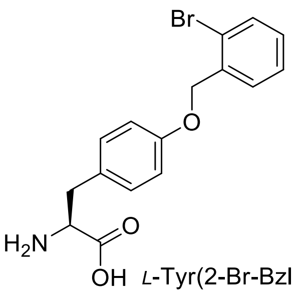 *L*-Tyr(2-Br-Bzl)

128 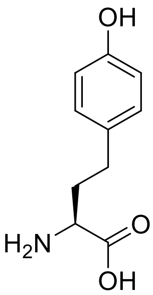 *L*-hTyr

129 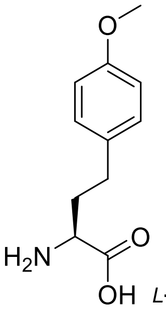 *L*-hTyr(Me)

130 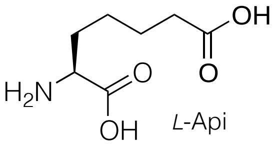 *L*-Api

131 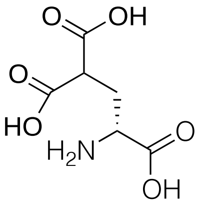 *D*-Gla

132 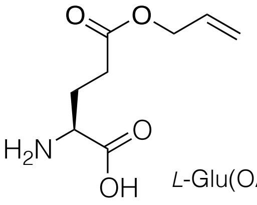 *L*-Glu(OAll)

132 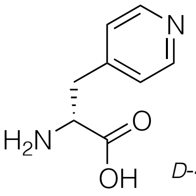 *D*-4-Pal-OH

133 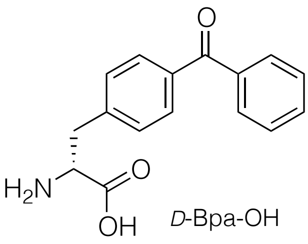 *D*-Bpa-OH

134 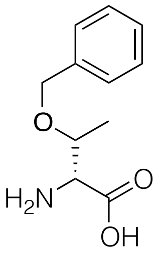 *D*-Thr(Bzl)

135 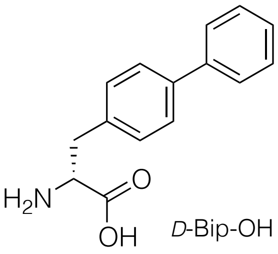 *D*-Bip-OH

136 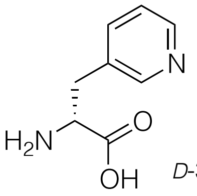 *D*-3-Pal-OH

137 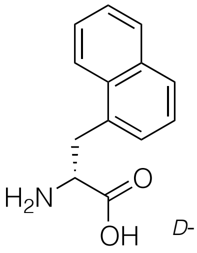 *D*-1-Nal

138 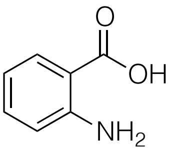 2-Abz

139 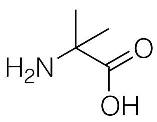 *Aib-OH*

140 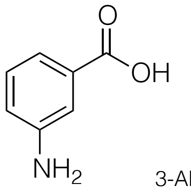 3-Abz

141 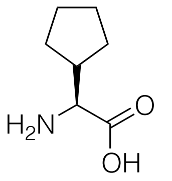 *L-Cpg-OH*

142 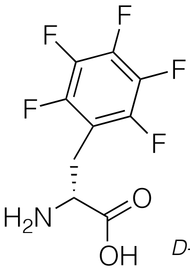 *D-Phe(5F)*

143 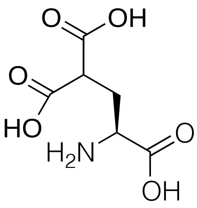 *L*-Gla

144 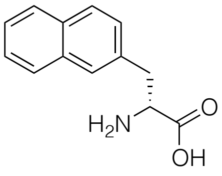 *D*-2-Nal

145 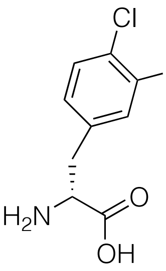 *D-Phe(3,4-Cl$_2$)*

146 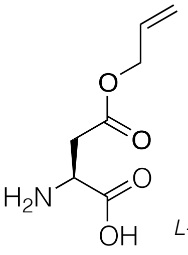 *L-Asp(OAll)*

147 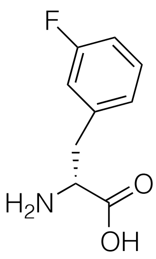 *D-Phe(3-F)*

148 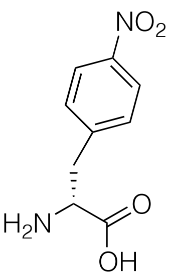 *D-Phe(4-NO$_2$)*

149 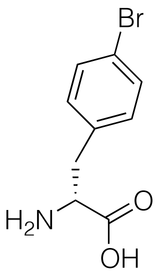 *D-Phe(4-Br)*

150 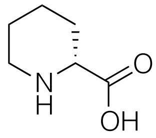 D-Pip

151 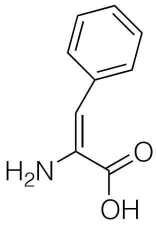 *dhPhe*

152 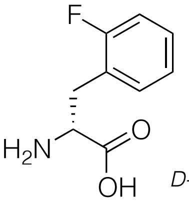 *D-Phe(2-F)*

153 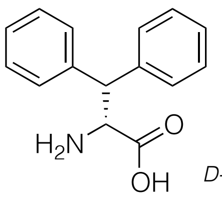 *D-Dip*

154 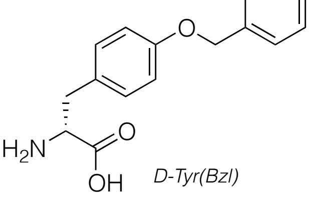 *D-Tyr(Bzl)*

155 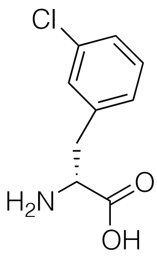 *D-Phe(3-Cl)*

156
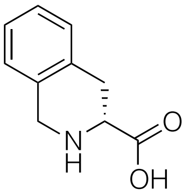
D-Tic

157
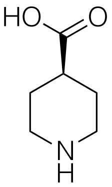
*L*-Inp

158
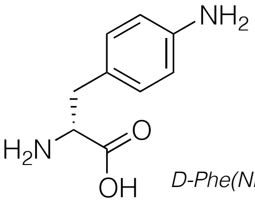
*D-Phe($NH_2$)*

157
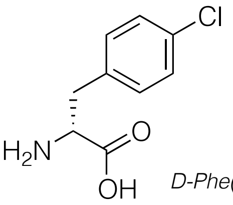
*D-Phe(4-Cl)*

158
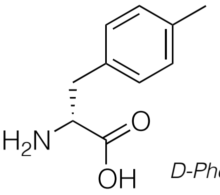
*D-Phe(4-Me)*

159
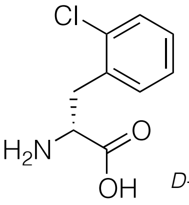
*D-Phe(2-Cl)*

158
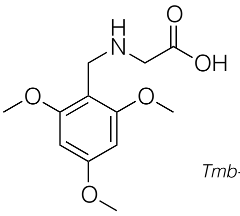
*Tmb-Gly-OH*

159
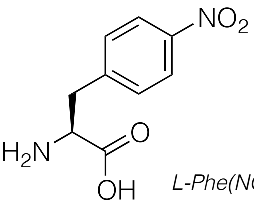
*L-Phe($NO_2$)*

160
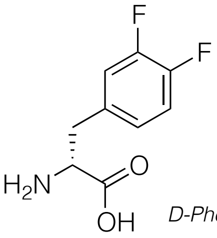
*D-Phe(3,4-$F_2$)*

161
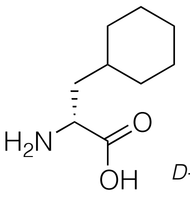
*D-Cha*

162
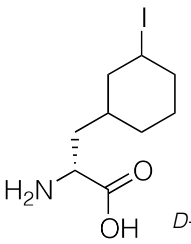
*D-Phe(3-I)*

163
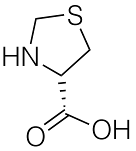
D-Thz

164
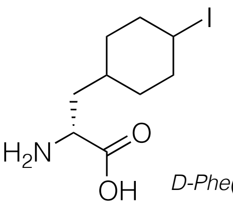
*D-Phe(4-I)*

165
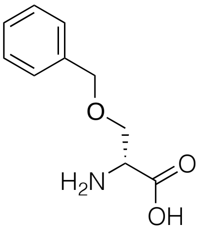
*D-Ser(Bzl)*

166
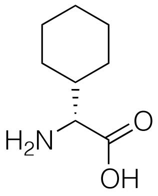
*D-Chg*

165 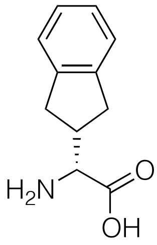 *D-Igl*

166 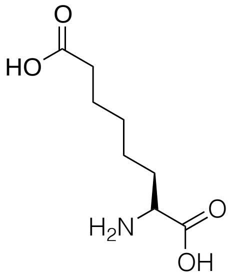 *L-Asu*

167 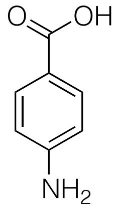 *4-Abz*

168 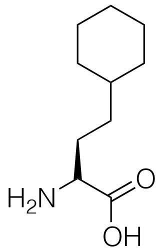 *L-hCha*

169 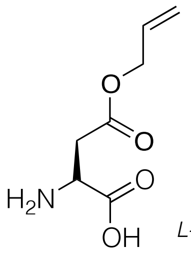 *L-Asp(OAll)*

170 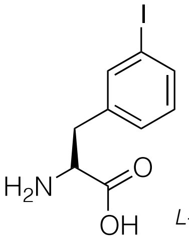 *L-Phe(3-I)*

171 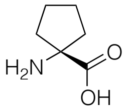

*L*-1-amino-cyclopentane carboxylic acid

172 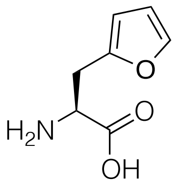 *L-2-Fal*

173 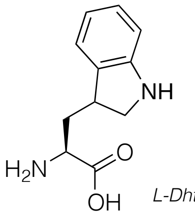 *L-Dht*

174 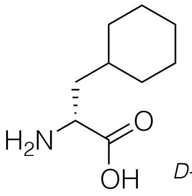 *D-Cha*

**Figure S2**. Structures of amino acids used in P1 library with general structure Ac-P5-X-Glu-X-Asp-ACC.

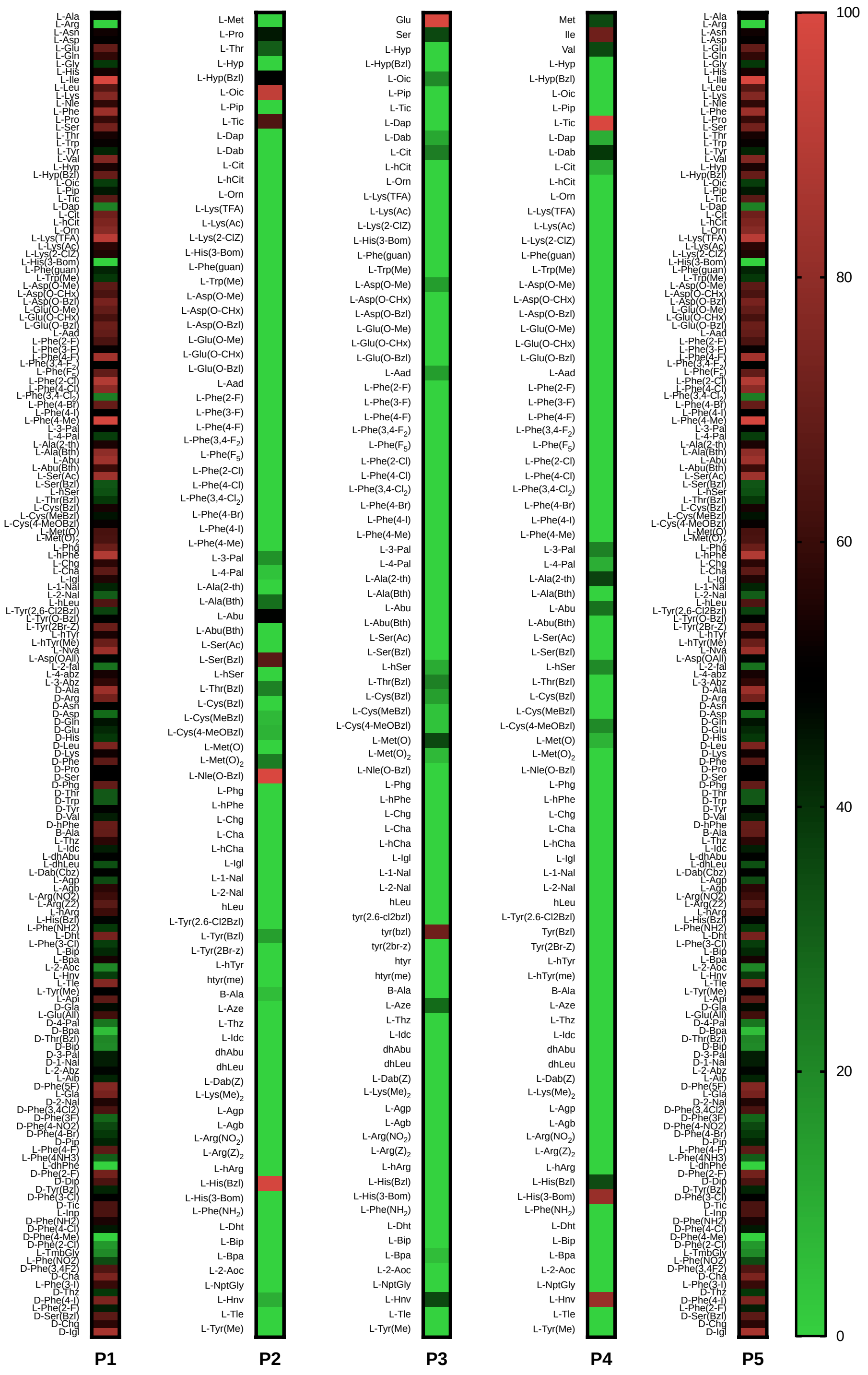


**Fiure S3.** GrB catalytic preferences in nonprime enzyme pockets. General structure for P1 library was: Ac-Ile-Ser-Pro-P1-ACC, for P2-P4 was Ac-X-X-X-Asp-ACC and for P5 was Ac-X-Glu-X-Asp-ACC, where “P1” and “P5” were defined amino acids and “X” was isokinetic mixture.

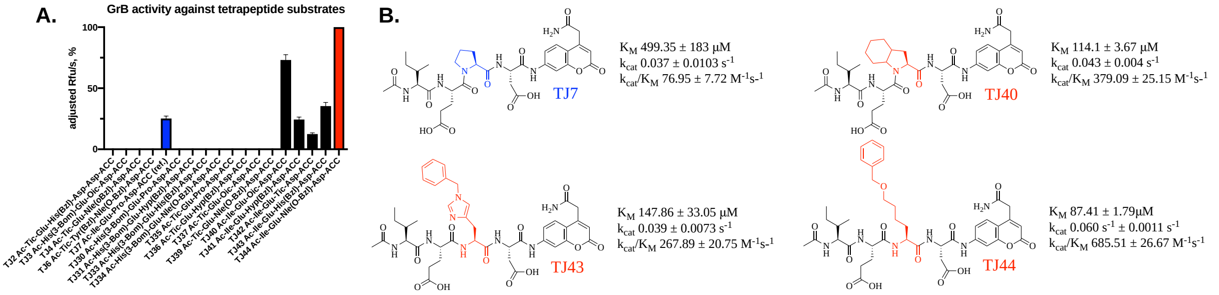


***Figure S4 A) GrB activity against tetrapeptide substrates.*** *GrB activity was tested against substrates TJ2-7, TJ30-44 (an equal concentration in an assay buffer). Results are presented in a column diagram as adjusted percentage values of RFU/s. The bar for the best substrate was labeled red and reference one blue.* ***B) Selected tetrapeptide substrates detailed kinetic analysis.*** *$k_{cat}/K_M$ value calculation was analyzed using GraphPad Prism software. Data are depicted as mean ± standard deviation and represent at least 2 independent experiments.*

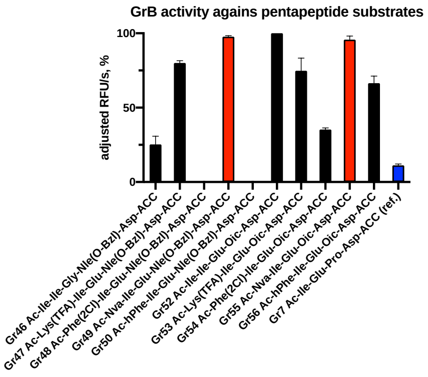


***Figure S5 GrB activity against pentapeptide substrates.*** *GrB activity was tested against substrates Tj46-56 (an equal concentration in an assay buffer). Results are presented in a column diagram as adjusted percentage values of RFU/s. The bar for the best substratse was labeled red and reference one blue*

A)

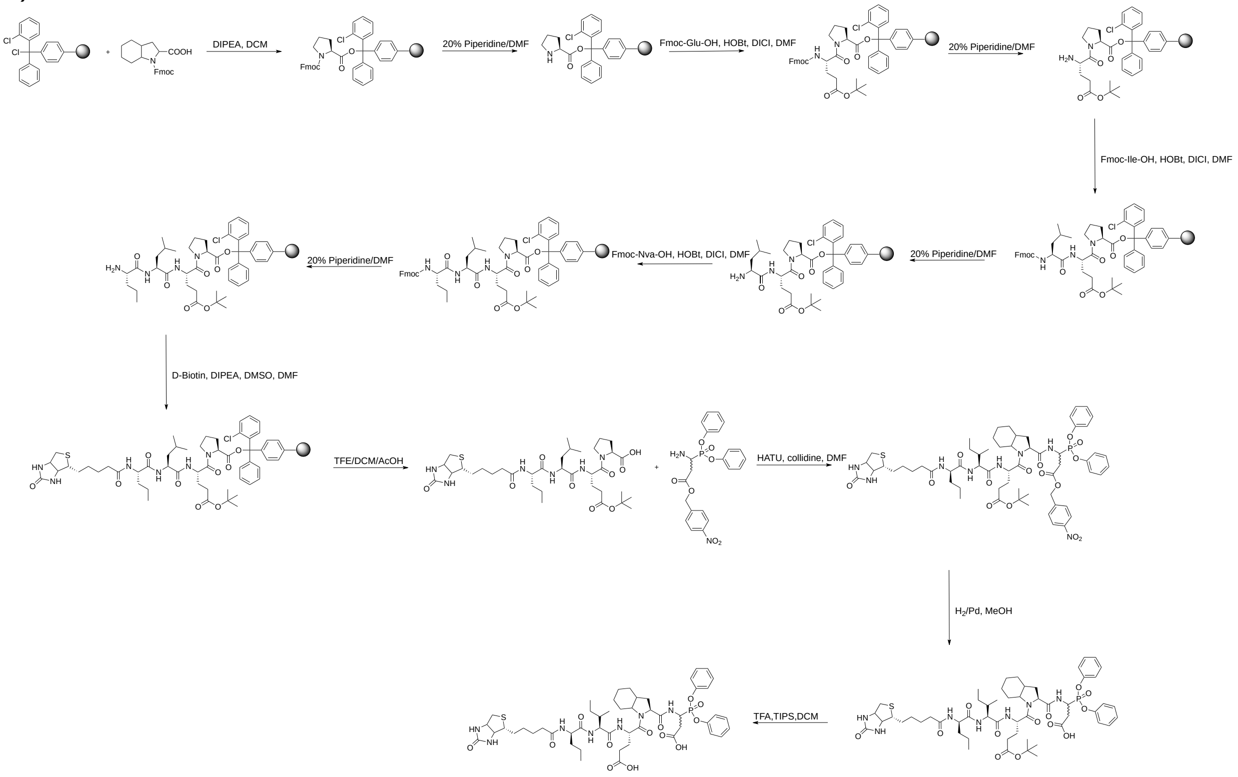


B)

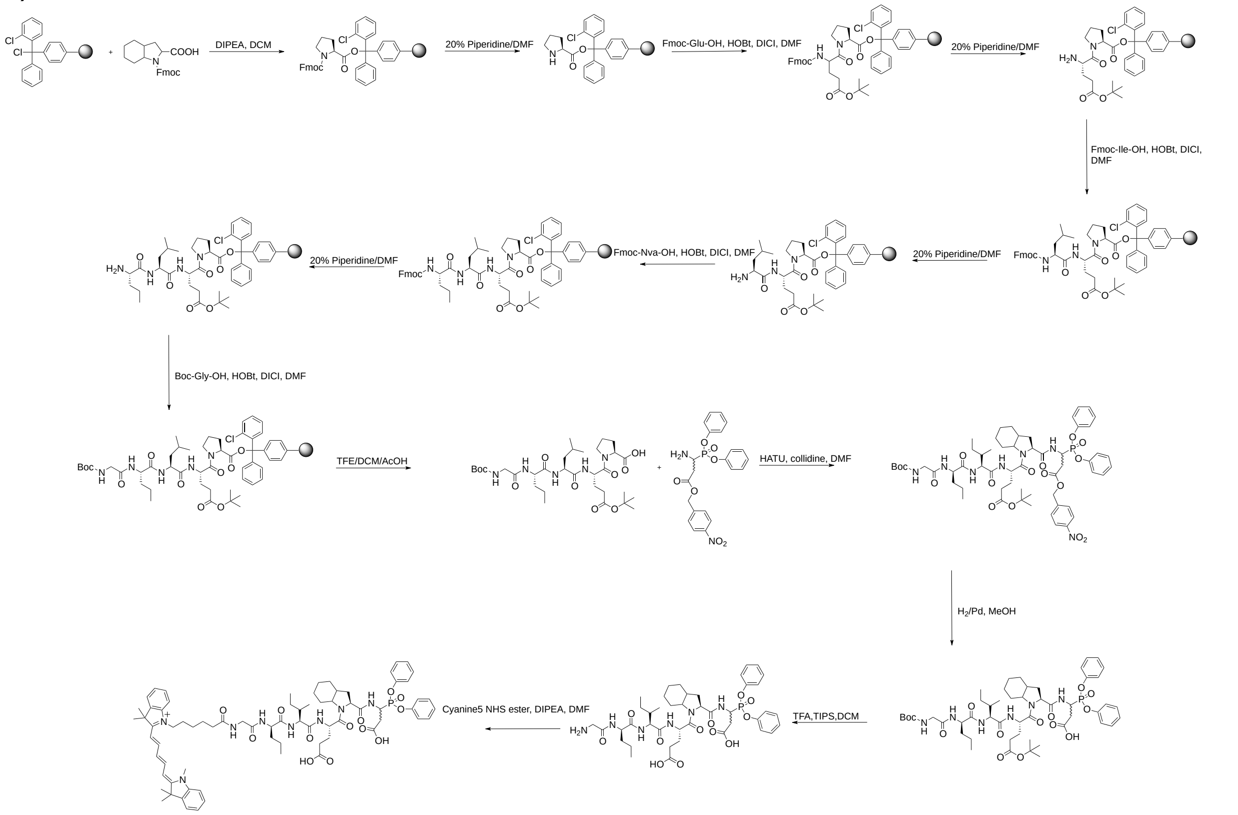


**Figure S6**. A) TJ55.Bt synthesis scheme B) TJ55.5 synthesis scheme. The probes were synthesized using a mix of classic solid phase and in solution peptide synthesis methods.

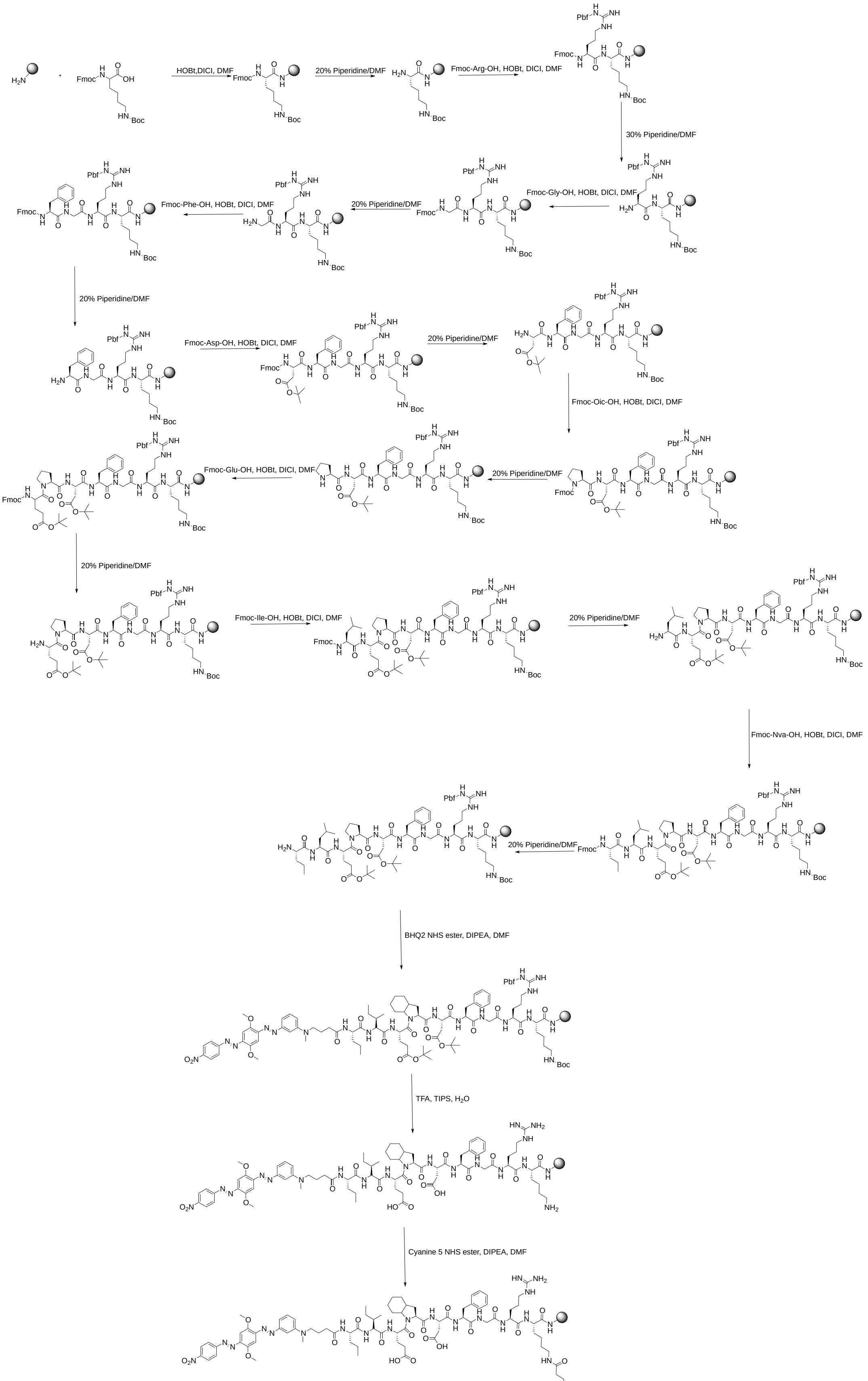


**Figure S7** qTJ71 synthesis scheme. The quenched fluorescent substrate was synthesized using a mix of classic solid phase and in solution peptide synthesis methods.

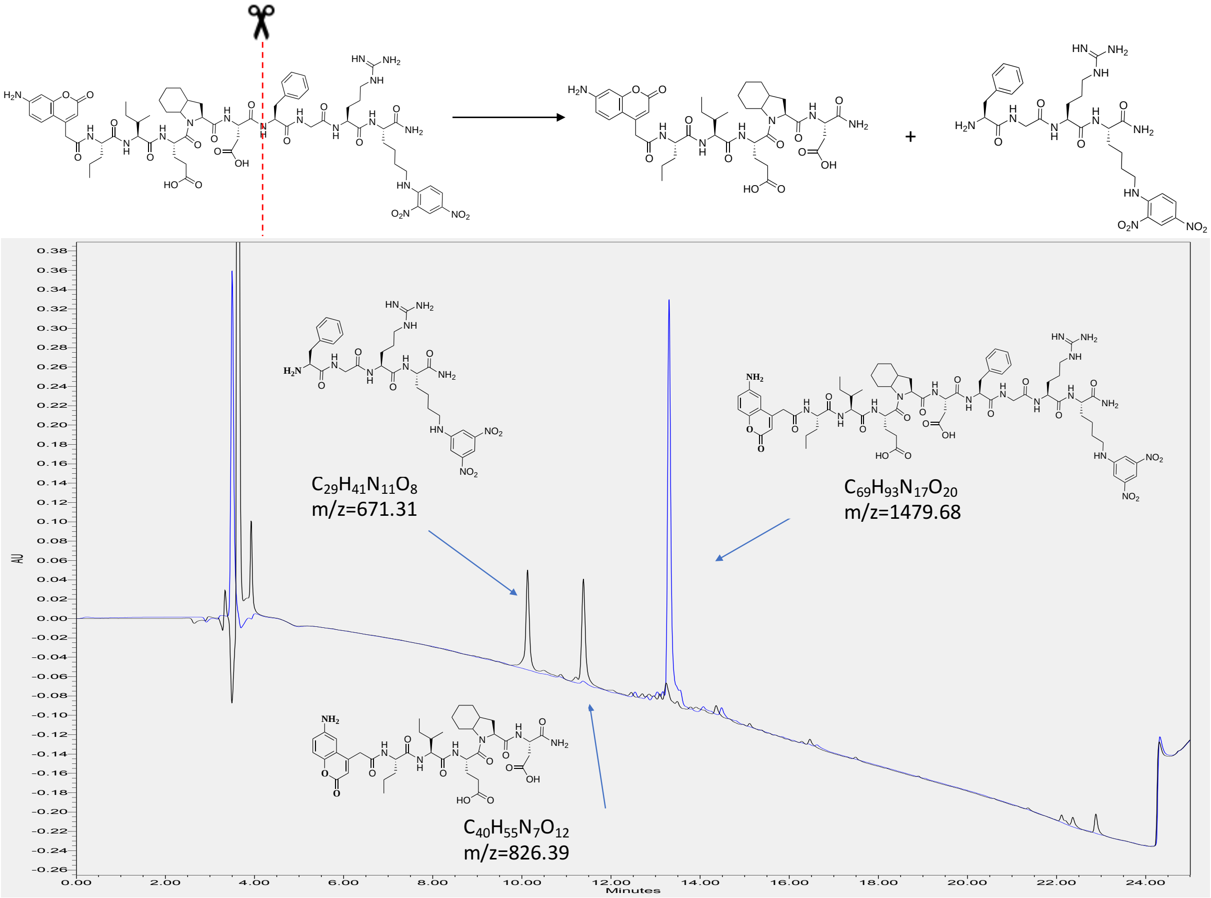


**Figure S8.** Cleavage of Gr71 by GrB. Chromatogram from LCMS showing substrate and products after cleavage. Blue line shows substrate before reaction, black after cleavage. The peak at the 3rd minute is from buffer components. While designing IQF substrates, there was expected cleavage of the peptide at one specific site, after the aspartic acid. To test this hypothesis, IQF substrate (50μM) was treated with the granzyme B (43nM) and monitored the hydrolytic yield with analytical HPLC at two separate time points. After 30 minutes of hydrolysis, two new peaks appeared, and the signal from the intact substrate decreased over time, while the signal from two new products increased over time (up to 30 minutes). Therefore, it was confirmed that granzyme B hydrolyzes the new IQF substrate at one site only. Thus, there was performed mass spectrometry of the reaction mixture. As expected, the mass spectrum demonstrated that cleavage occurred between the P1 and P1' amino acid residues. There were no any other peaks that could indicate a different peptide bond hydrolysis site. After hydrolysis, the fluorescence is no longer quenched, and the intensity of the signal increases, making these substrates valuable tools for granzyme B activity measurements in extracts or cell lysates.

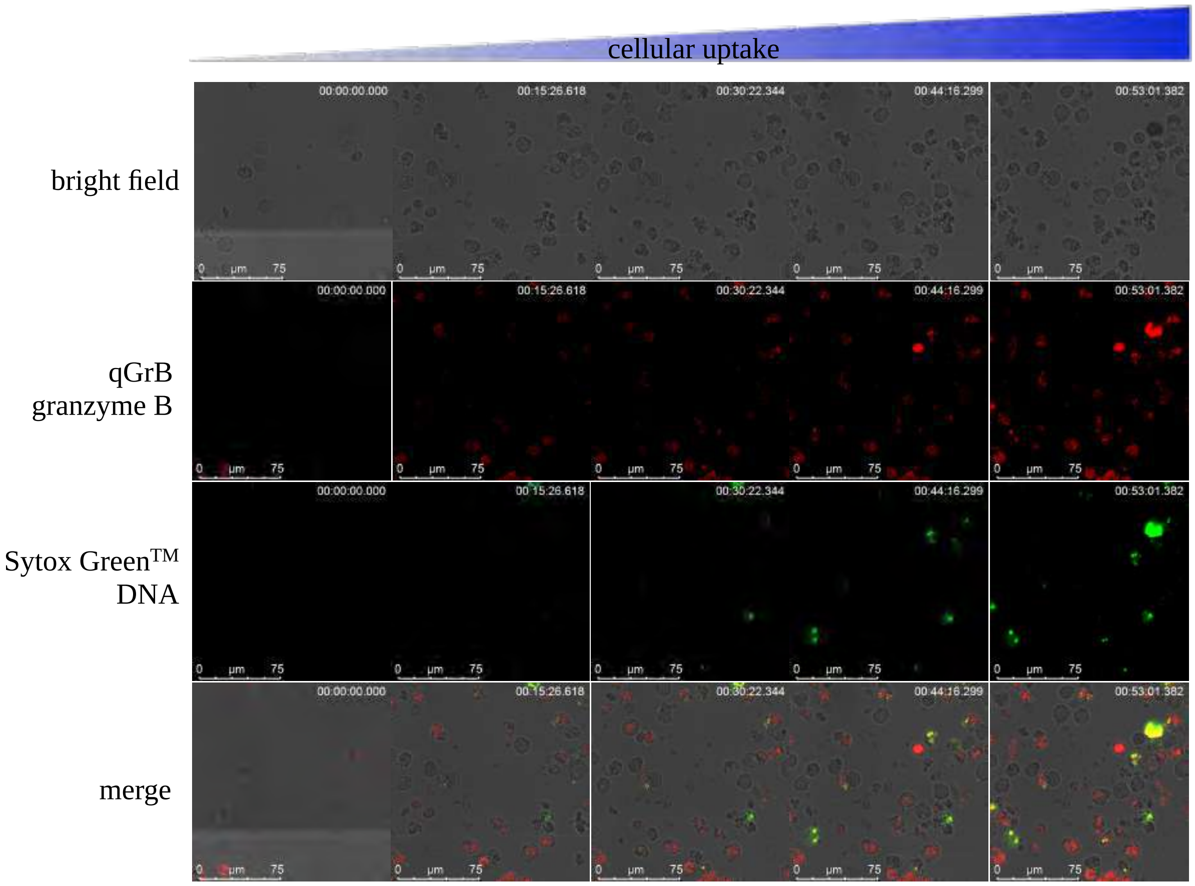


**Figure S9. qTJ71 cellular uptake.** Experiment was performed using Leica TCS SP8 confocal microscope. YT cells ($1 \times 10^5$ cells/ml) in YT cell culture media (RPMI1640 media supplemented with 10% of Fetal Bovine Serum and 1% Penicillin/Streptomycin) were treated with 1µM of qTJ71, 1µM of Sytox Green™ for 1 minute. The experiment was carried out for a 1 hour using laser wavelengths of: 488 nm for Brightfield and Sytox™ Green [S7020]; 552 nm for Cyanine 3. Imaging was carried out in the chamber [(the box life imaging services] in the presence of 5% $CO_2$.

### Substrates and probes structutes, MS and HPLC analysis

**TJ2 Ac-Tic-Glu-His(Bzl)-Asp-ACC**

HRMS (m/z) [$MH^+$] calcd. for $C_{45}H_{46}N_8O_{12}$, 890.32, found, 891.32

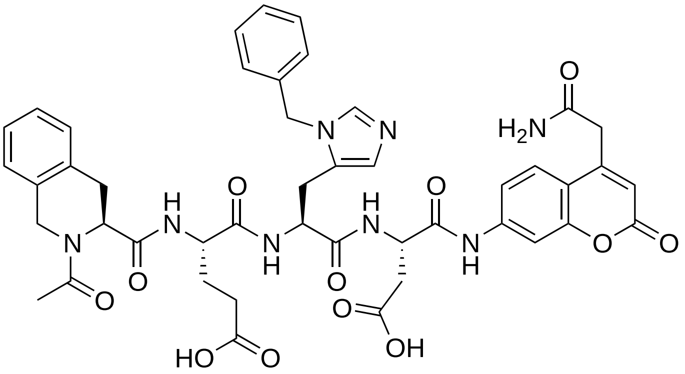


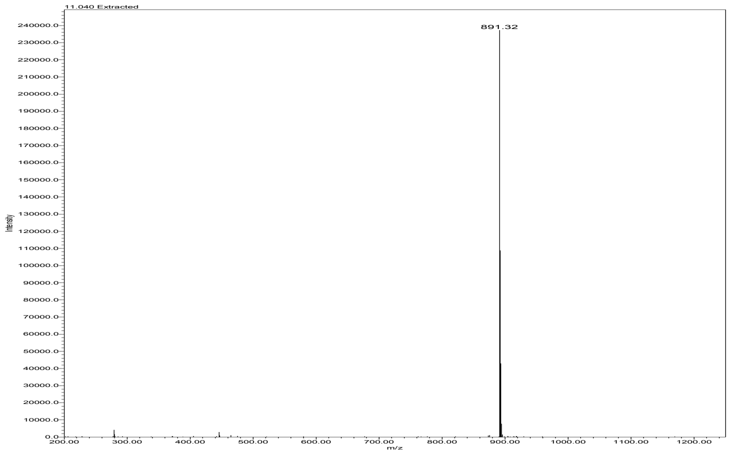


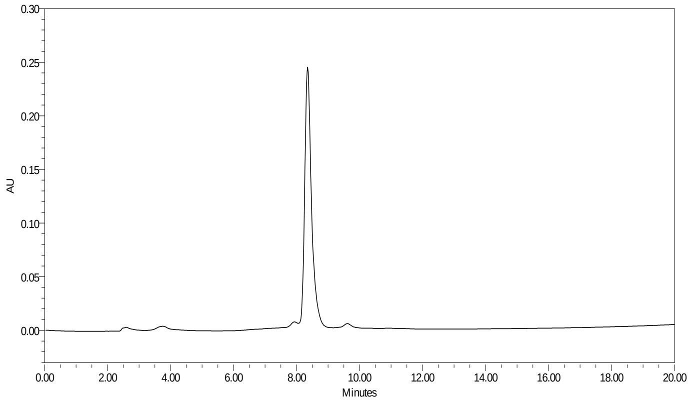

**TJ3 Ac-His(3-Bom)-Glu-Oic-Asp-ACC**

HRMS (m/z) [MH$^{+}$] calcd. for $C_{45}H_{52}N_{8}O_{13}$, 912.37, found, 913.37

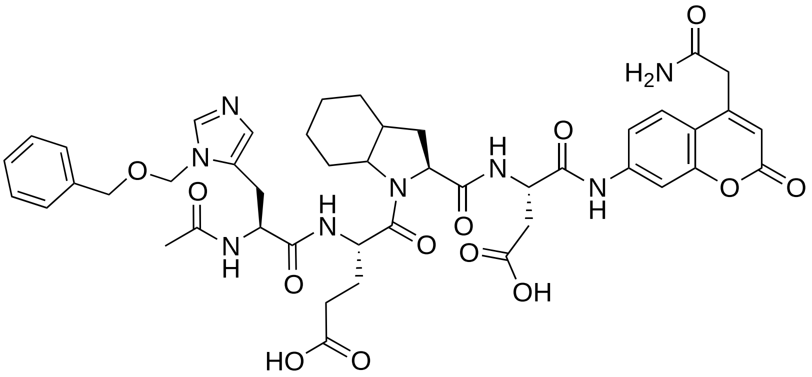

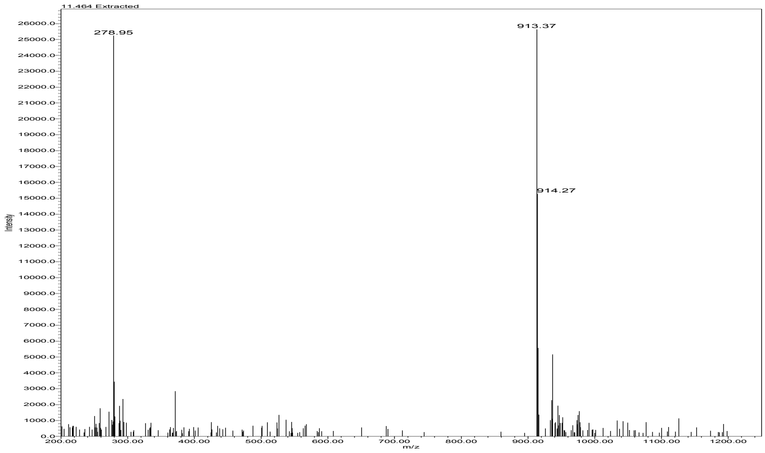


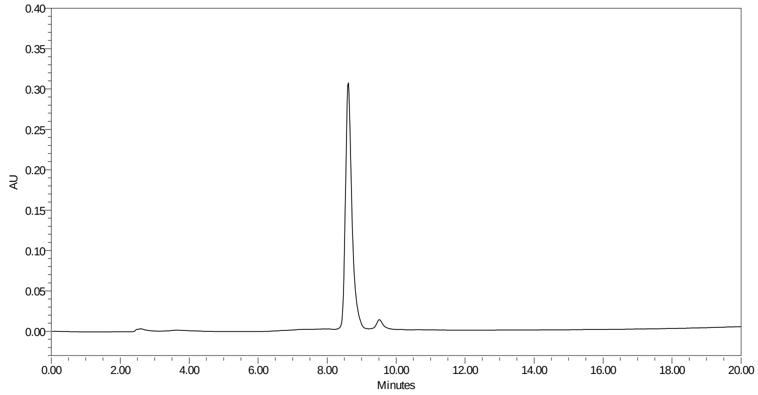

**TJ4 Ac-Tic-Glu-Nle(O-Bzl)-Asp-ACC**

HRMS (m/z) [MH$^+$] calcd. for $C_{45}H_{50}N_6O_{13}$, 882.91, found, 883.33

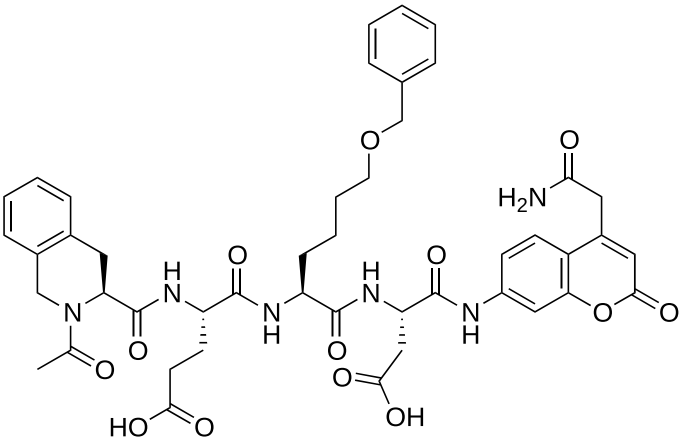

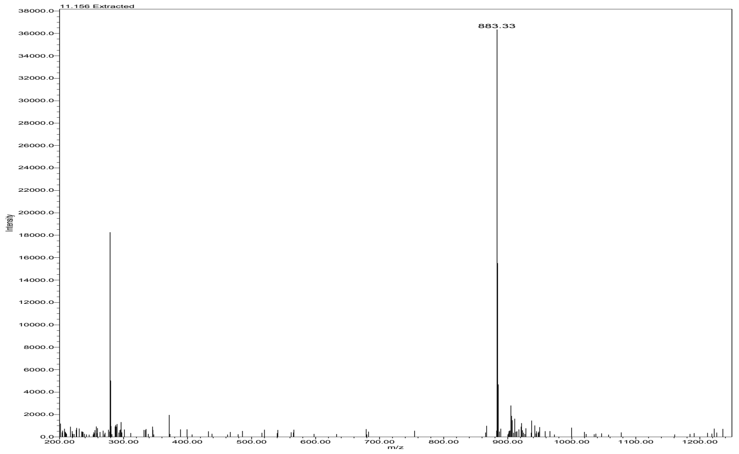


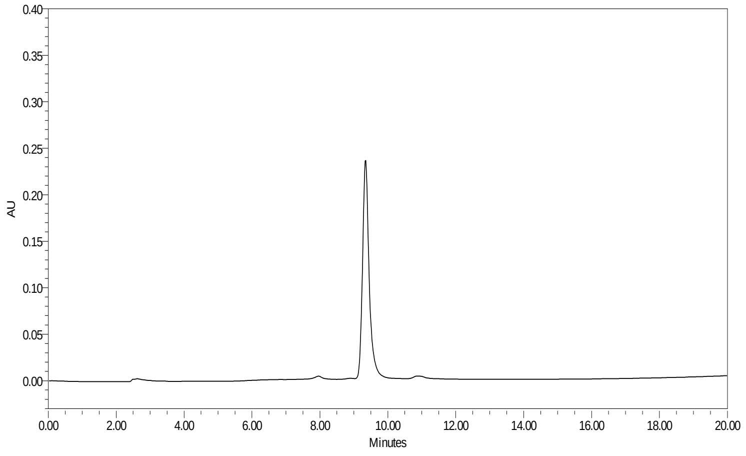

**TJ6 Ac-Tic-Tyr(Bzl)-Nle(O-Bzl)-Asp-ACC**

HRMS (m/z) [MH$^+$] calcd. for $C_{56}H_{58}N_6O_{12}$, 1006.41, found, 1007.44

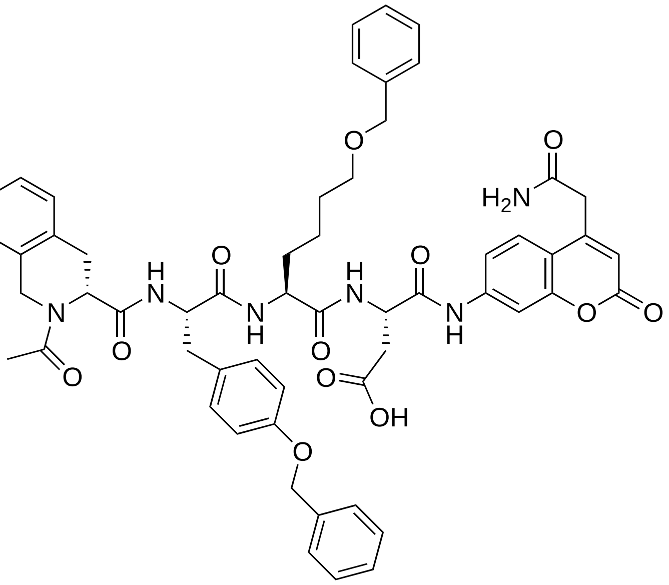


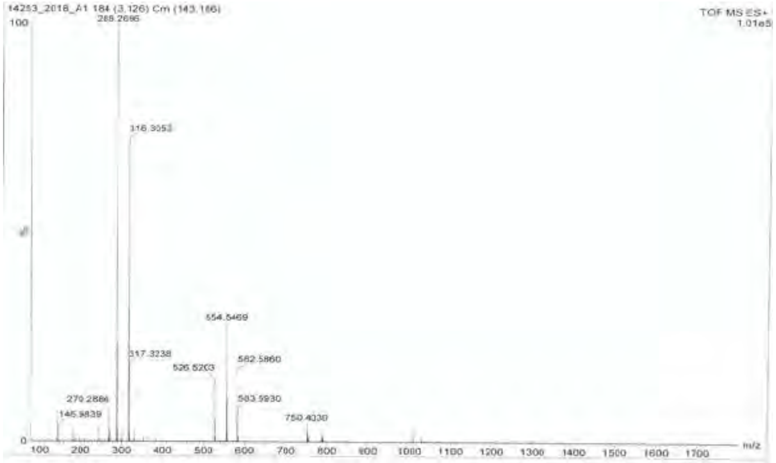


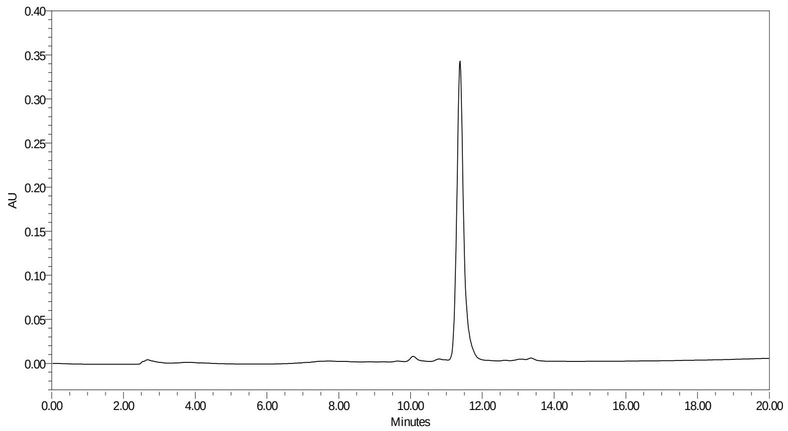

**TJ7 Ac-Ile-Glu-Pro-Asp-ACC**

HRMS (m/z) [$MH^+$] calcd. for $C_{33}H_{42}N_6O_{12}$, 714.29, found, 715.17

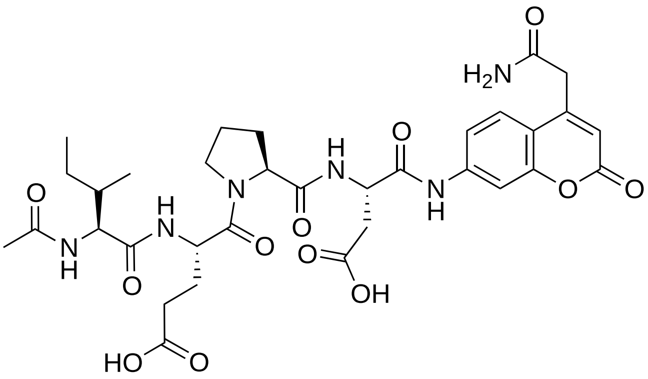

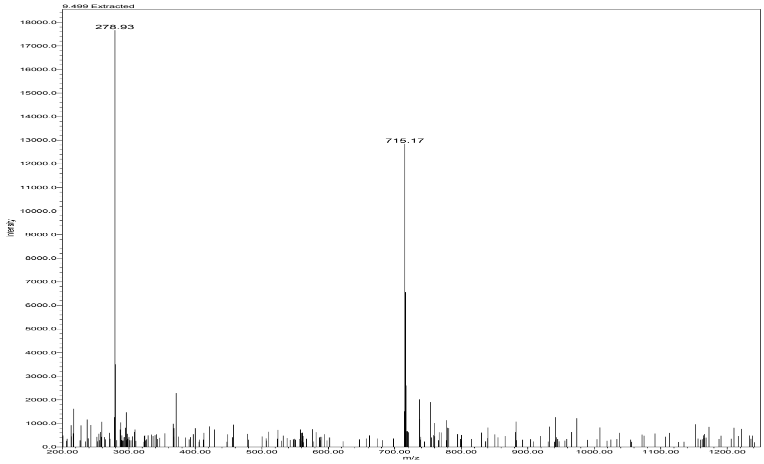


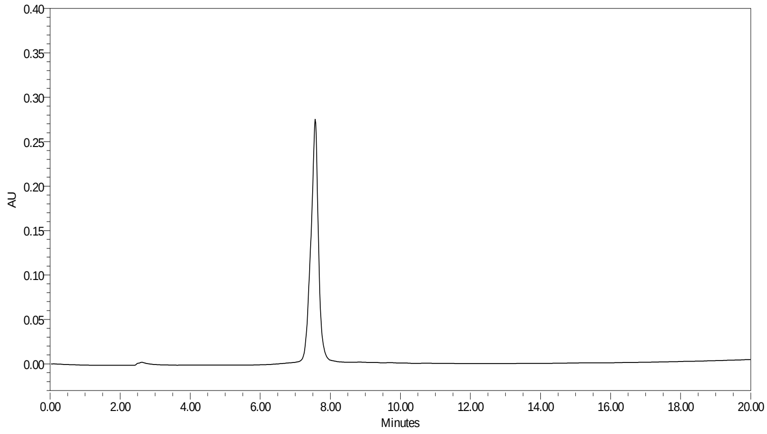

**TJ30 Ac-His(3-Bom)-Glu-Pro-Asp-ACC**

HRMS (m/z) [$MH^+$] calcd. for $C_{41}H_{46}N_8O_{13}$, 858.32, found, 859.33

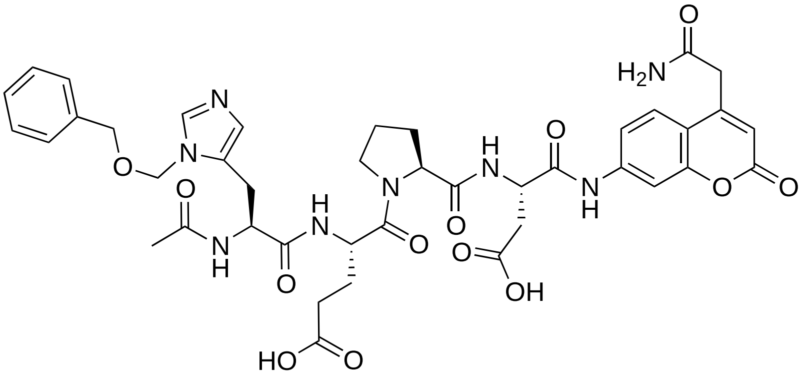

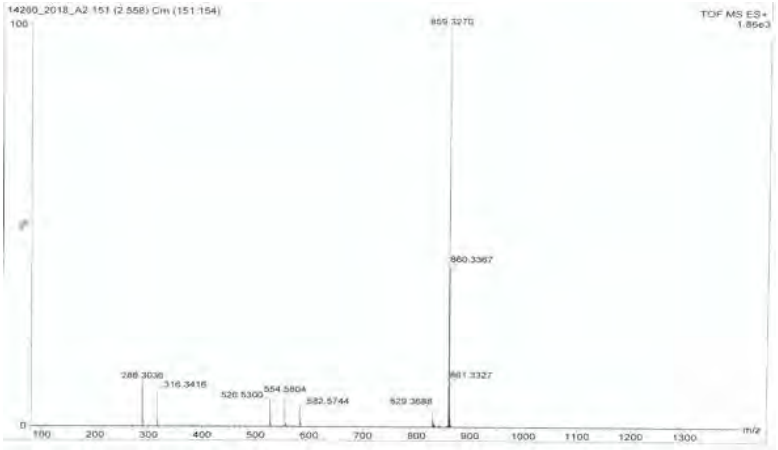


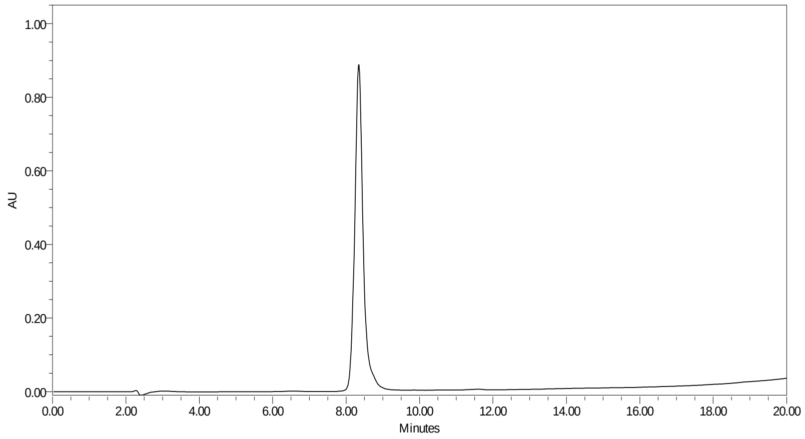

**TJ31 Ac-His(3-Bom)-Glu-Hyp(Bzl)-Asp-ACC**

HRMS (m/z) [MH$^+$] calcd. for $C_{48}H_{52}N_8O_{14}$, 964.36, found, 965.37

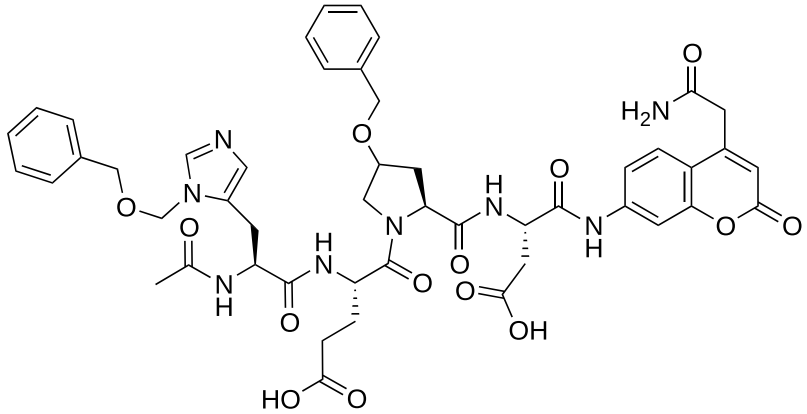

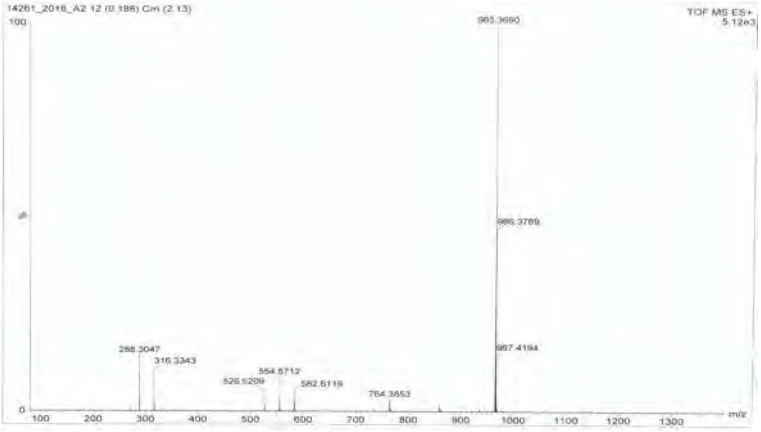


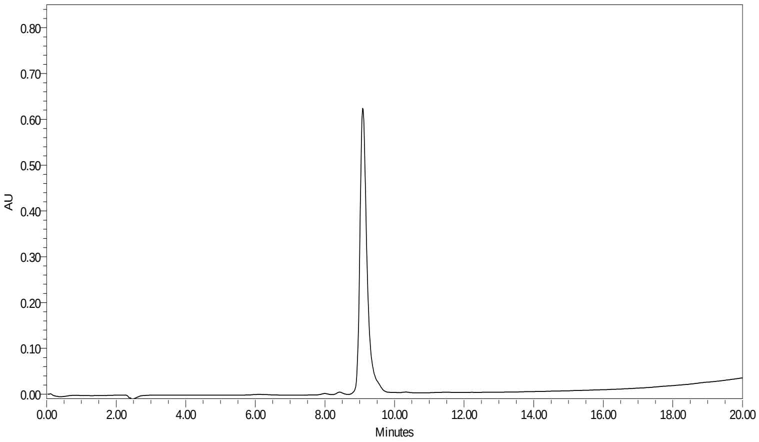

**TJ33 Ac-His(3-Bom)-Glu-His(Bzl)-Asp-ACC**

HRMS (m/z) [MH$^+$] calcd. for $C_{49}H_{52}N_{10}O_{13}$, 988.37, found, 989.39

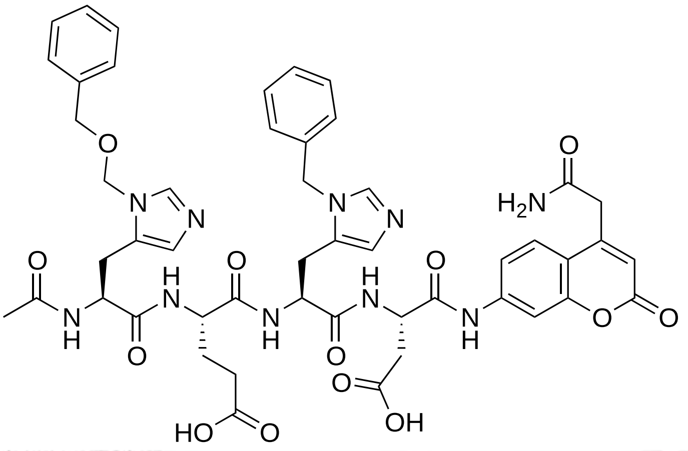

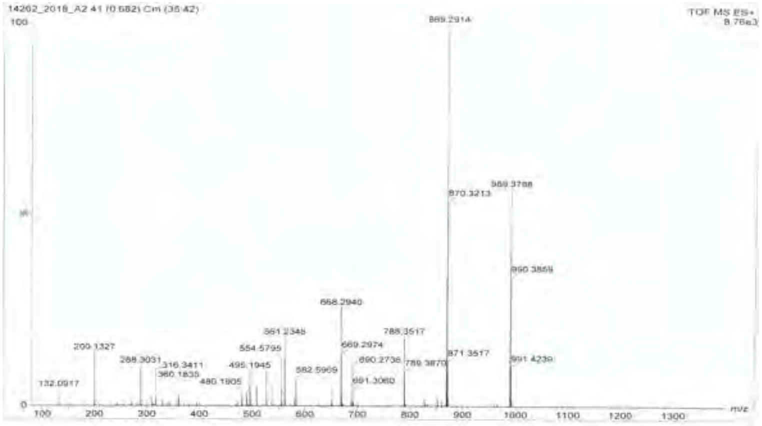


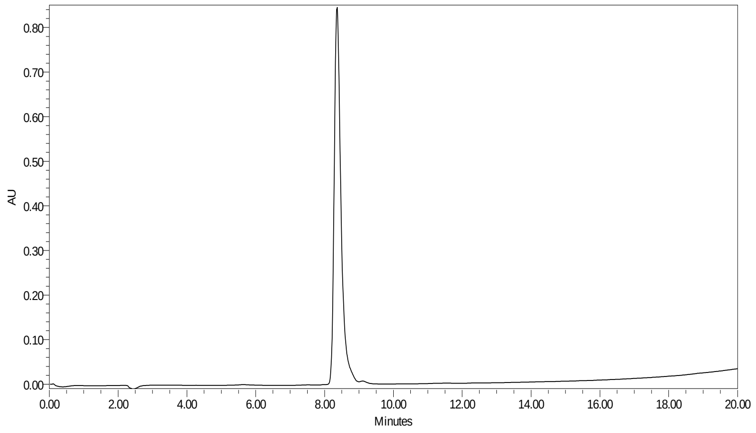

**TJ34 Ac-His(3-Bom)-Glu-Nle(OBzl)-Asp-ACC**

HRMS (m/z) [MH$^+$] calcd. for $C_{49}H_{56}N_8O_{14}$, 980.39, found, 981.40

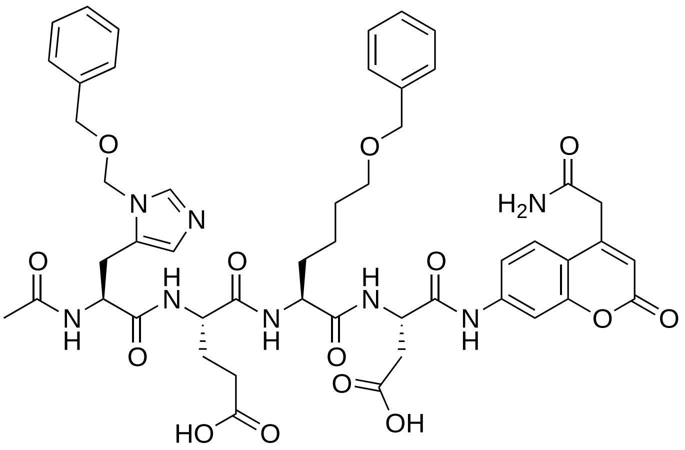

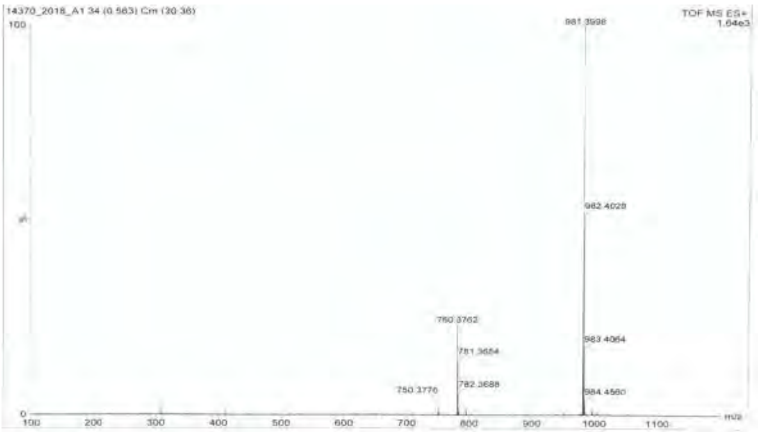


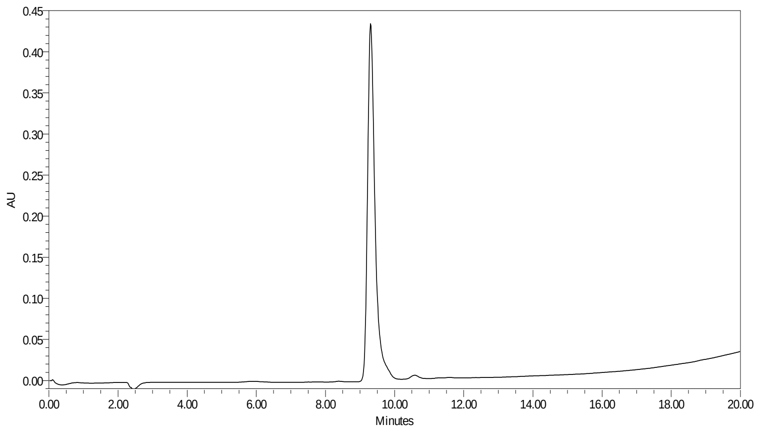

**TJ35 Ac-Tic-Glu-Pro-Asp-ACC**

HRMS (m/z) [MH$^+$] calcd. for $C_{37}H_{40}N_6O_{12}$, 760.27, found, 783.26

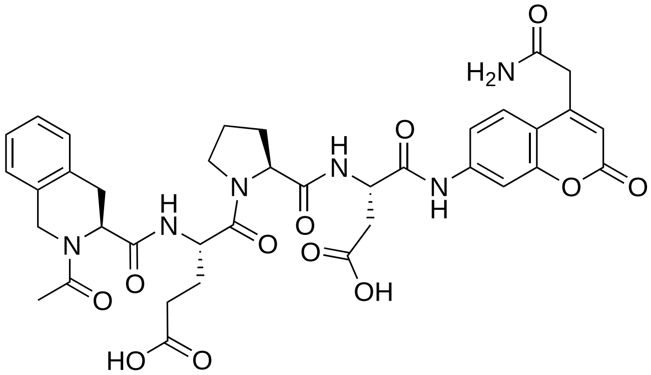

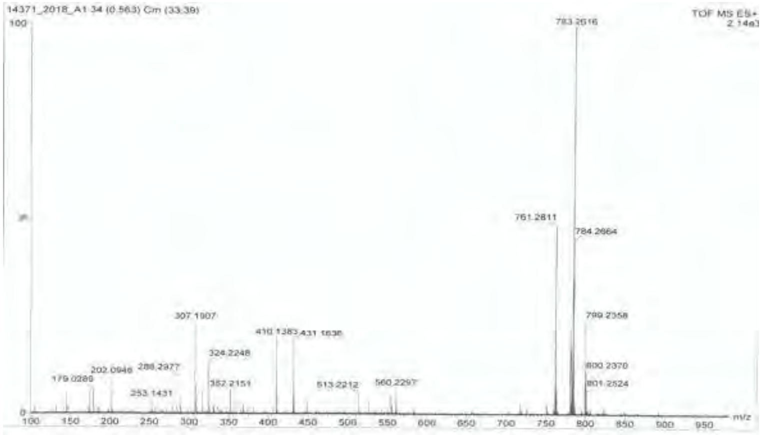


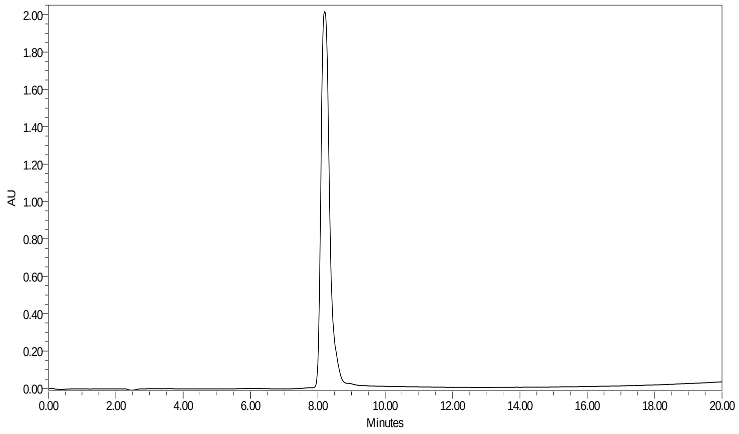

**TJ36 Ac-Tic-Glu-Hyp(Bzl)-Asp-ACC**

HRMS (m/z) [MH$^+$] calcd. for $C_{44}H_{46}N_6O_{13}$, 866.31, found, 867.32

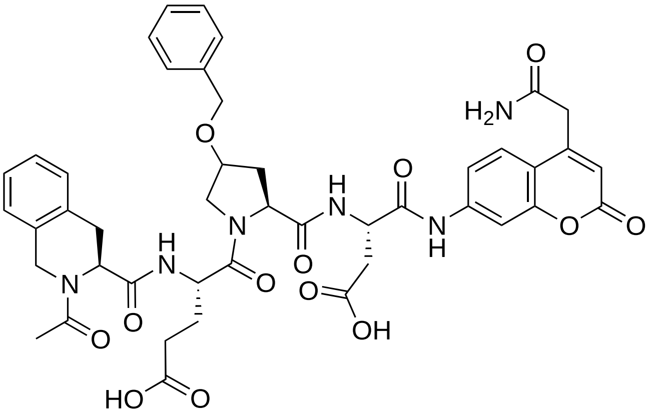

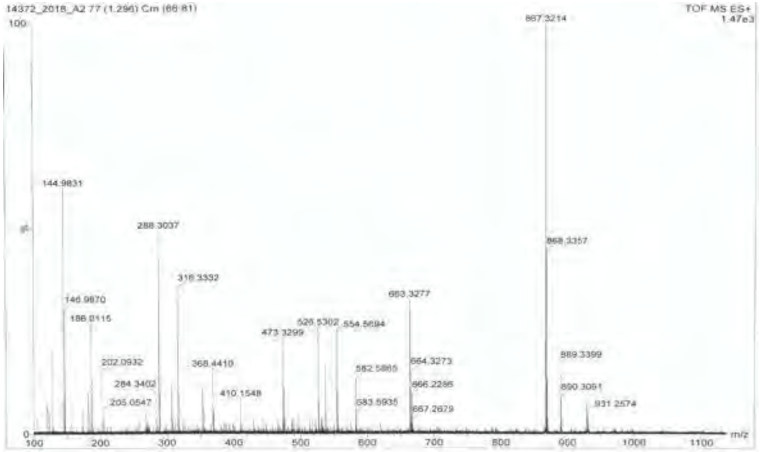


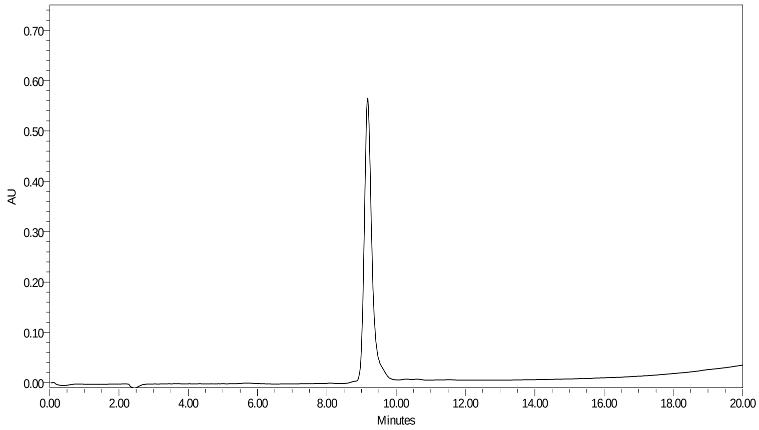

**TJ37 Ac-Tic-Glu-Oic-Asp-ACC**

HRMS (m/z) [MH$^+$] calcd. for $C_{41}H_{46}N_6O_{12}$, 814.32, found, 815.33

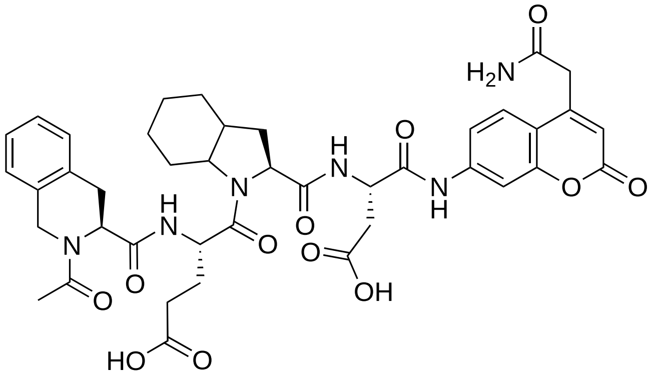

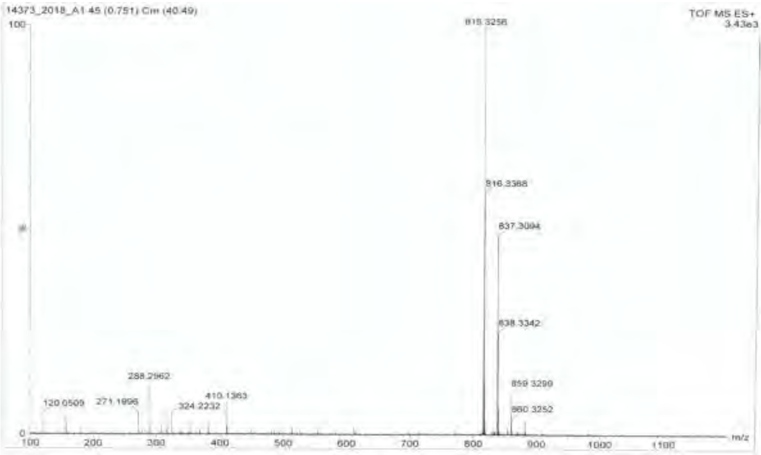


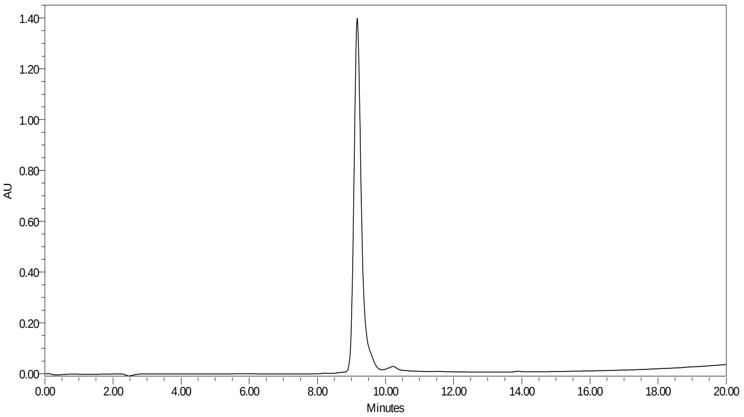

**TJ39 Ac-Tic-Glu-Nle(O-Bzl)-Asp-ACC**

HRMS (m/z) [MH$^+$] calcd. for $C_{45}H_{50}N_6O_{13}$, 882.91, found,905.33

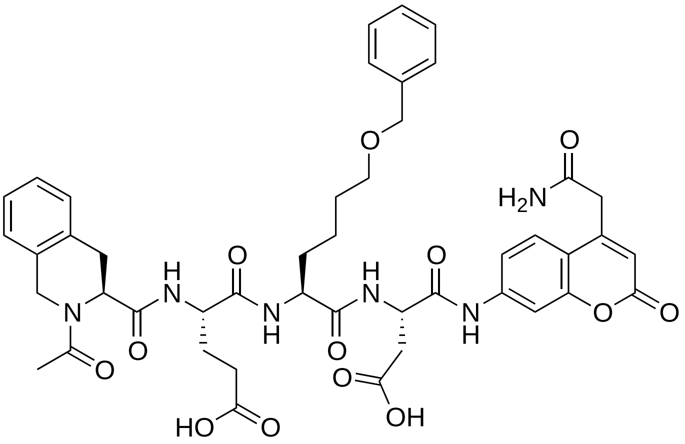

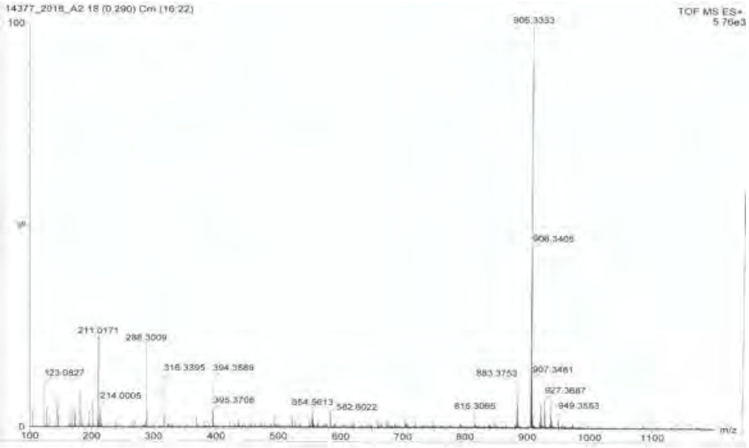


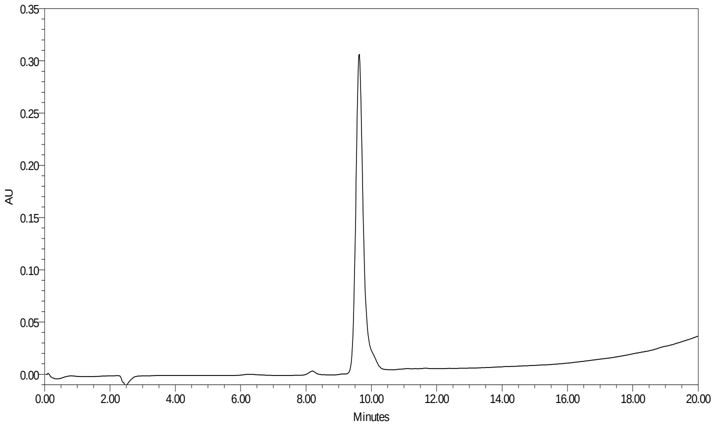

**TJ40 Ac-Ile-Glu-Oic-Asp-ACC**

HRMS (m/z) [MH$^+$] calcd. for $C_{37}H_{48}N_6O_{12}$, 768.33, found,769.35

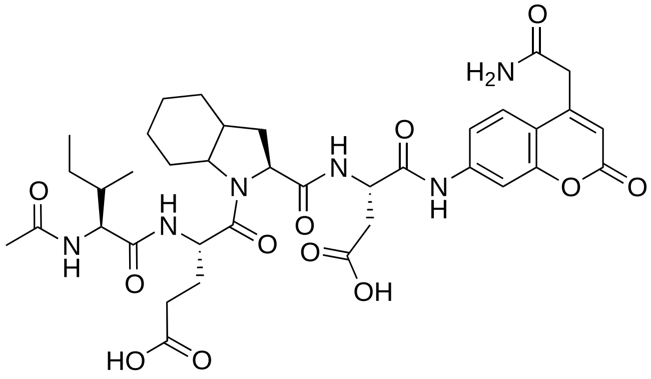

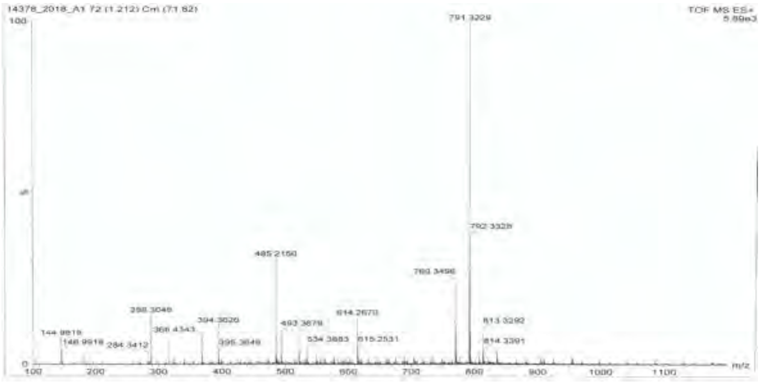

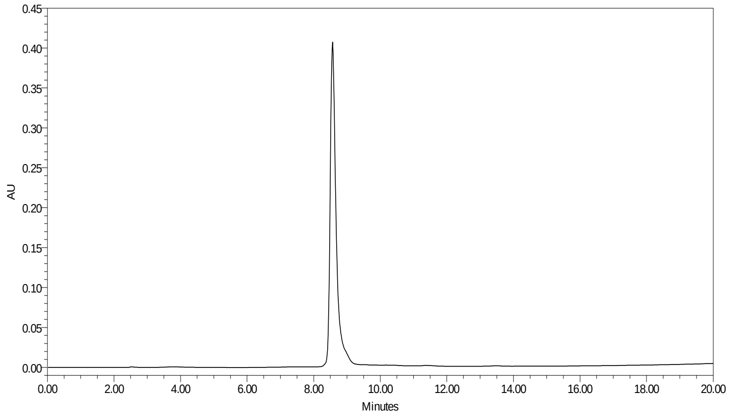

**TJ41 Ac-Ile-Glu-Hyp(Bzl)-Asp-ACC**

HRMS (m/z) [MH$^+$] calcd. for $C_{40}H_{48}N_6O_{13}$, 820.33, found, 821.21

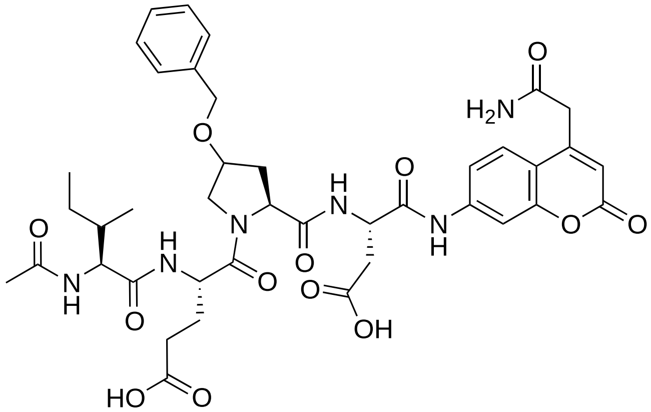

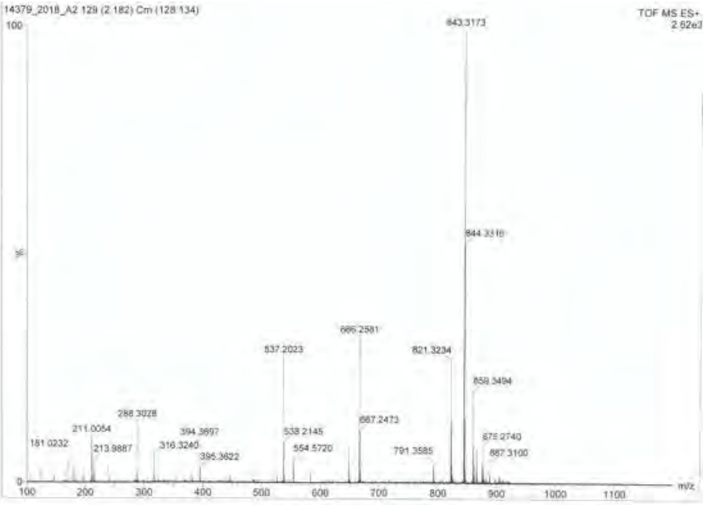


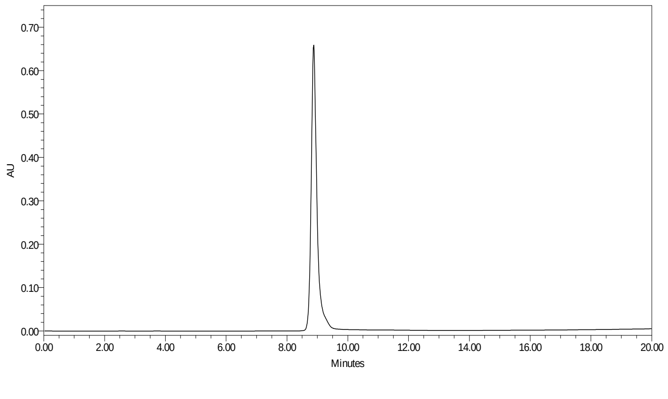

**TJ42 Ac-Ile-Glu-Tic-Asp-ACC**

HRMS (m/z) [$MH^+$] calcd. for $C_{38}H_{44}N_6O_{12}$, 776.30, found, 771.33

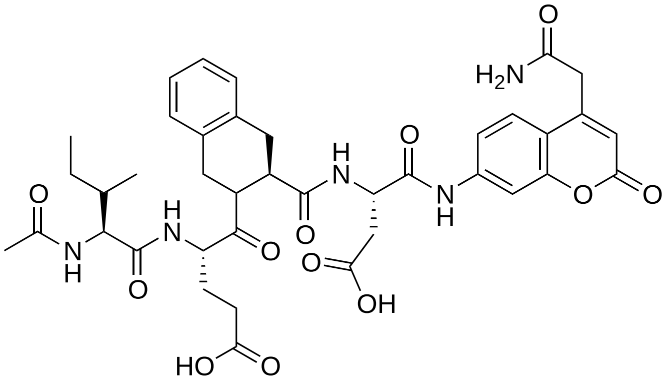

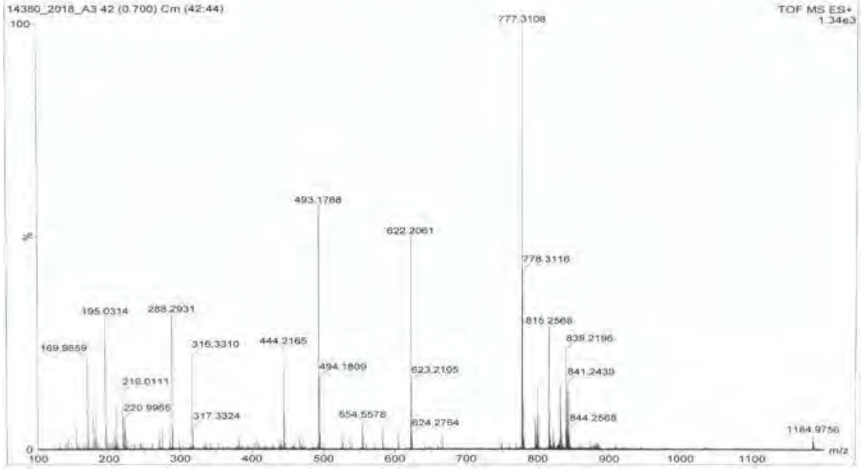


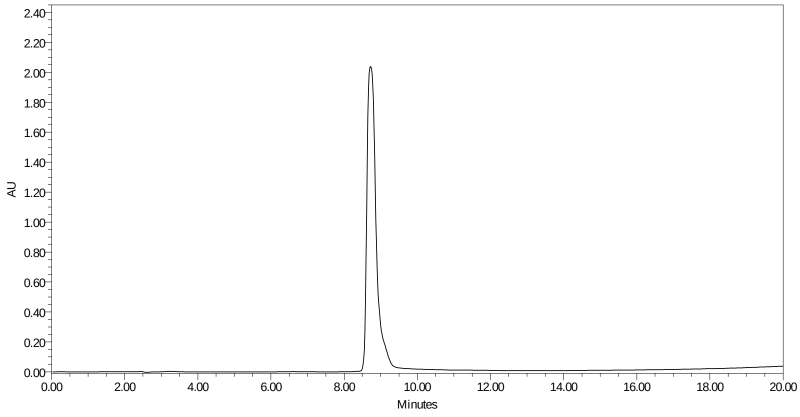

**TJ43 Ac-Ile-Glu-His(Bzl)-Asp-ACC**

HRMS (m/z) [MH$^+$] calcd. for $C_{41}H_{48}N_8O_{12}$, 844.34, found, 845.35

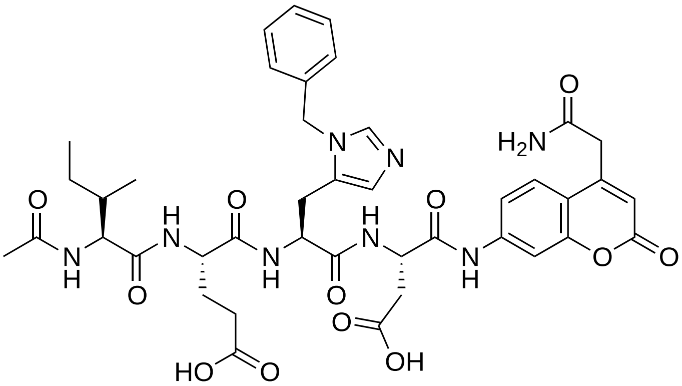


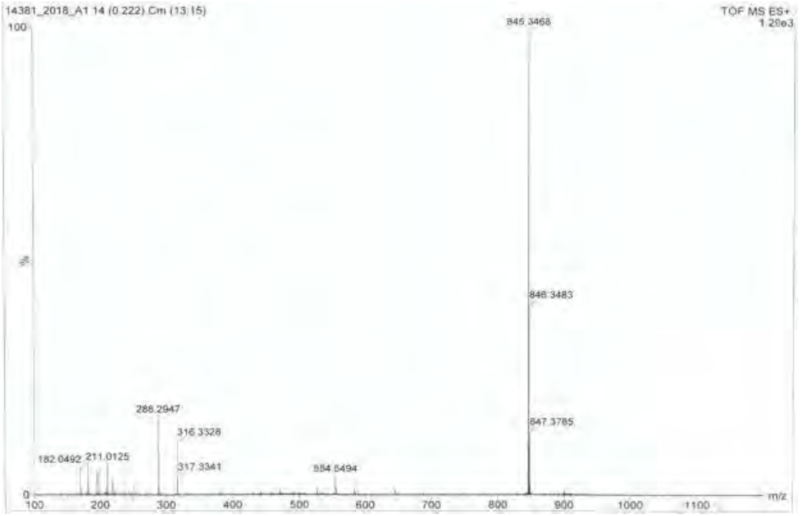


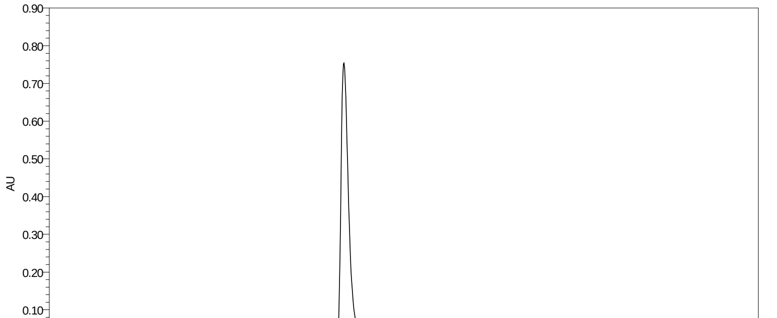

**TJ46 Ac-Ile-Ile-Glu-Nle(O-Bzl)-Asp-ACC**

HRMS (m/z) [$MH^+$] calcd. for $C_{47}H_{63}N_7O_{14}$, 949.44, found, 948.43

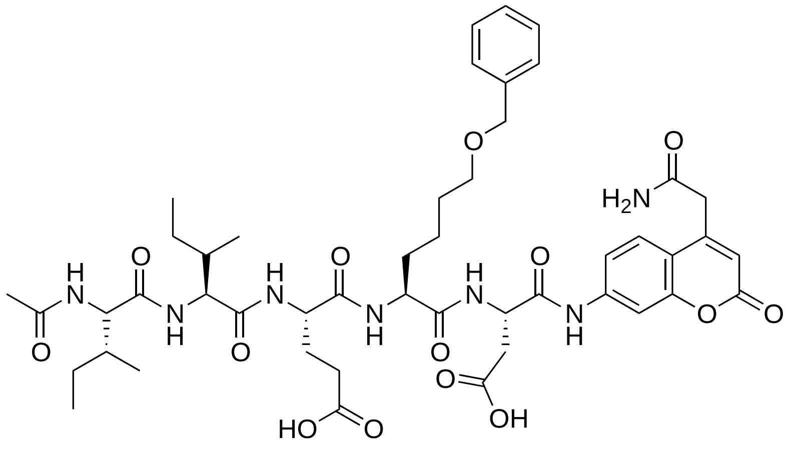

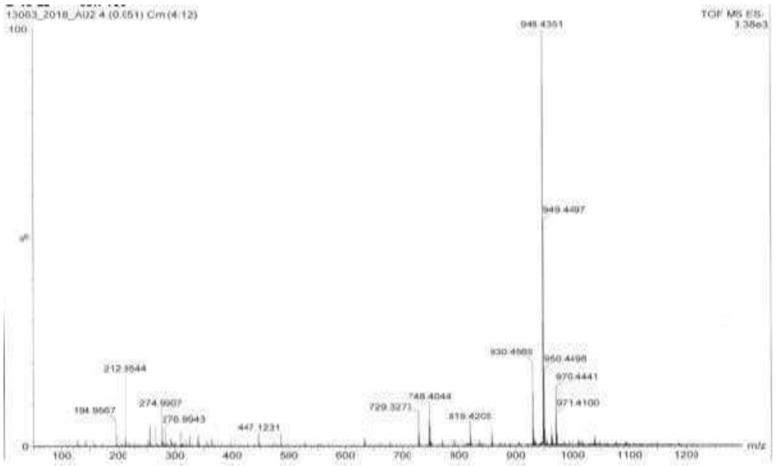

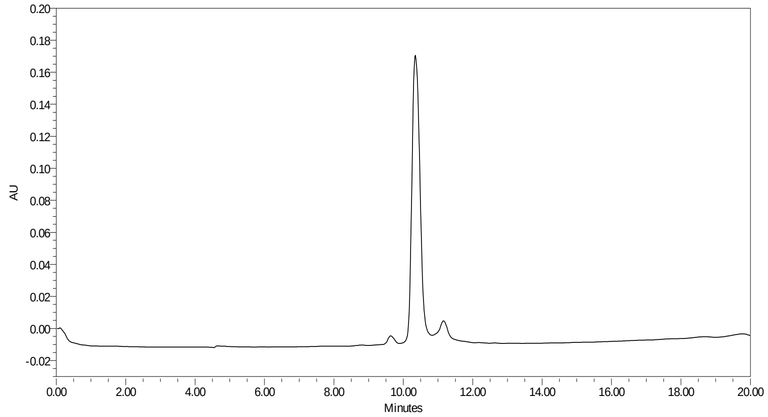

**TJ47 Ac-Lys(TFA)-Ile-Glu-Nle(O-Bzl)-Asp-ACC**

HRMS (m/z) [MH$^+$] calcd. for $C_{49}H_{63}FN_8O_{15}$, 1060.44, found, 1061.44

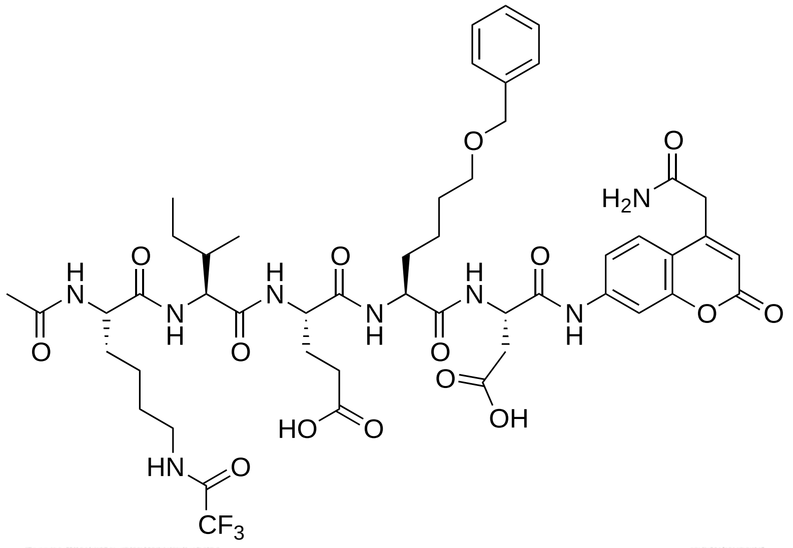


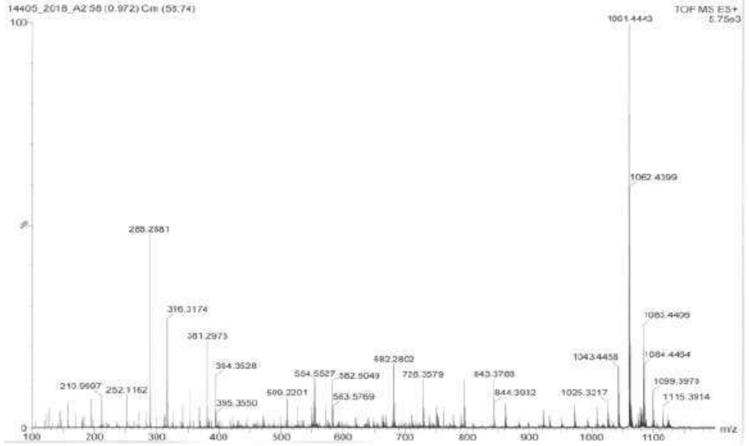


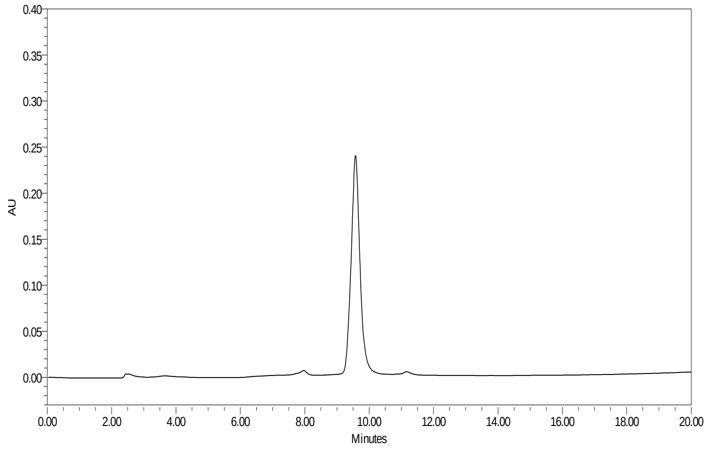

**TJ48 Ac-Phe(2-Cl)-Ile-Glu-Nle(O-Bzl)-Asp-ACC**

HRMS (m/z) [$MH^+$] calcd. for $C_{50}H_{60}ClN_7O_{14}$, 1017.39, found, 1018.55

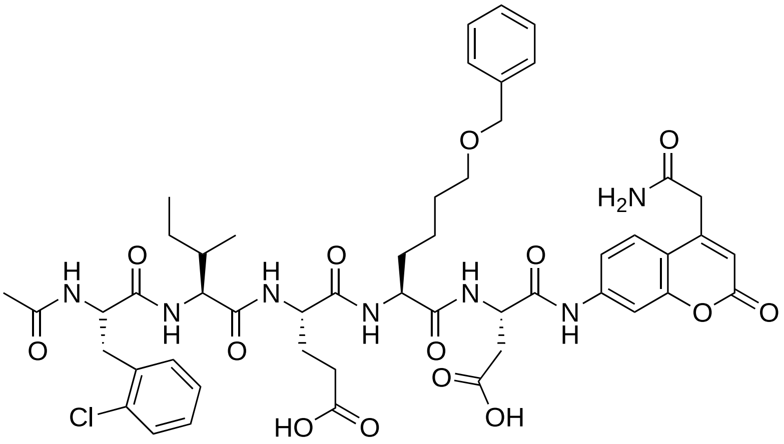


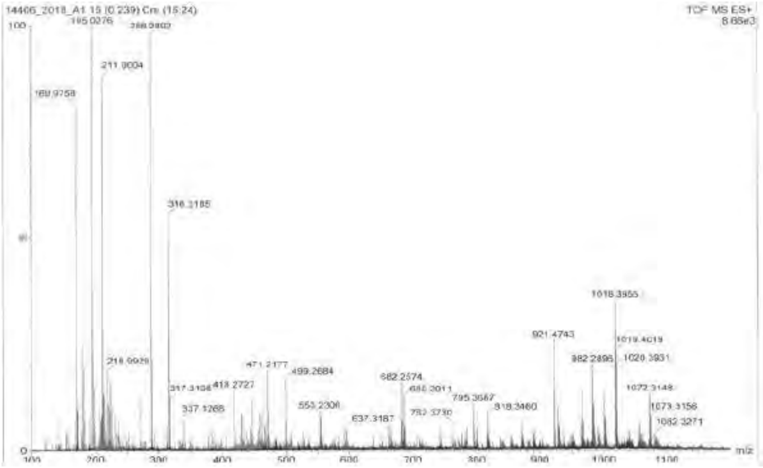


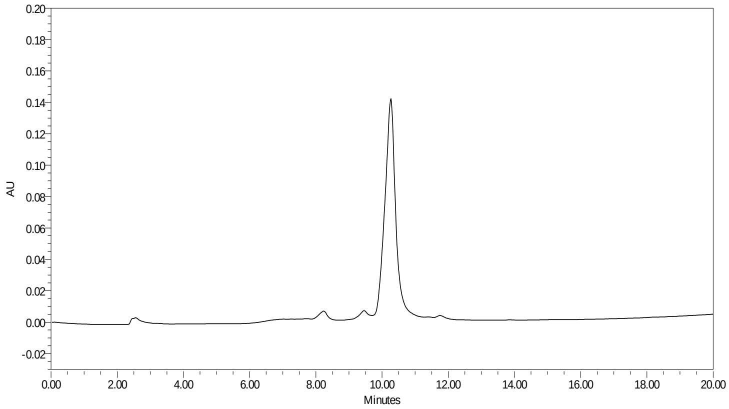

**TJ49 Ac-Nva-Ile-Glu-Nle(O-Bzl)-Asp-ACC**

HRMS (m/z) [$MH^+$] calcd. for $C_{46}H_{61}N_7O_{14}$, 935.43, found, 936.44

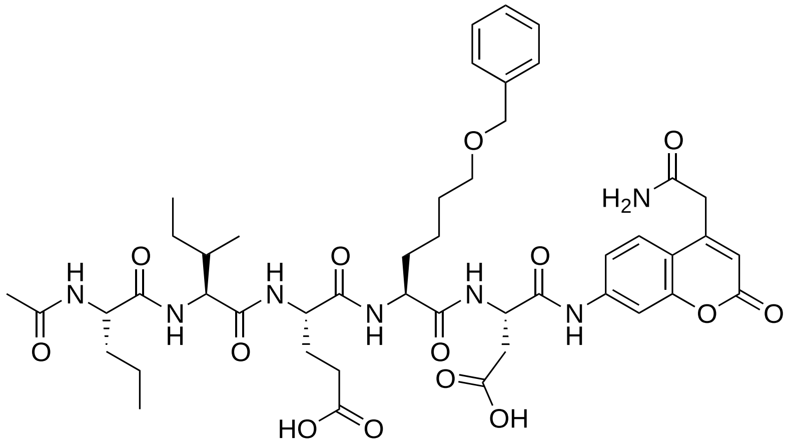

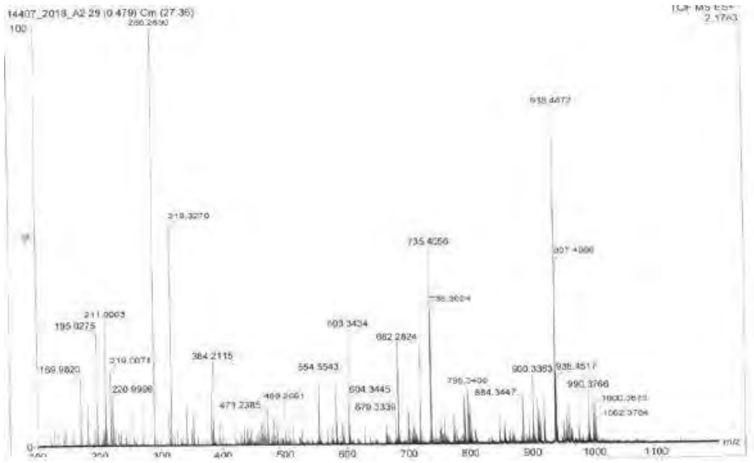

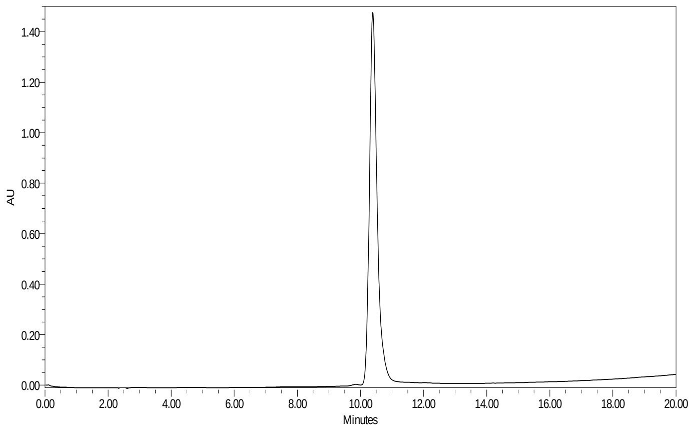

**TJ50 Ac-hPhe-Ile-Glu-Nle(O-Bzl)-Asp-ACC**

HRMS (m/z) [$MH^+$] calcd. for $C_{51}H_{63}N_7O_{14}$, 997.44, found, 998.45

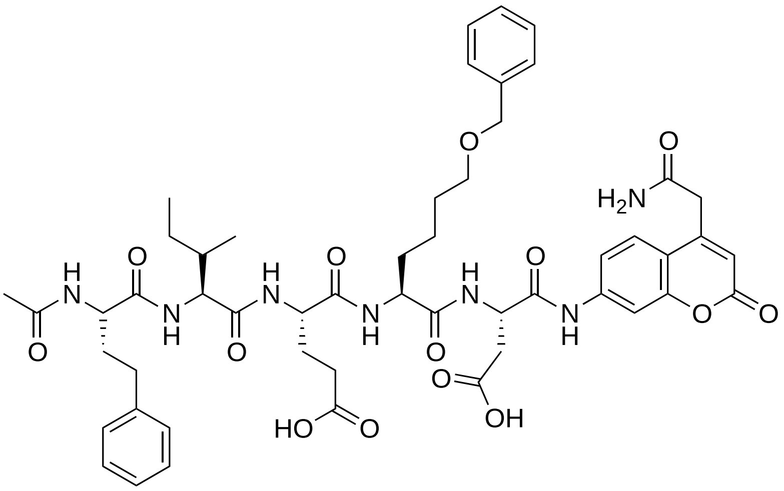

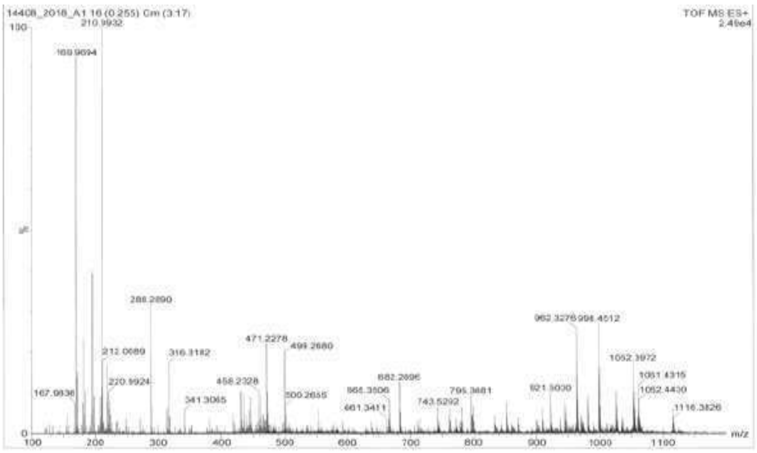


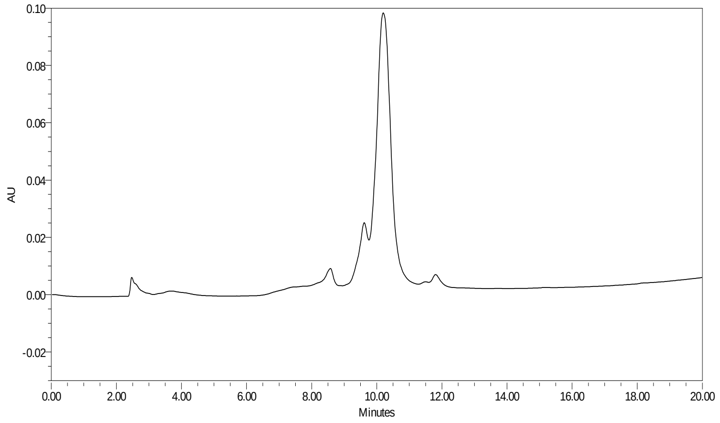

**TJ52 Ac-Ile-Ile-Glu-Oic-Asp-ACC**

HRMS (m/z) [MH$^+$] calcd. for $C_{43}H_{59}N_7O_{13}$, 881.42, found, 882.42

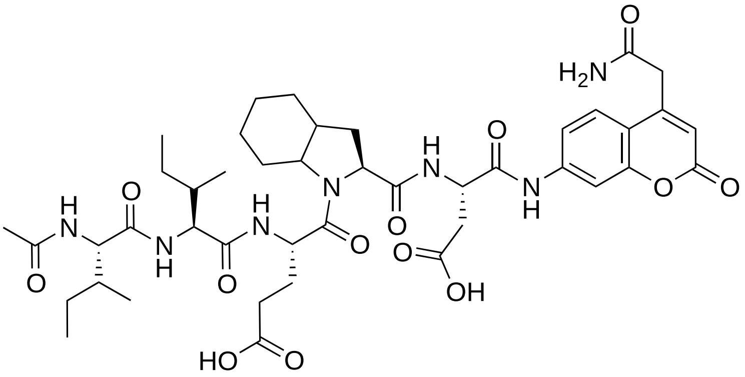


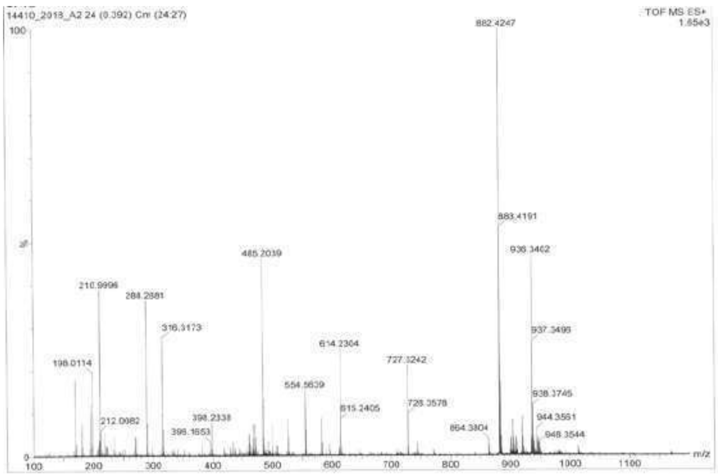


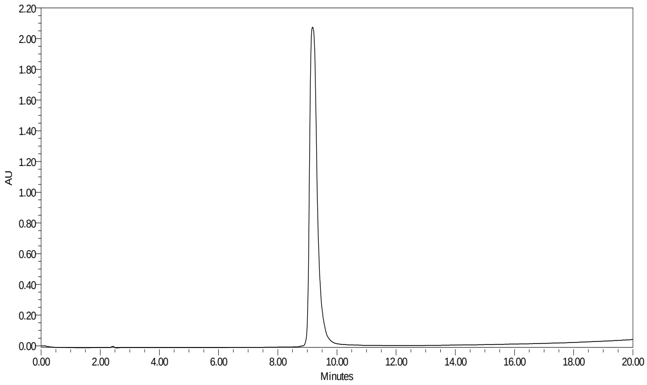

**TJ53 Ac-Lys(TFA)-Ile-Glu-Oic-Asp-ACC**

HRMS (m/z) [MH$^+$] calcd. for $C_{45}H_{59}F_3N_8O_{14}$, 992.41, found, 993.43

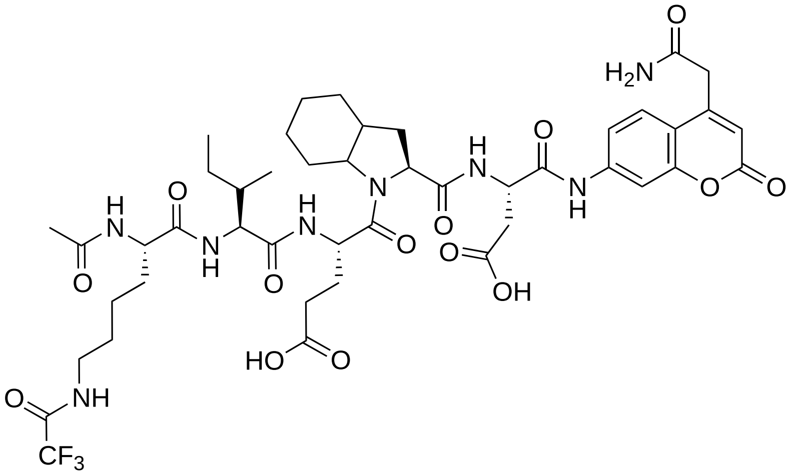

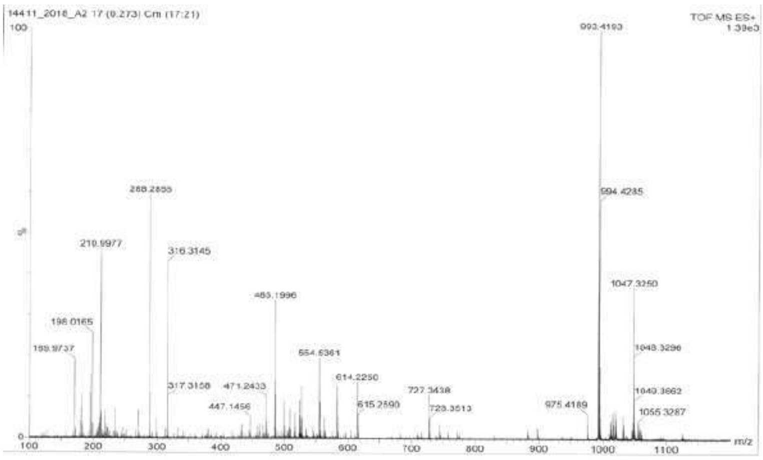


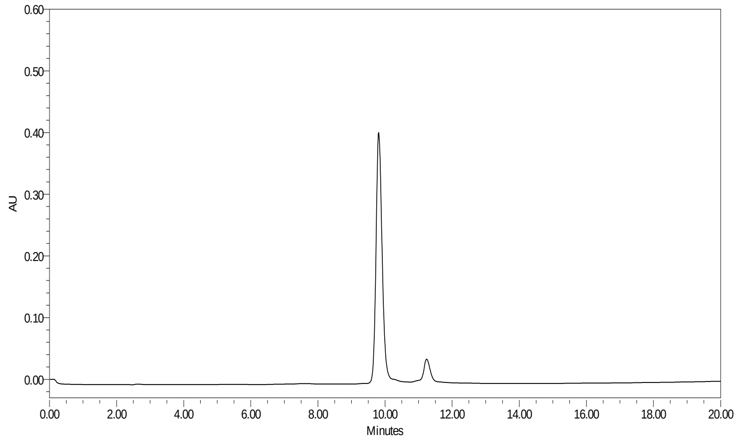

**TJ54 Ac-Phe(2-Cl)-Ile-Glu-Oic-Asp-ACC**

HRMS (m/z) [$MH^+$] calcd. for $C_{46}H_{56}ClN_7O_{13}$, 949.36, found, 950.37

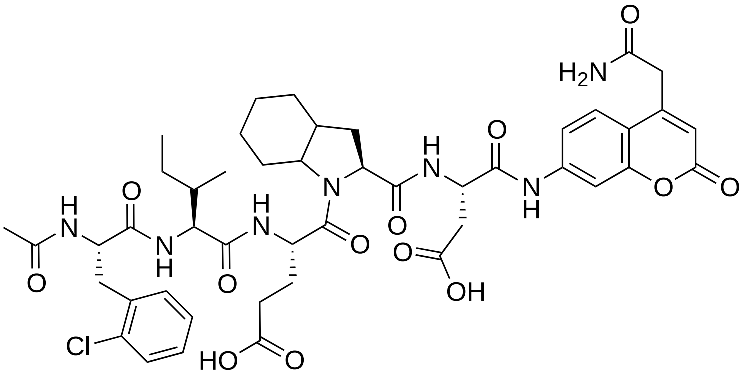


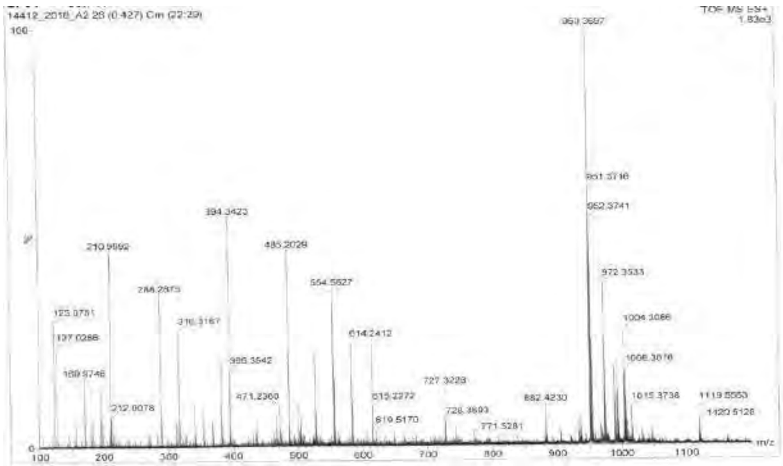


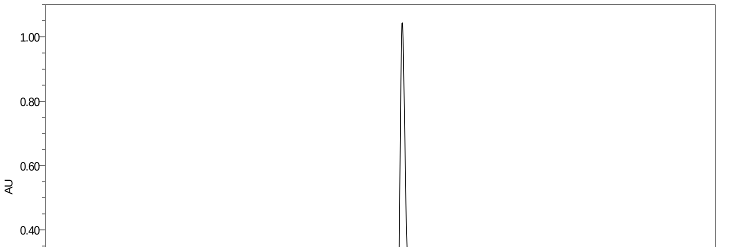

**TJ55 Ac-Nva-Ile-Glu-Oic-Asp-ACC**

HRMS (m/z) [MH$^+$] calcd. for $C_{42}H_{57}N_7O_{13}$, 867.40, found, 868.40

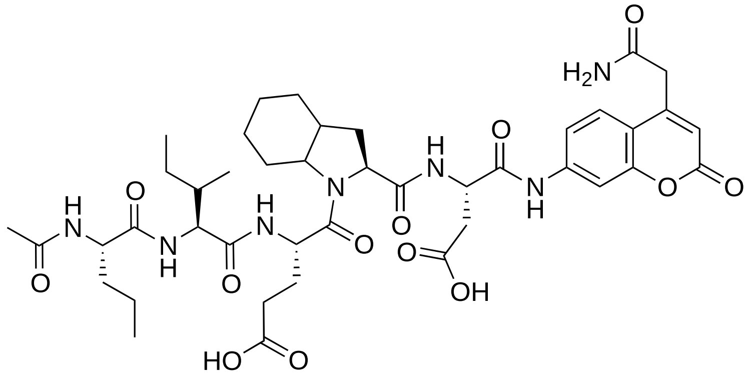

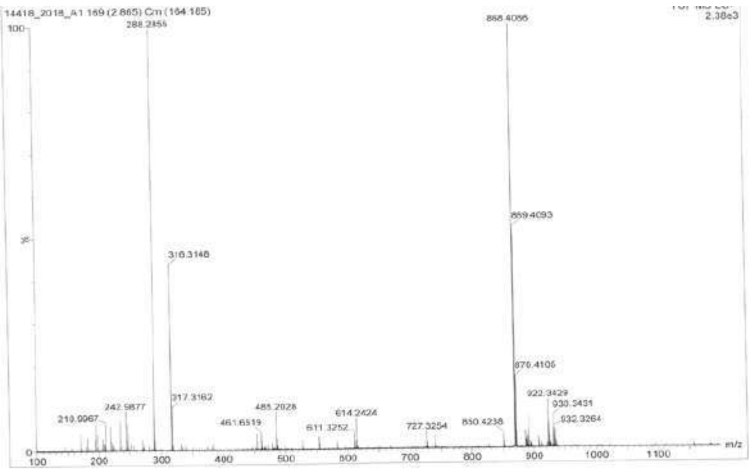


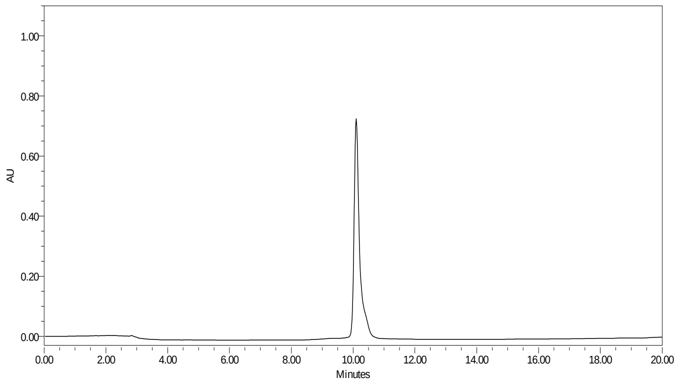

**TJ56 Ac-hPhe-Ile-Glu-Oic-Asp-ACC**

HRMS (m/z) [$MH^+$] calcd. for $C_{47}H_{59}N_7O_{13}$, 929.42, found, 930.42

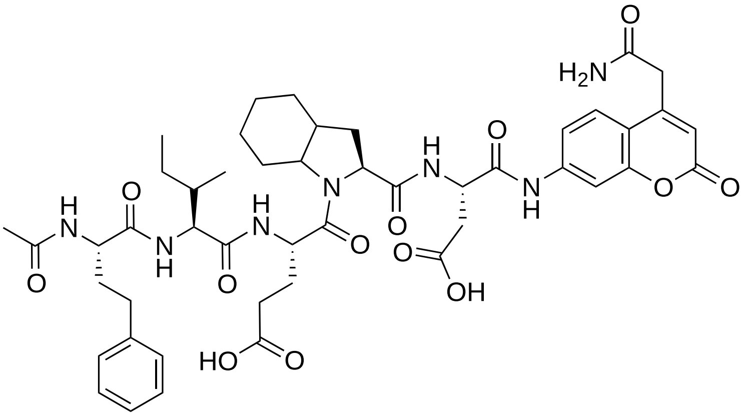

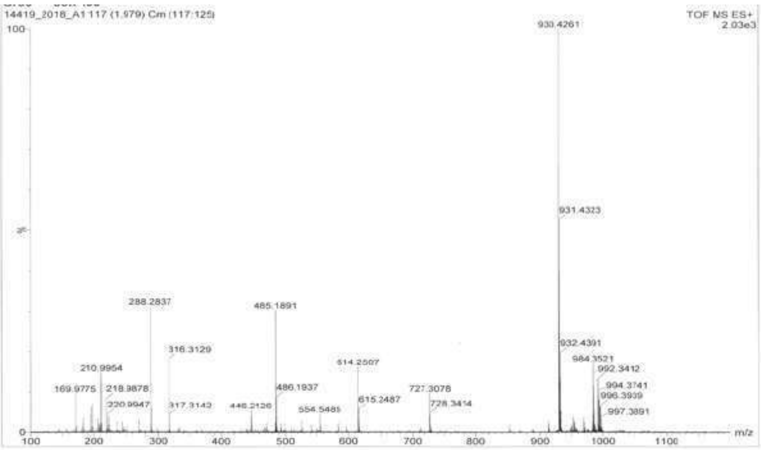


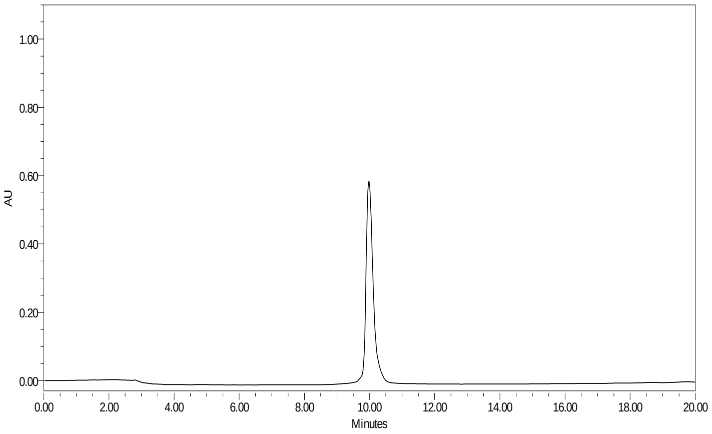

**TJ65 $H_2N$-ACC-Peg(4)-Nva-Ile-Glu-Oic-Asp-Phe-Gly-Arg-Lys(Dnp)**

HRMS (m/z) [$MH^+$] calcd. for $C_{80}H_{114}N_{18}O_{25}$, 1726.82, found, 864.88

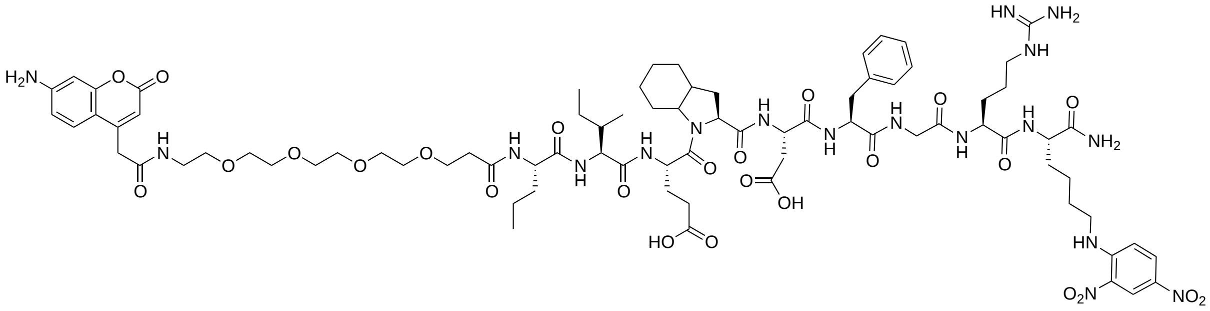

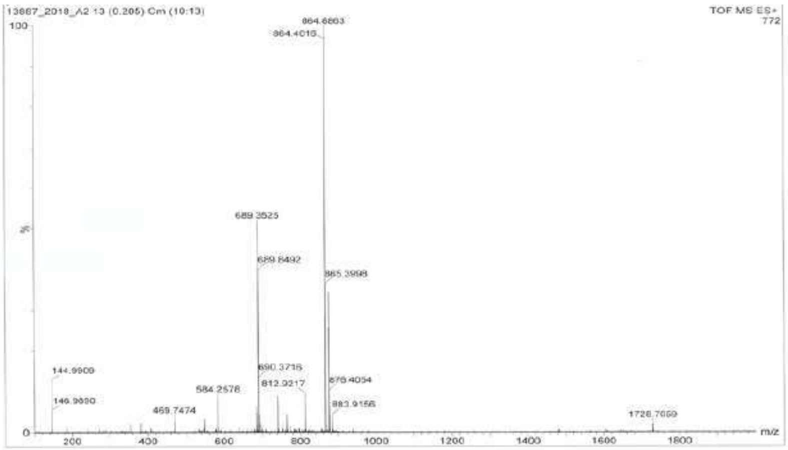


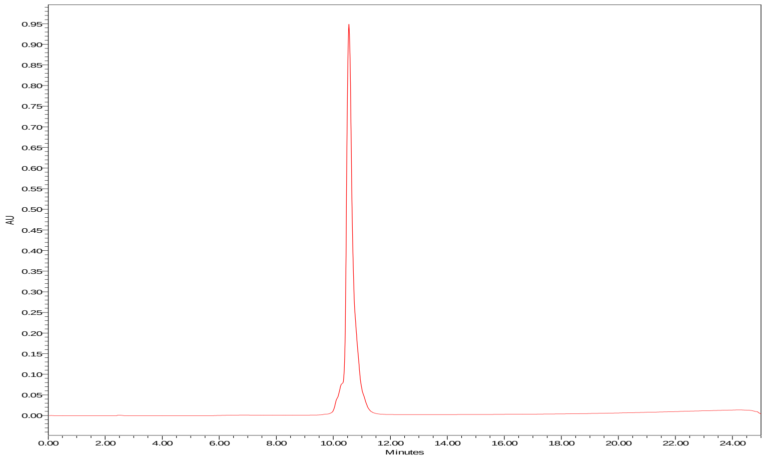

**TJ66 $H_2N$-ACC-Peg(4)-Nva-Ile-Glu-Oic-Asp-Gly-Gly-Gly-Lys(Dnp)**

HRMS (m/z) [$MH^+$] calcd. for $C_{69}H_{99}N_{15}O_{15}$, 1537.39, found, 1538.71

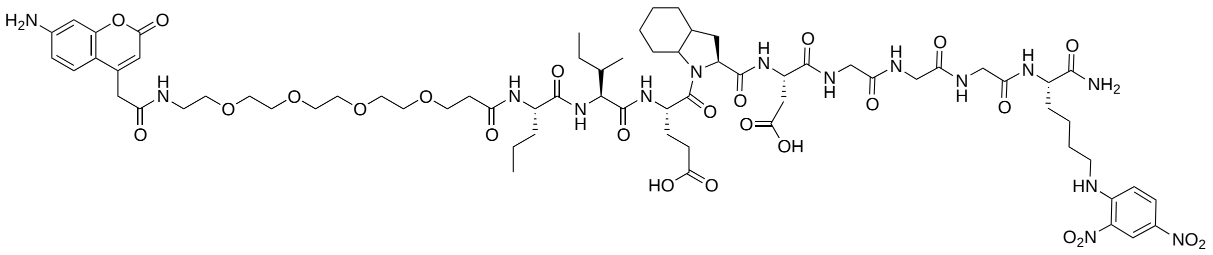

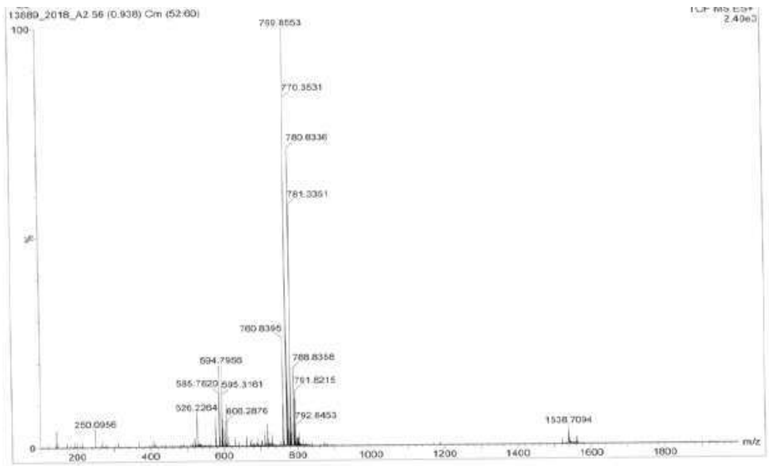


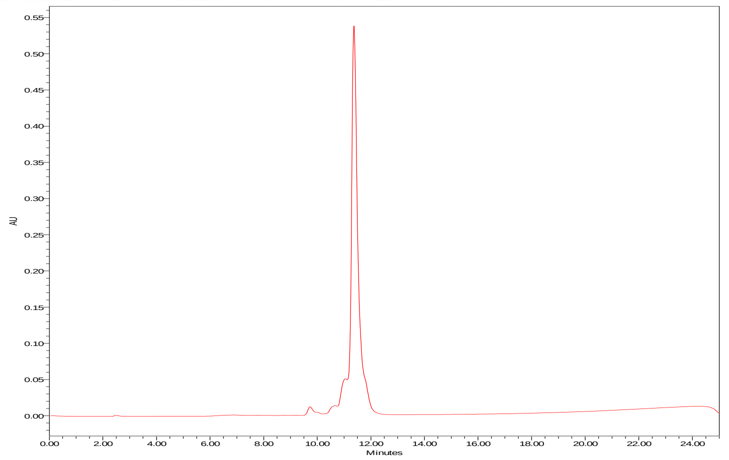

**TJ67 $H_2N$-ACC-Peg(4)-Nva-Ile-Glu-Oic-Asp-Peg(4)-Lys(Dnp)**

HRMS (m/z) [$MH^+$] calcd. for $C_{74}H_{111}N_{13}O_{27}$, 1613.77, found, 1614.76

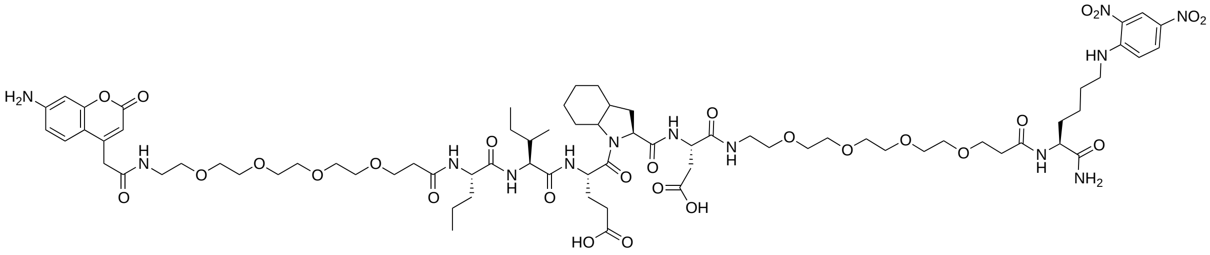


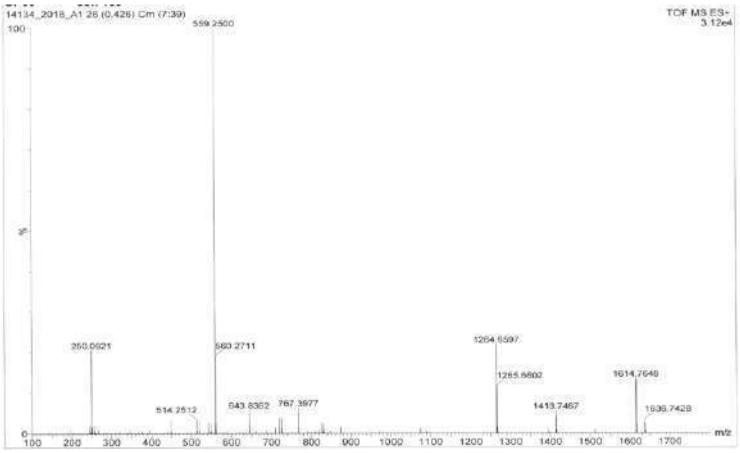


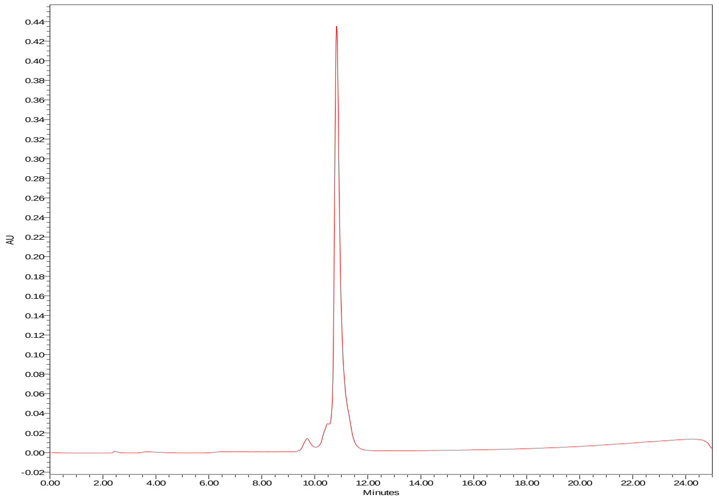

**TJ68 $H_2N$-ACC-Peg(4)-Nva-Ile-Glu-Nle(O-Bzl)-Asp-Phe-Gly-Arg-Lys(Dnp)**

HRMS (m/z) [MH$^+$] calcd. for $C_{84}H_{118}N_{18}O_{26}$, 1794.85, found, 909.93

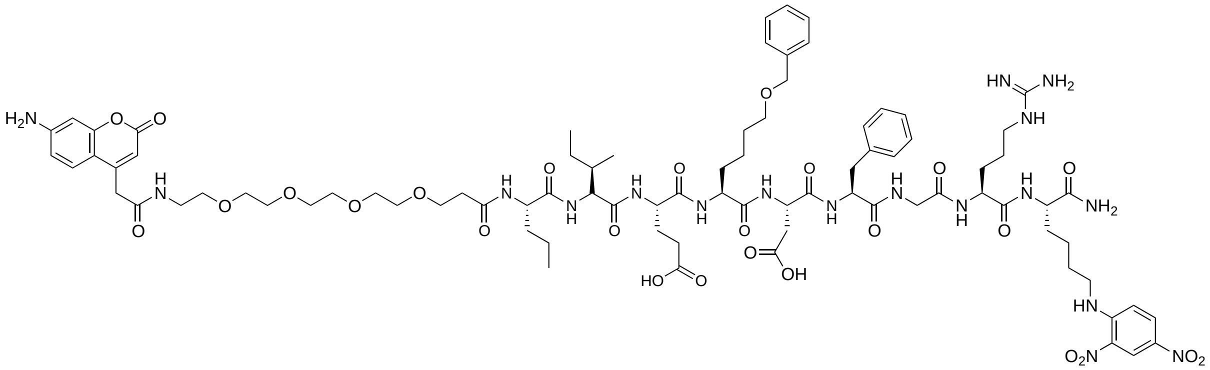


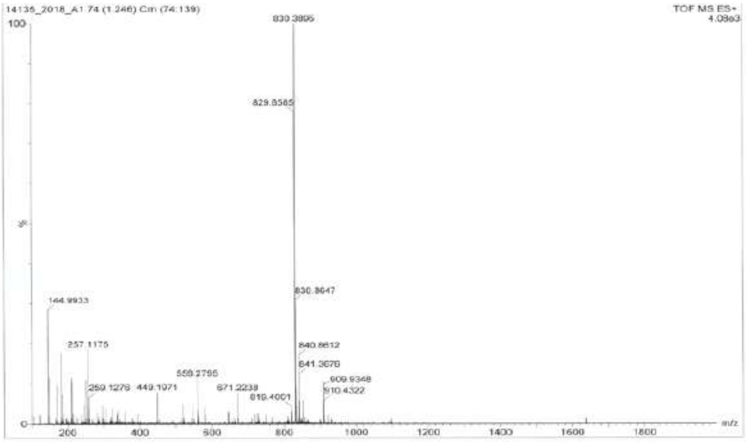


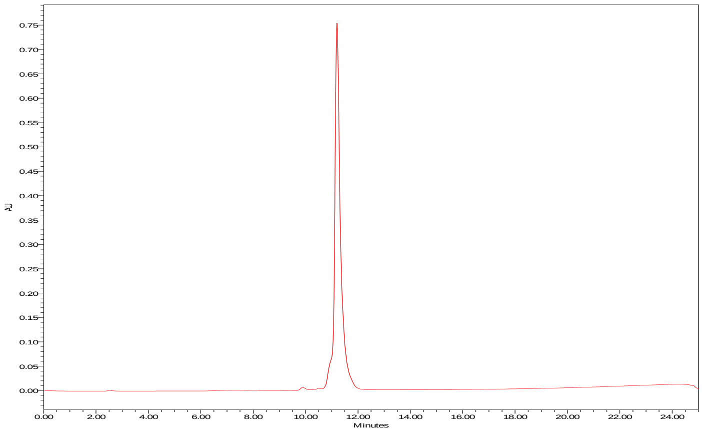

**TJ69 $H_2N$-ACC-Peg(4)-Nva-Ile-Glu-Nle(O-Bzl)-Asp-Gly-Gly-Gly-Lys(Dnp)**

HRMS (m/z) [$MH^+$] calcd. for $C_{73}H_{103}N_{15}O_{26}$, 1605.72, found, 1606.72

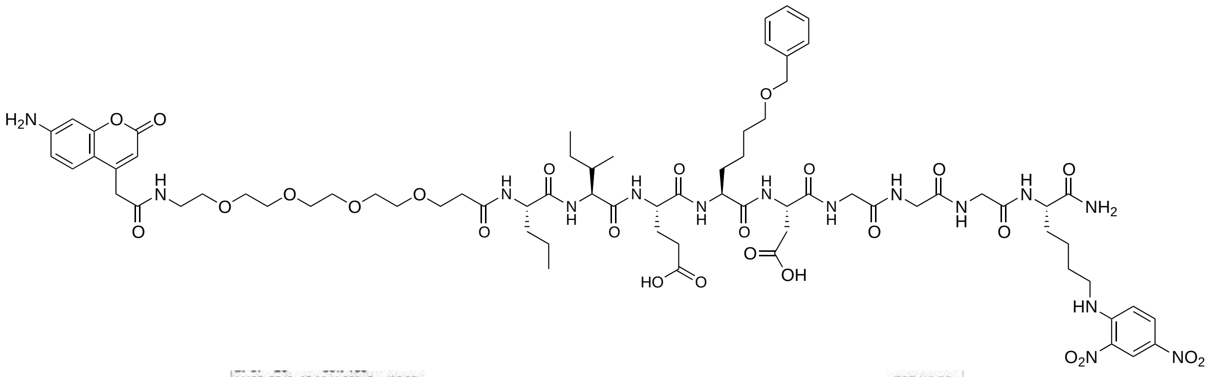

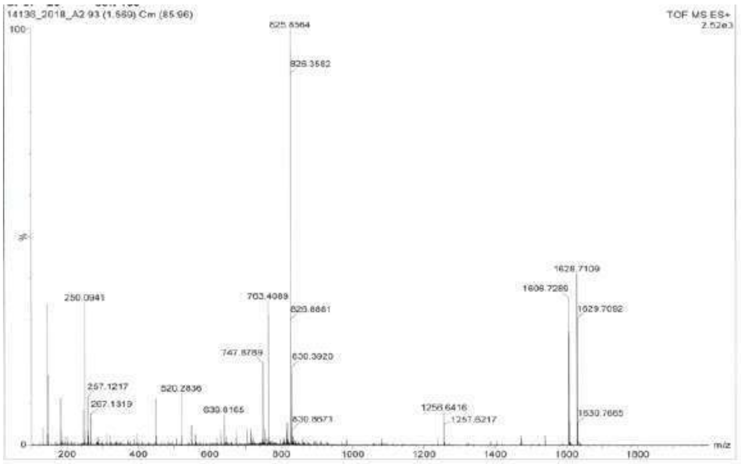


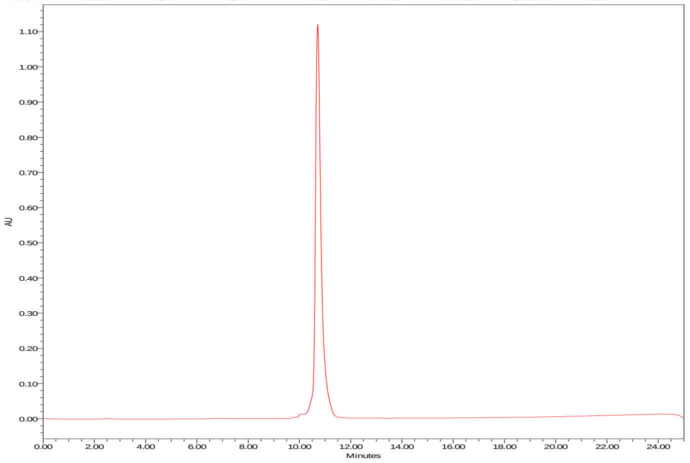

**TJ70 $H_2N$-ACC-Peg(4)-Nva-Ile-Glu-Nle(O-Bzl)-Asp-Peg(4)-Lys(Dnp)**

HRMS (m/z) [$MH^+$] calcd. for $C_{78}H_{115}N_{13}O_{28}$, 1681.91, found, 842.13

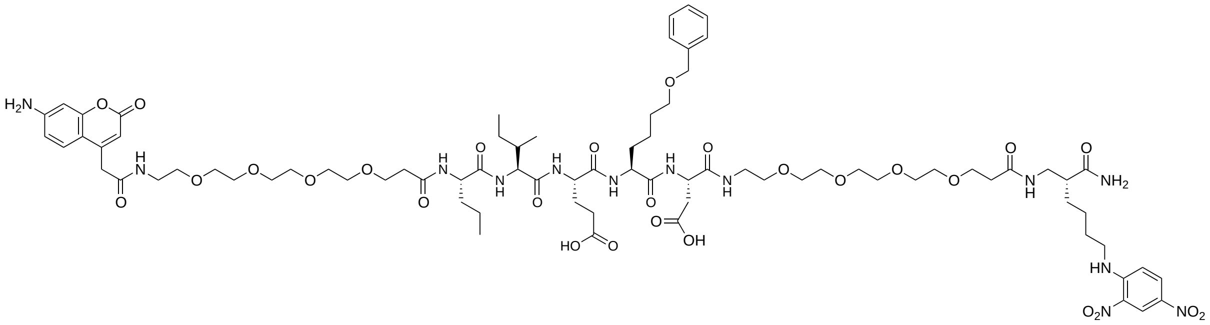

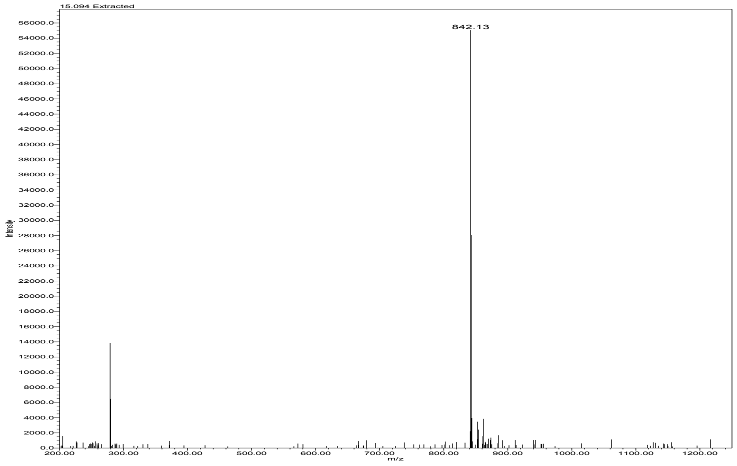


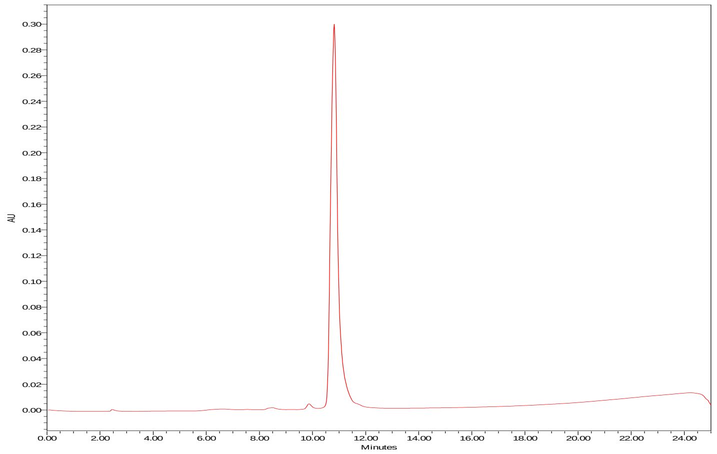

**TJ71 $H_2N$-ACC-Nva-Ile-Glu-Oic-Asp-Phe-Gly-Arg-Lys(Dnp)**

HRMS (m/z) [$MH^+$] calcd. for $C_{69}H_{93}N_{17}O_{20}$, 1479.68, found, 1480.68

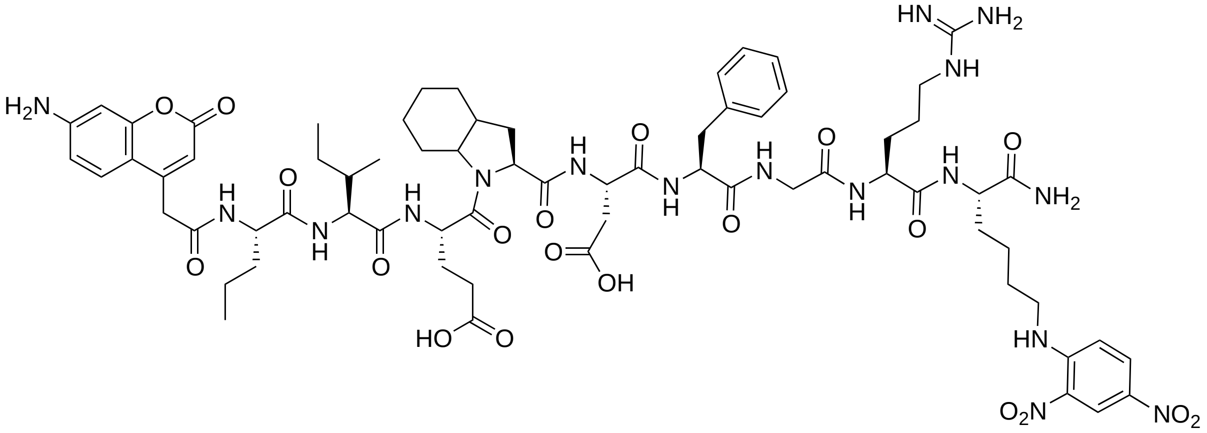

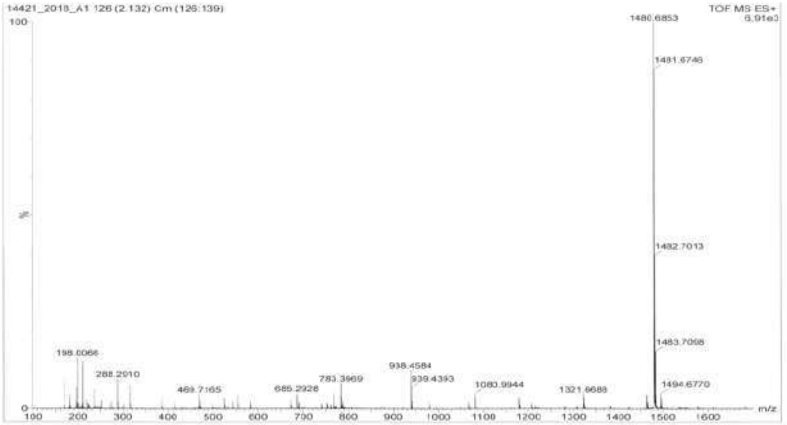


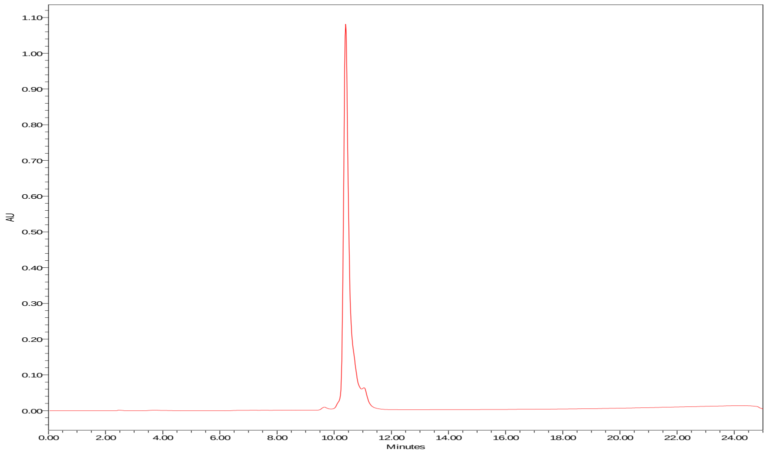

**TJ72 $H_2N$-ACC-Nva-Ile-Glu-Oic-Asp-Gly-Gly-Gly-Lys(Dnp)**

HRMS (m/z) [$MH^+$] calcd. for $C_{58}H_{78}N_{14}O_{20}$, 1290.55, found, 1291.56

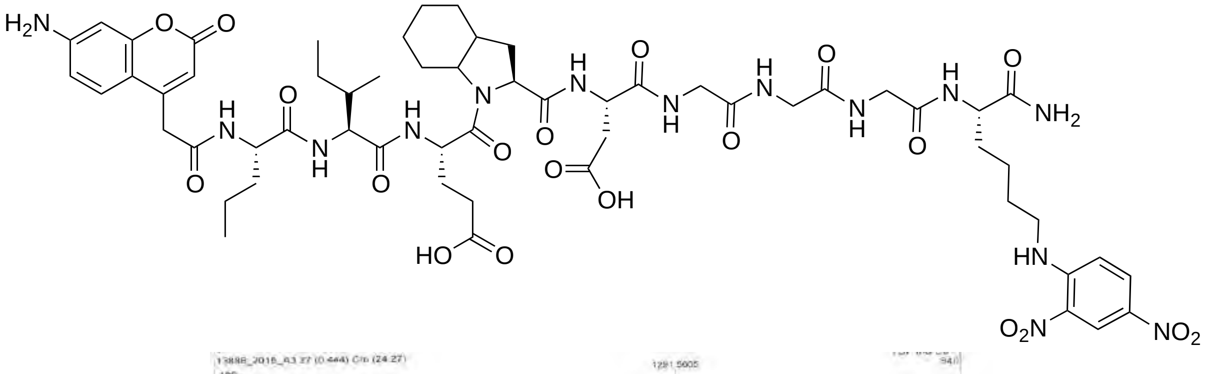

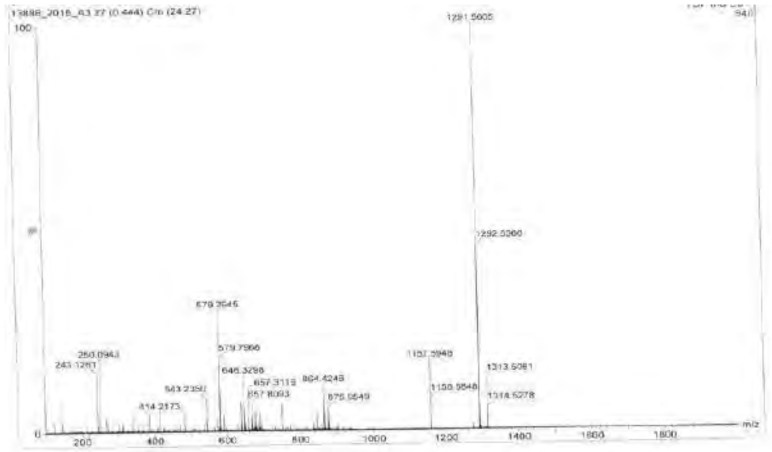

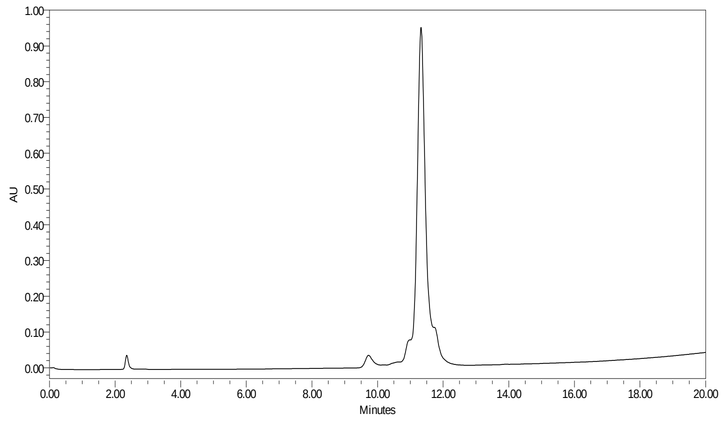

**TJ73 $H_2N$-ACC-Nva-Ile-Glu-Oic-Asp-Peg(4)-Lys(Dnp)**

HRMS (m/z) [$MH^+$] calcd. for $C_{63}H_{90}N_{12}O_{22}$, 1366.63, found, 1367.33

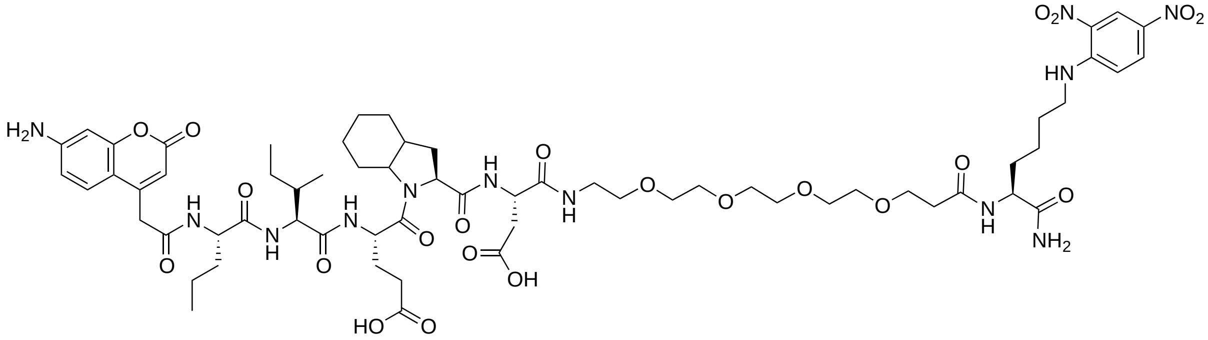

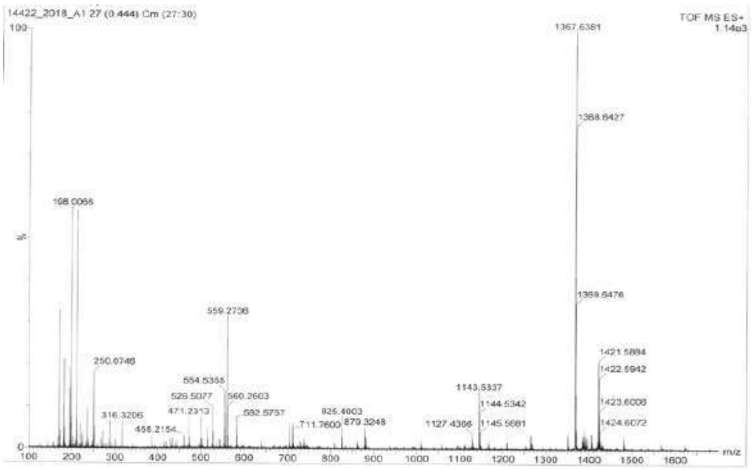


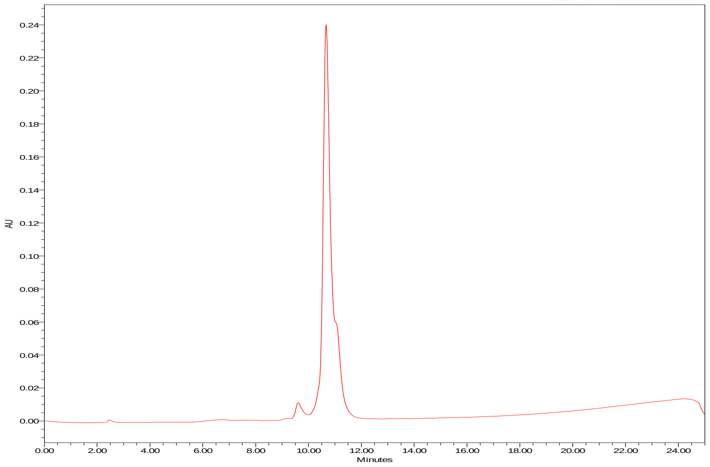

**TJ74 $H_2N$-ACC-Nva-Ile-Glu-Nle(O-Bzl)-Asp-Phe-Gly-Arg-Lys(Dnp)**

HRMS (m/z) [$MH^+$] calcd. for $C_{73}H_{97}N_{17}O_{21}$, 1547.70, found, 1548.71

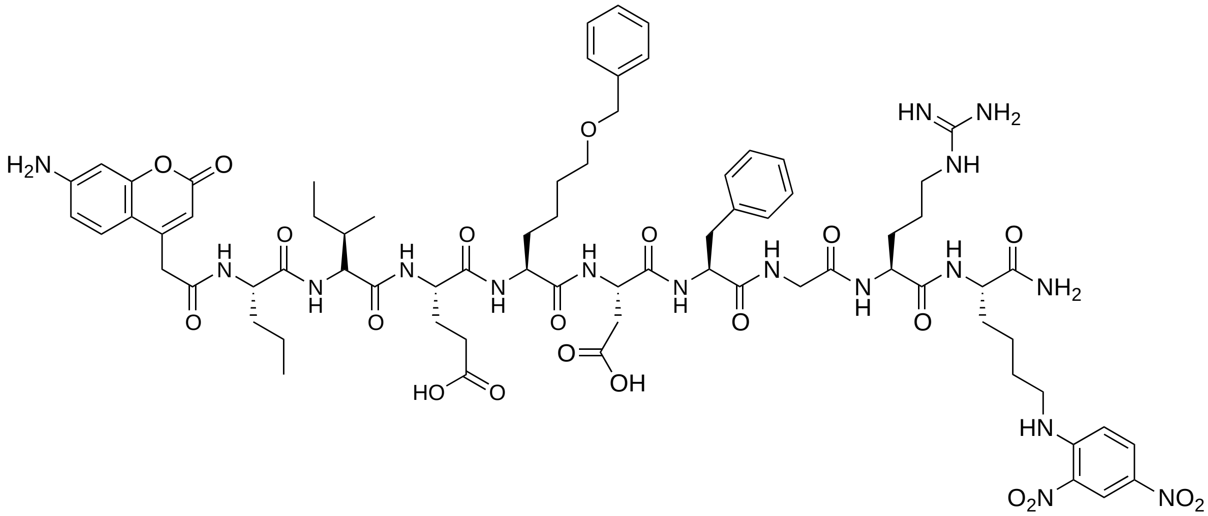

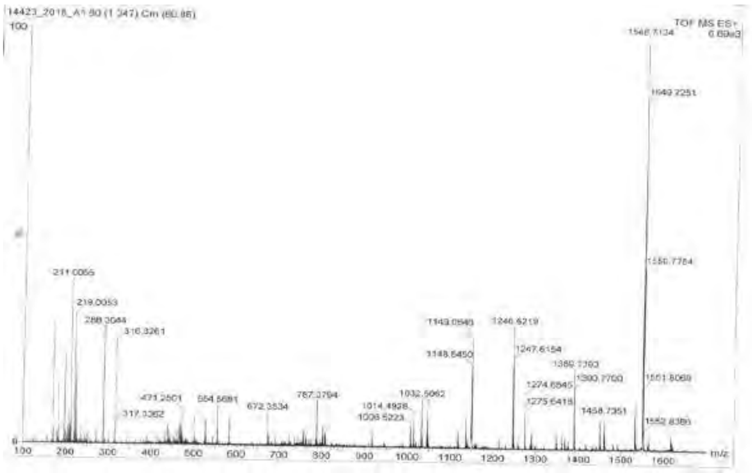


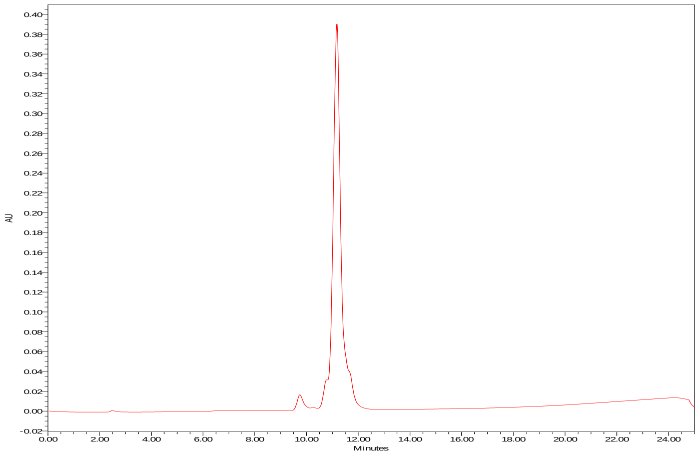

**TJ75 $H_2N$-ACC-Nva-Ile-Glu-Nle(O-Bzl)-Asp-Gly-Gly-Gly-Lys(Dnp)**

HRMS (m/z) [$MH^+$] calcd. for $C_{62}H_{82}N_{14}O_{21}$, 1358.58, found, 1359.58

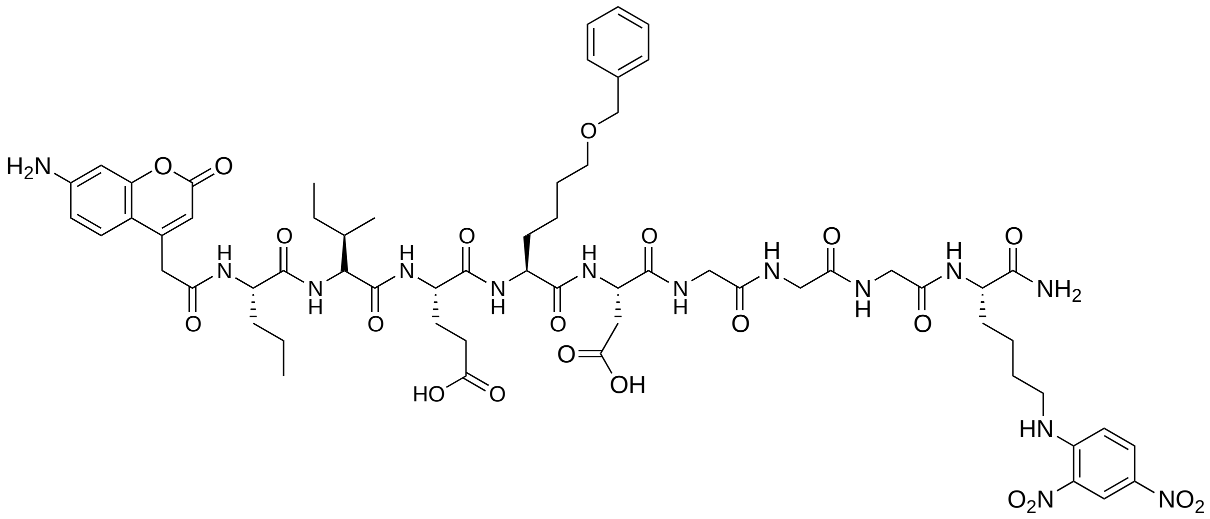

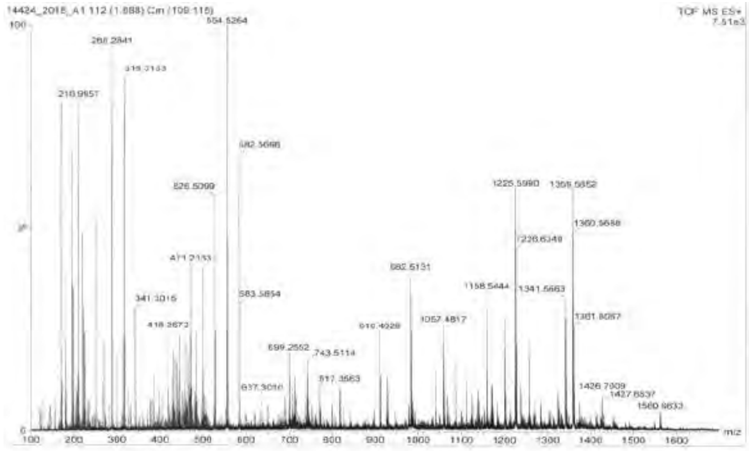

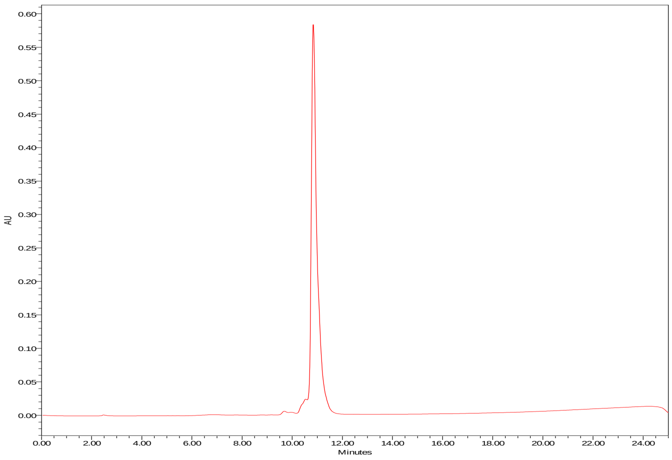

**TJ76 $H_2N$-ACC-Nva-Ile-Glu-Nle(O-Bzl)-Asp-Peg(4)-Lys(Dnp)**

HRMS (m/z) [$MH^+$] calcd. for $C_{69}H_{94}N_{12}O_{23}$, 1434.66, found, 1435.66

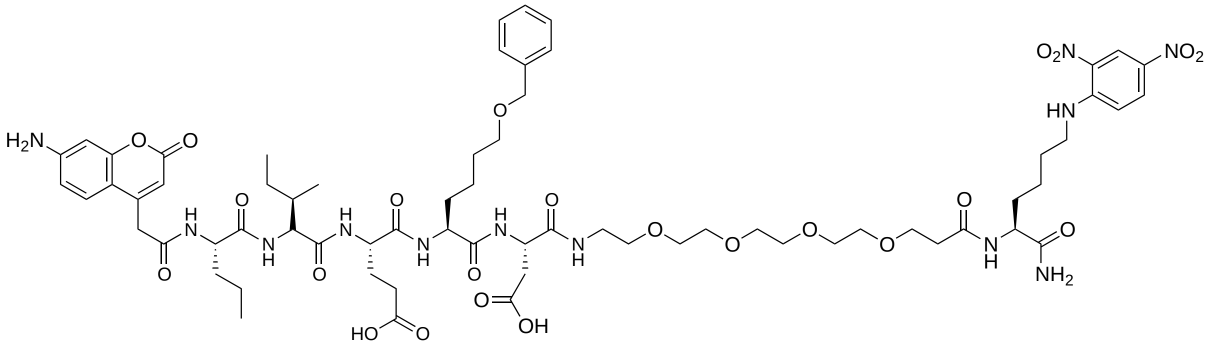

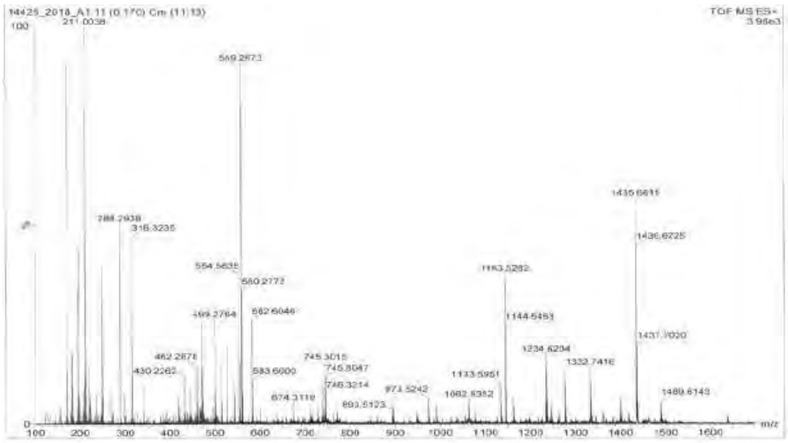


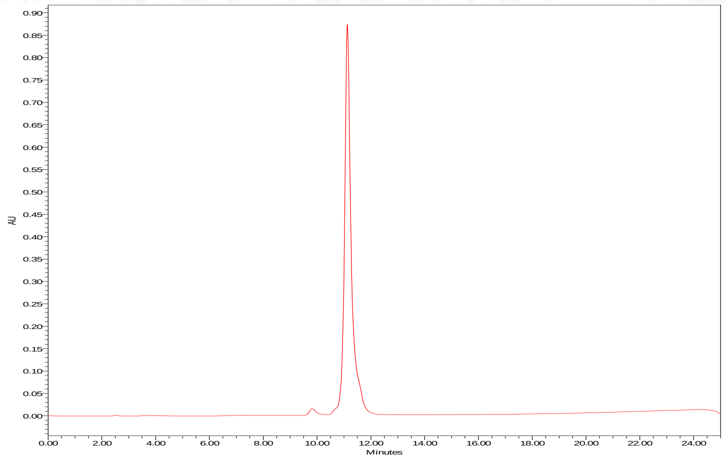

**qTJ71 BHQ2-Nva-Ile-Glu-Oic-Asp-Phe-Gly-Arg-Cy3**

HRMS (m/z) [$MH^+$] calcd. for $C_{107}H_{143}N_{22}O_{19}^+$, 2041.09, found,1021.07

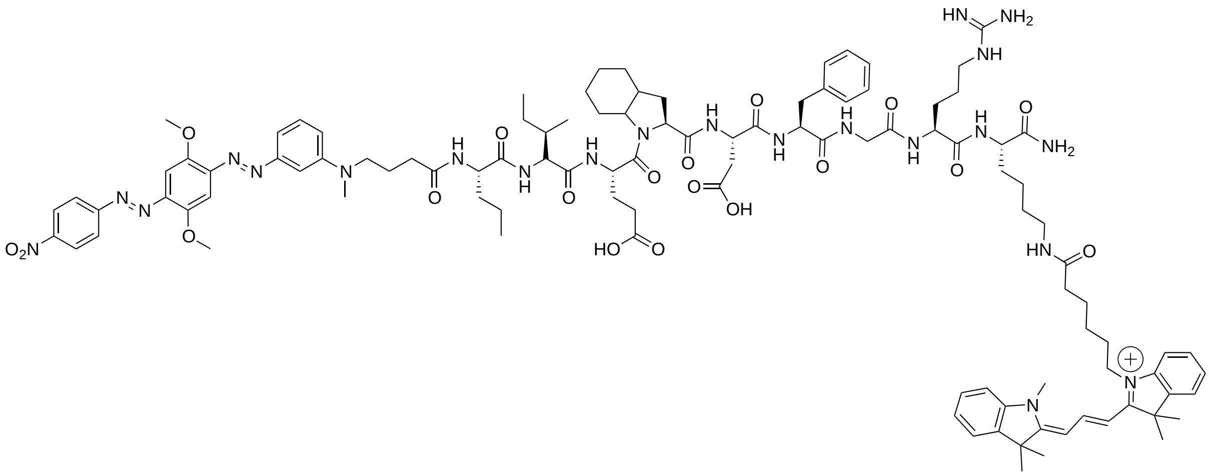


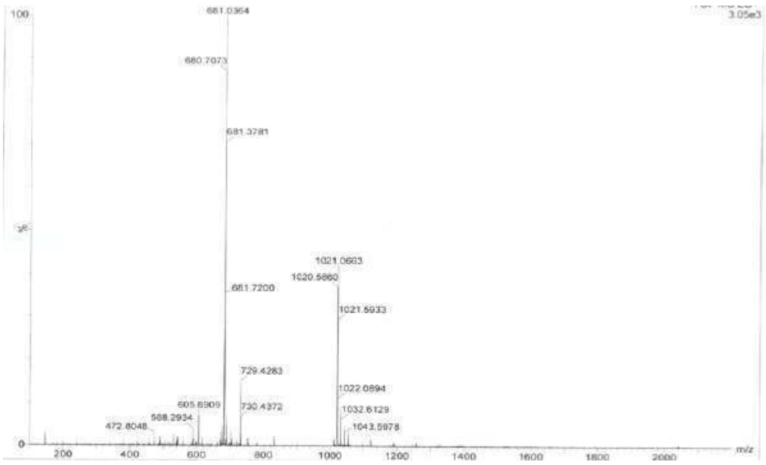


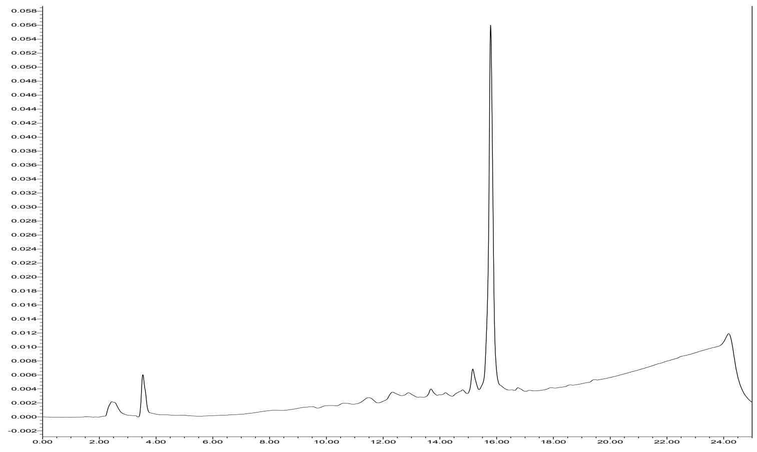

**TJ55.Bt Biot-Nva-Ile-Glu-Oic-Asp-PO$_3$Ph$_2$**

HRMS (m/z) [MH$^+$] calcd. for $C_{50}H_{70}N_7O_{13}PS$, 1039.45, found, 1040.47

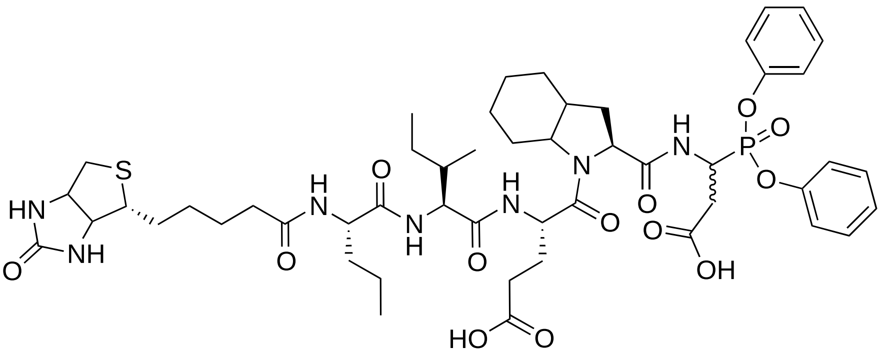

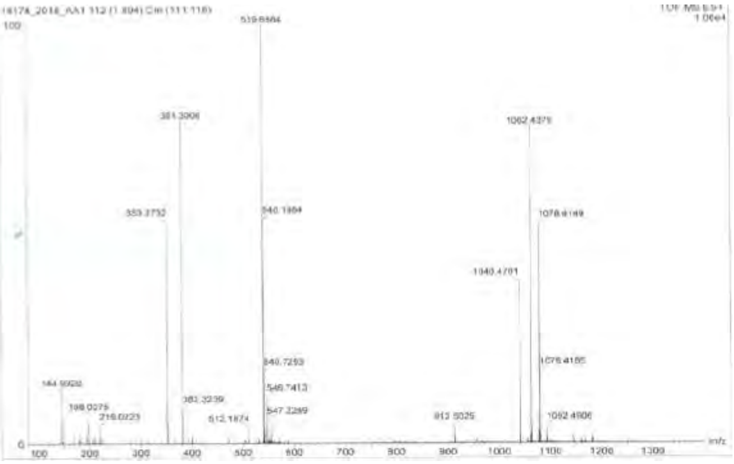

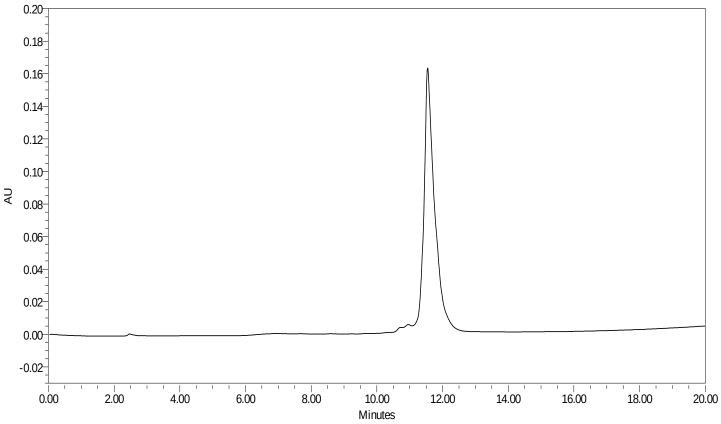

**TJ71i Ac-Nva-Ile-Glu-Oic-Asp-$PO_3Ph_2$**

HRMS (m/z) [$MH^+$] calcd. for $C_{42}H_{58}N_5O_{12}P$, 855.38, found, 856.32

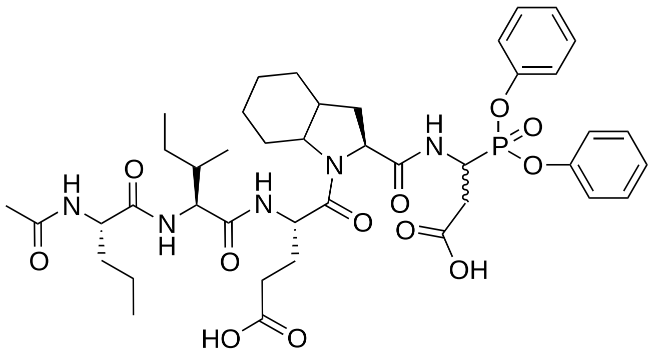

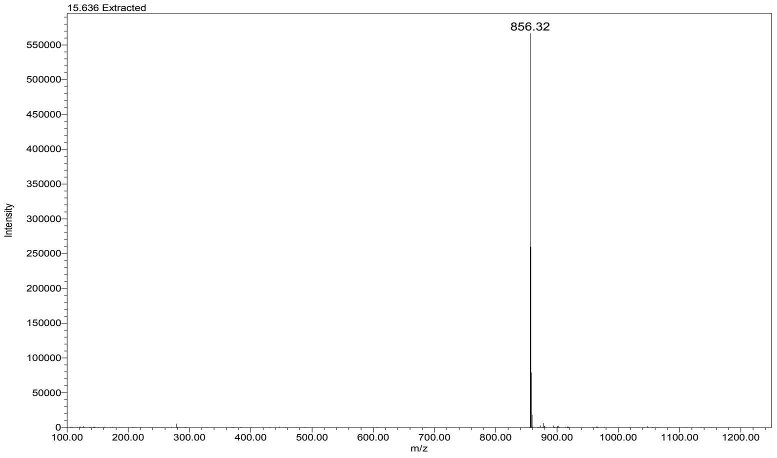


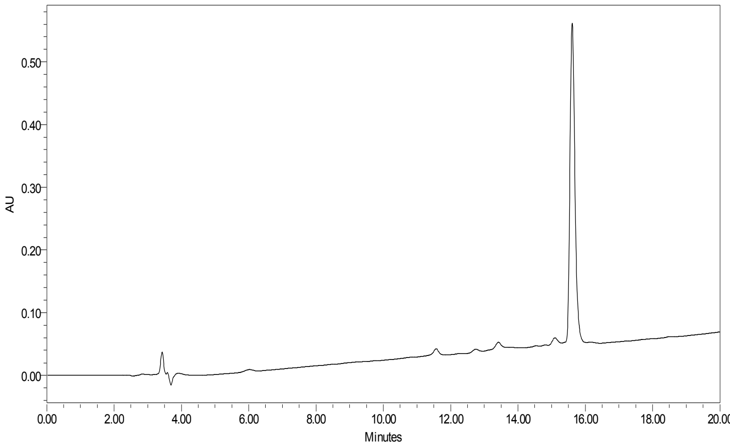

**TJ55.5 Cy5-Gly-Nva-Ile-Glu-Oic-Asp-$PO_3Ph_2$**

HRMS (m/z) [$MH^+$] calcd. for $C_{74}H_{96}N_8O_{13}P^+$, 1335.68, found, 668.28

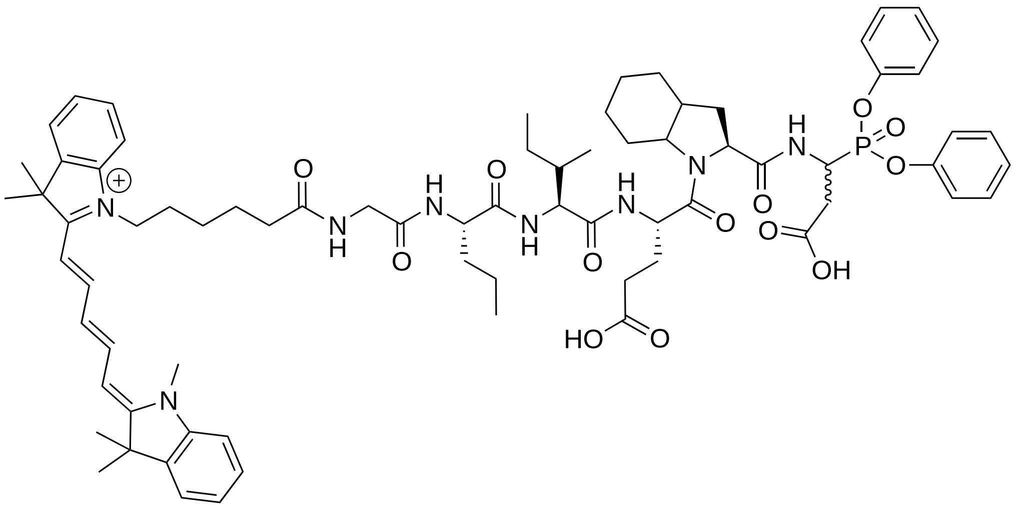

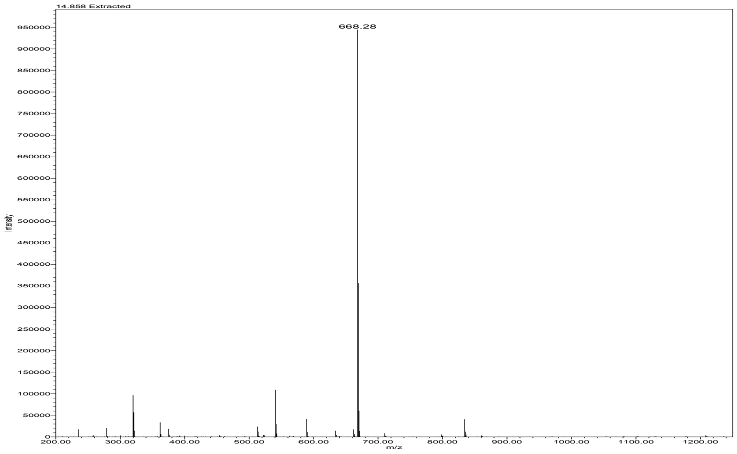


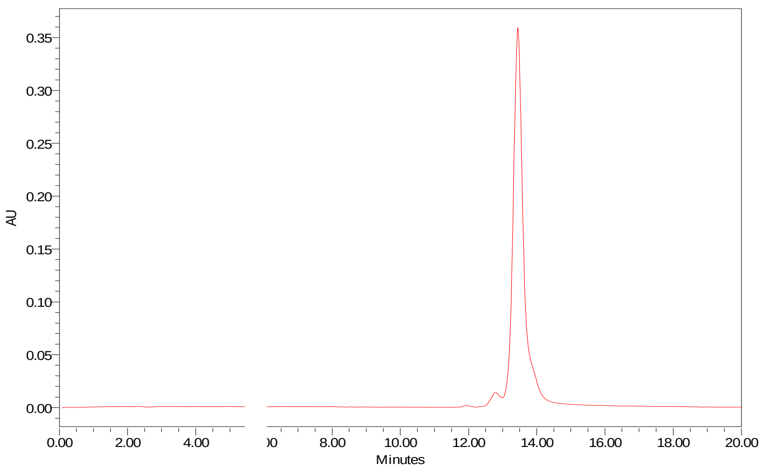

**TJ77 Ac-Asp-ACC**

HRMS (m/z) [$MH^+$] calcd. for $C_{17}H_{17}N_3O_7$, 375.11, found, 375.95

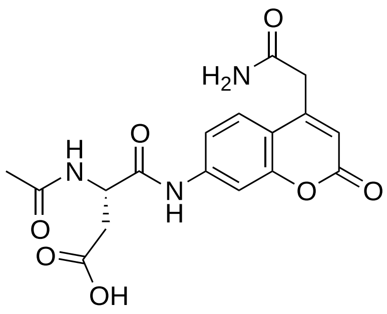


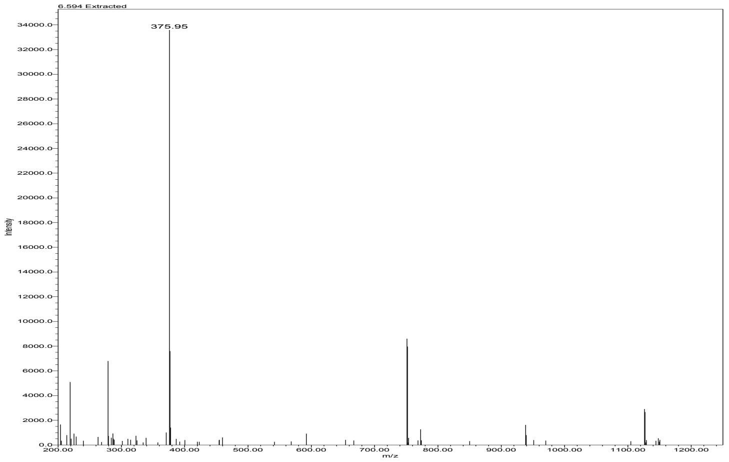


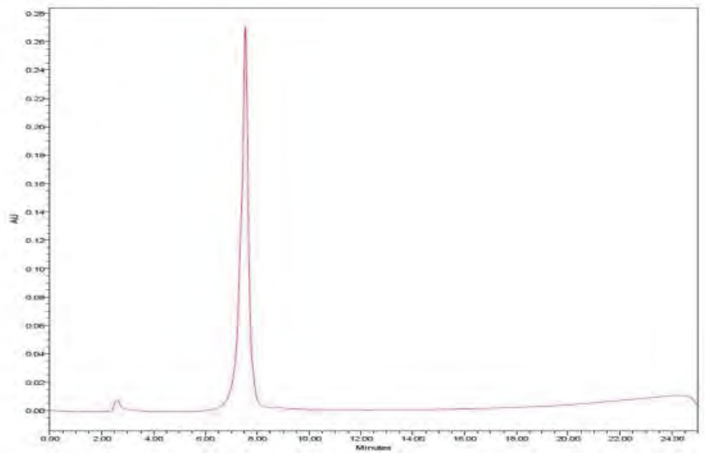

**TJ78 Ac-Pro-Asp-ACC**

HRMS (m/z) [MH$^+$] calcd. for $C_{22}H_{24}N_4O_8$, 472.16, found, 473.08

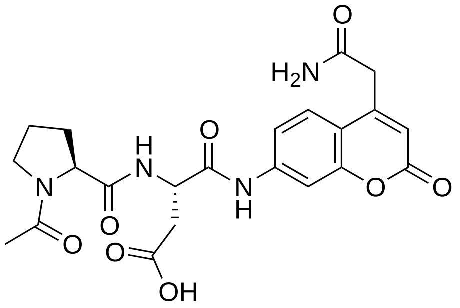


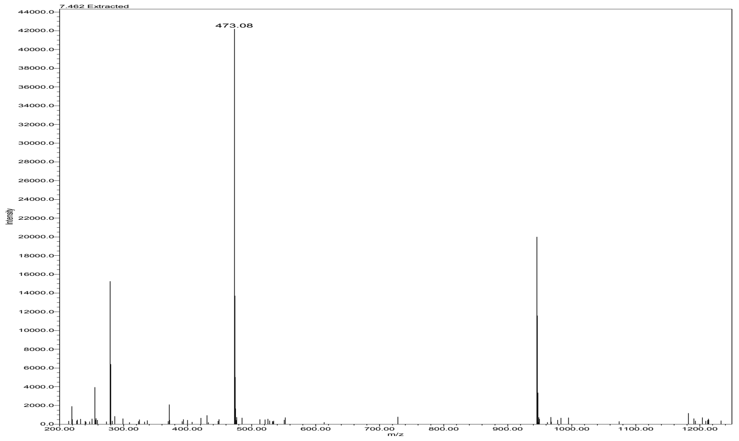


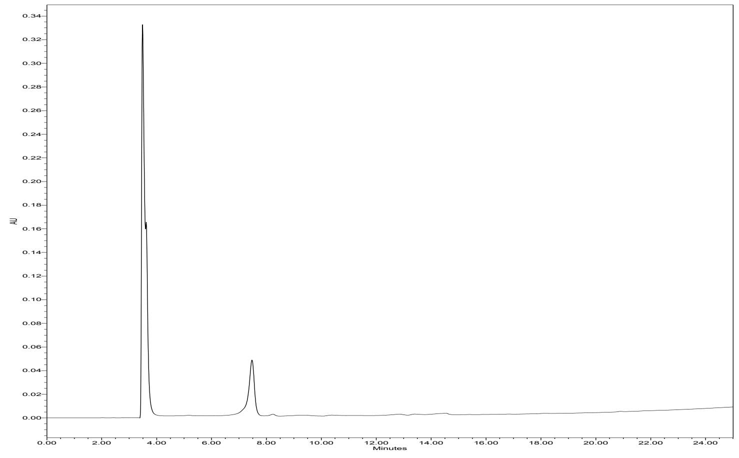

**TJ79 Ac-Glu-Pro-Asp-ACC**

HRMS (m/z) [MH+] calcd. for $C_{27}H_{31}N_5O_{11}$, 601.20, found, 602.24

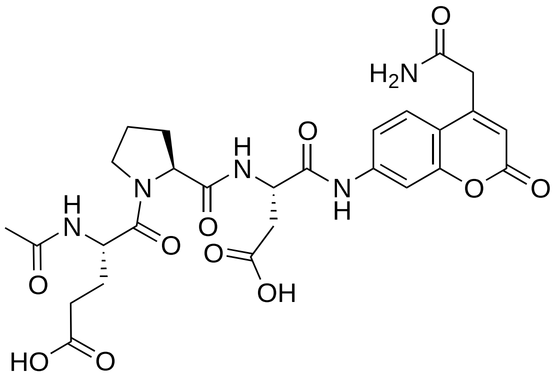

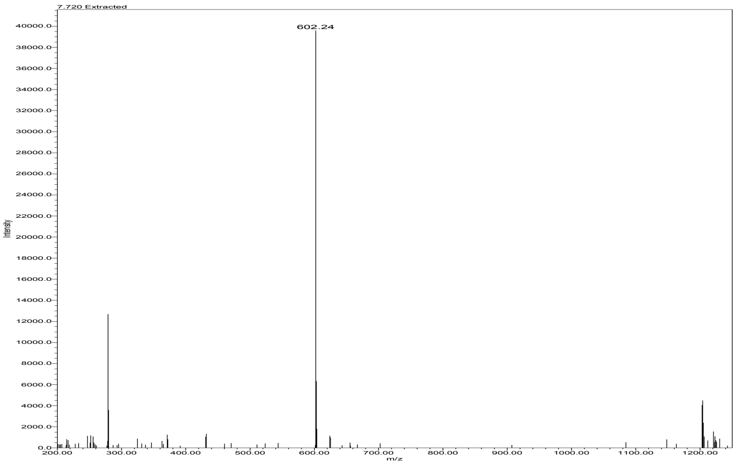


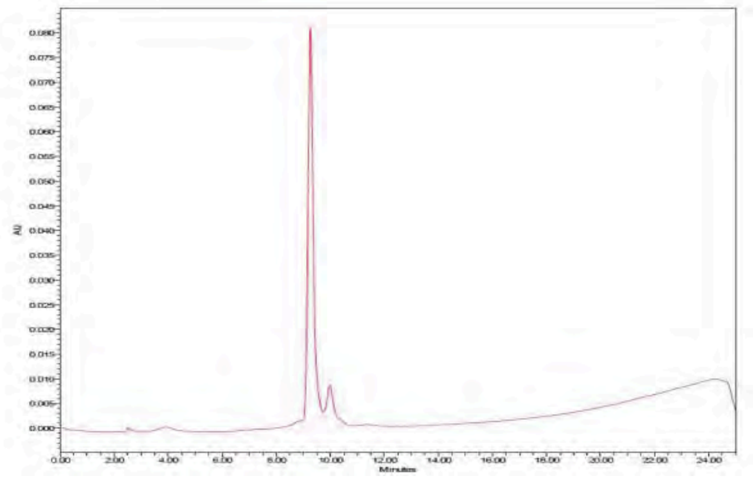

**TJ80 Ac-Ile-Glu-Pro-Asp-ACC**

HRMS (m/z) [MH$^+$] calcd. for $C_{33}H_{42}N_6O_{12}$, 714.29, found, 715.32

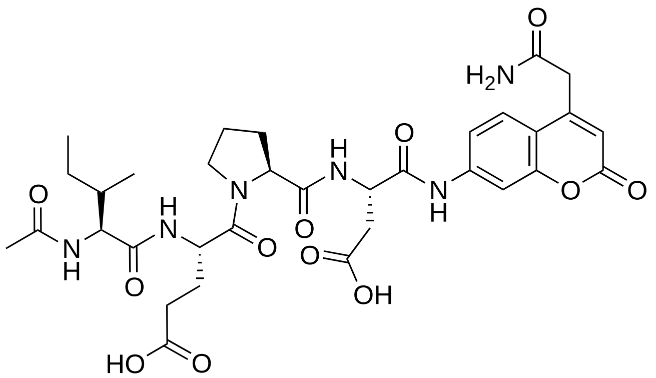

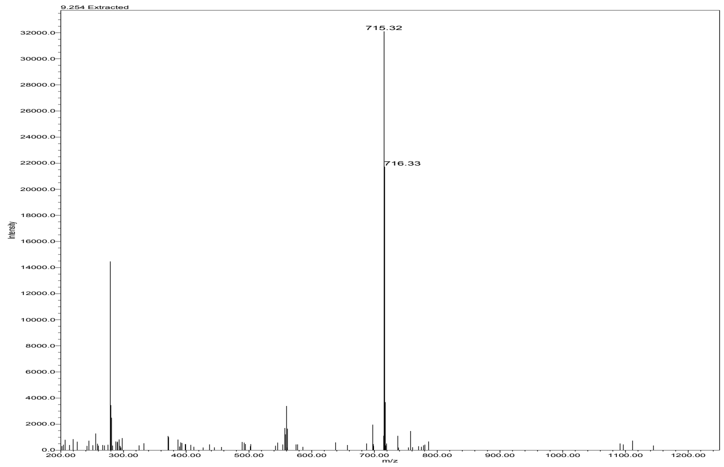


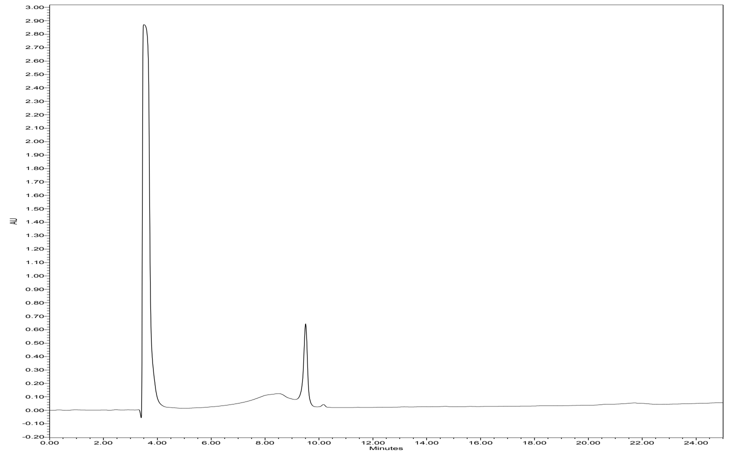

**TJ81 Ac-Ala-Ile-Glu-Pro-Asp-ACC**

HRMS (m/z) [$MH^+$] calcd. for $C_{36}H_{47}N_7O_{13}$, 785.32, found, 786.31

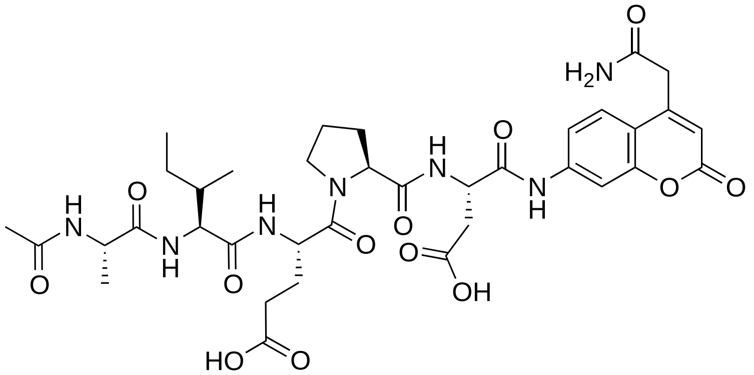

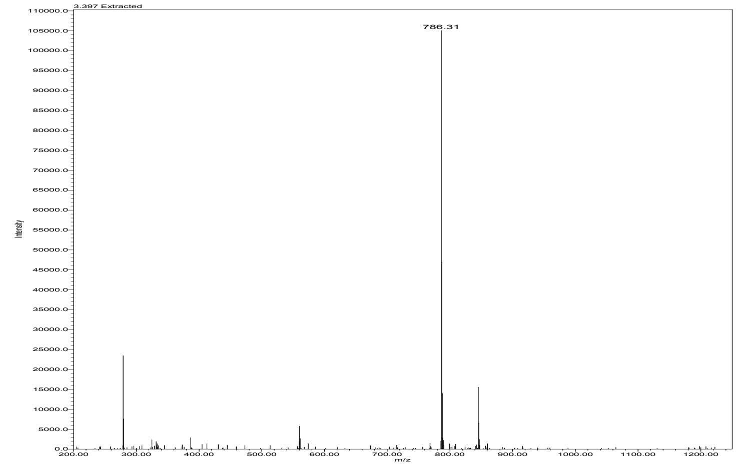


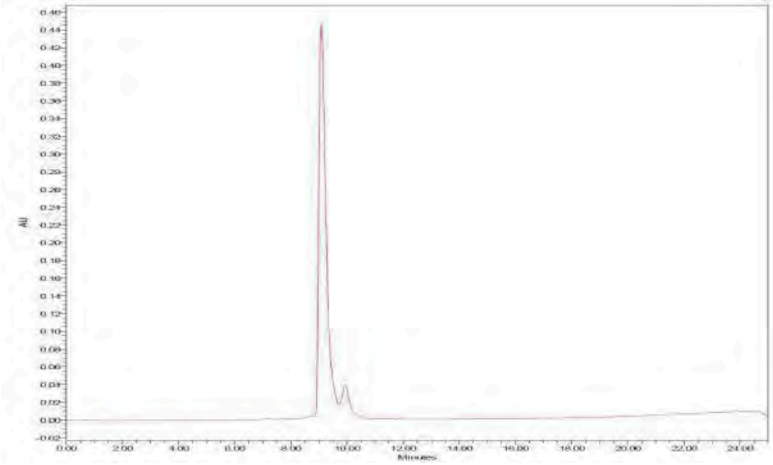

**TJ82 Ac-Ala-Ala-Ile-Glu-Pro-Asp-ACC**

HRMS (m/z) [MH$^+$] calcd. for $C_{39}H_{52}N_8O_{14}$, 856.36, found, 857.32

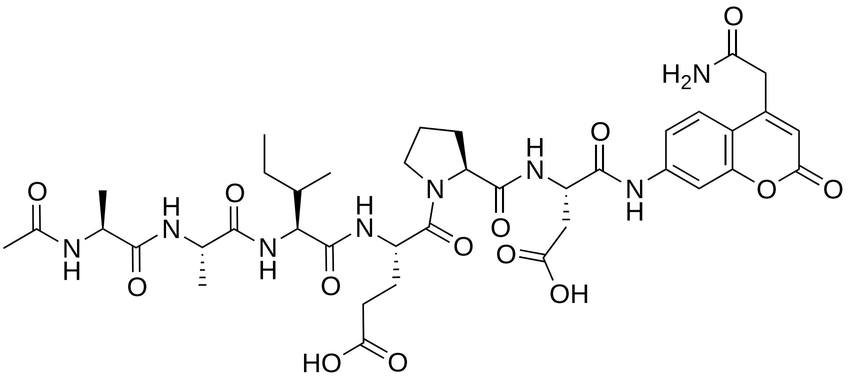

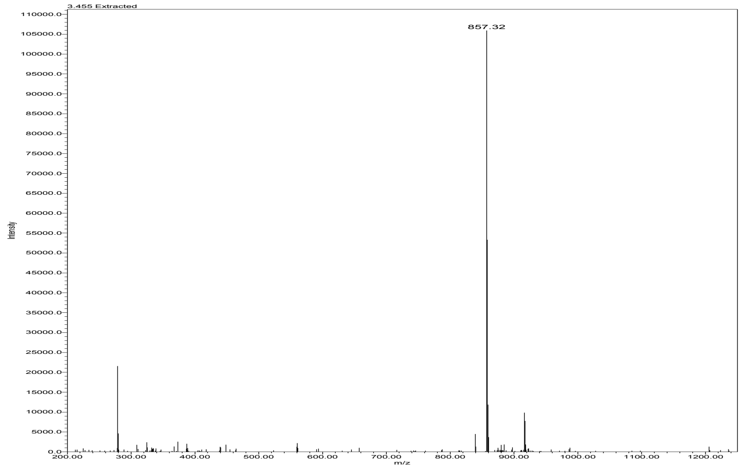


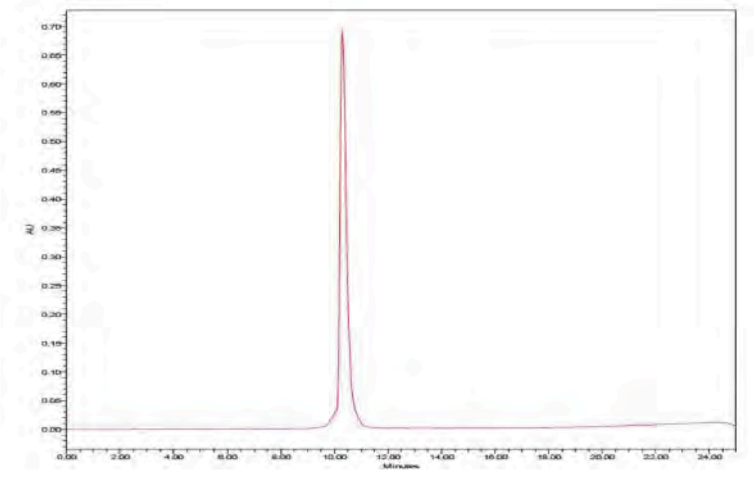
